# Quinone reductase 2 *reads* H3 serotonylation to support human cortical neuron development

**DOI:** 10.64898/2026.03.17.712426

**Authors:** Min Chen, Celi Yang, Xin Li, Lingchun Kong, Benjamin H. Weekley, Xiaoran Wei, Jennifer C O’Chan, Claudia De Sanctis, Victoria Flores Almazan, David A. Vinson, Bulent Cetin, Aarthi Ramakrishnan, Li Shen, Rongsheng Zeng, Zheng Liu, Juner Zhang, Kaylee M. Cappuccio, Joshua R. Sokol, Erdene Baljinnyam, Ruiqi Hu, Kobi Rosenblum, Henrik Molina, Zeng Lin, Qingfei Zheng, Yael David, Marco M. Hefti, Samuele G. Marro, Tom W. Muir, Xiang David Li, Haitao Li, Ian Maze

## Abstract

Histone H3 Gln5 serotonylation (H3Q5ser) is a posttranslational modification that guides transcriptional permissiveness in the brain and periphery. Because it can co-occur with H3K4me3, prior studies focused on its modulation of H3K4me3 *readers*, leaving open whether H3Q5ser acts as an autonomous chromatin mark. Using proteomic, structural, molecular, epigenomic, and cellular approaches, we show that Quinone reductase 2 (QR2), previously known only as a cytosolic enzyme, *reads* H3Q5ser independently of H3K4me3. QR2 recognizes the serotonyl group through its FAD-dependent quinone-binding pocket, discriminating against unmodified H3, H3K4me3, and other monoaminyl marks. In human iPSC-derived neurons, CRISPR-Cas9 disruption of H3 serotonylation or QR2 binding impairs transcriptional programs, synaptic connectivity, and electrophysiological maturation. In cortical organoids, these disruptions cause progenitor over-proliferation at the expense of excitatory neurons and uncouple POU2F1-linked regulatory elements from the excitatory neuron maturation program. Together, these findings reveal an H3 serotonylation-QR2 signaling axis essential for human neurodevelopment.

## INTRODUCTION

Biogenic amines, such as serotonin (5-hydroxytryptamine or 5-HT), are a large class of evolutionarily ancient signaling molecules that contain one or more amino groups. These metabolites are found in diverse tissues in animals and can elicit biological effects locally at the site of production or remotely following transmission through the circulating blood system^1,2^. 5-HT plays critical and broad roles in normal animal physiology, including embryonic development, inflammation, cardiovascular, and gastrointestinal regulation, as well as the stimulation of a broad spectrum of behaviors^3–6^. By extension, aberrant levels of 5-HT, and/or disruption of its signaling pathways, are associated with many diseases, including digestive, cardiac, respiratory, and neuronal disorders^4,6,7^. In brain, 5-HT regulates neural activity via vesicular release from extended axonal projections originating in the dorsal raphe nucleus (DRN), where it acts on multiple G protein-coupled receptor families to regulate behavioral plasticity^3–6^.

In addition to receptor-mediated signaling, 5-HT can be conjugated to Gln residues within certain protein substrates via transamidation by the Ca^2+^-dependent transglutaminase 2 (TG2) enzyme^8,9^. While initially described in the context of cytosolic substrates, we discovered that these monoaminylation reactions additionally occur in the nucleus. For example, 5-HT (as well as dopamine^10–12^ and histamine^13^) can be covalently bound to histone H3 at Gln 5 (H3Q5) to establish H3Q5 serotonylation (H3Q5ser) on nucleosomes^14–16^. Notably, TG2 acts not only as the *writer* of these marks but also as their *eraser* and *exchanger*, resolving a common intermediate either by hydrolysis, which deamidates Gln5, or by transfer to a different monoamine, which substitutes one adduct for another^13^; its activity on chromatin is dictated by steric accessibility rather than by local sequence or pre-existing modifications^16^. H3Q5ser can occur independently of neighboring posttranslational modifications (PTMs); however, this PTM often co-occurs with H3 Lys4 (K4) trimethylation (me3), establishing the dual H3K4me3Q5ser mark at transcriptional start sites (TSSs). The presence of H3K4me3Q5ser serves to potentiate transcription via: 1) increased binding of TAF3, a TFIID subunit, which aids in RNA Pol II-mediated transcription^14,15,17^; and 2) inhibition of H3K4 demethylases (KDM5 family and LSD1)^15^. Moreover, H3Q5ser has been shown to engage WDR5, a member of MLL and SETD1 H3K4 methyltransferase complexes, where it functions to stimulate deposition of H3K4 methylation^13,18^. In addition, the SET-domain containing protein KMT2E is likewise recruited to chromatin by H3K4me3Q5ser during neurodevelopment^19^, and follows the same logic: its PHD finger engages H3K4me3 on its own, and serotonylation of the adjacent glutamine enhances this interaction rather than establishing it.

H3 serotonylation levels have been observed to increase following post-mitotic differentiation of human induced pluripotent stem cells (hiPSCs) to 5-HTergic neurons, as well as in rat neuronal precursor cells from the 5-HTergic DRN (RN46A-B14 cells)^14^. Mutating H3Q5 to alanine (H3Q5A) in RN46A-B14 cells prevents H3 serotonylation, thereby reversing gene expression programs important for neural differentiation. In addition to its documented actions in 5-HTergic cells, the H3K4me3Q5ser mark has been detected *in vivo* in other monoaminergic and non-monoaminergic cells (both neurons and glia) of the brain^14,20^, with its non-monoaminergic distribution hypothesized to be regulated, at least in part, by the ubiquitously expressed high-capacity, low-affinity monoamine transporter OCT3/SLC22A3. In addition to its roles in neural development, H3Q5ser has been found to have important functions in regulating gene expression and cellular outputs in the context of sensory processing^20^, stress- and depression-related phenotypes^21^, placental gene regulation^22^, cancer progression^23–32^, and circadian rhythms^13^.

While H3Q5ser’s functions in the context of neighboring H3K4me3 are now well documented, it remains unclear whether H3Q5ser itself engages with dedicated – non-H3K4me3-associated – *reader* proteins to facilitate its functions in mammalian cells. The effectors characterized thus far do not settle this, because in each instance, appreciable binding occurs in the absence of the serotonyl adduct, which tunes interactions that already exist rather than creating new interactions. Here, employing a series of proteomic-, structural-, molecular-, epigenomic-, and cellular-based assays, we now demonstrate that H3Q5ser is *read* by the Quinone reductase 2 (QR2) enzyme, both *in vitro* and *in vivo*, and that this interaction can occur in the context or absence of adjacent H3K4me3. Quinone reductase 2 (QR2, encoded by *NQO2*) is a FAD-dependent flavoprotein that arose together with its paralog QR1 (NQO1) from an ancient gene duplication^33^. In the presence of a small dihydronicotinamide, QR2 is an efficient catalyst; what it cannot use, however, is NAD(P)H, and the one co-substrate that it can use, dihydronicotinamide riboside, has no demonstrated cellular source, so its canonical chemistry has never been explored in a physiological setting^34,35^. Ancestral-sequence reconstruction indicates that this divergence was directional rather than degenerative, with selection acting against efficient reduction by NAD(P)H along the QR2 lineage, and the same inefficiency is conserved throughout amniotes^35,36^. QR2, nonetheless, remains strongly conserved and retains its flavin-binding scaffold, is the molecular entity long known as the melatonin MT3 binding site^37–39^, and its genetic or pharmacological reduction enhances memory formation in rodents and ameliorates Alzheimer’s disease phenotypes^40–44^. What such a protein – held under selection yet separated from its own canonical chemistry – does in cells has remained unresolved, and QR2 has no described nuclear function. Importantly, implementing CRISPR-Cas9-mediated gene editing to specifically disrupt QR2-H3Q5ser interactions without perturbing QR2’s endogenous enzymatic functions, we show that QR2 binding to H3Q5ser is critical for establishing appropriate gene expression programs in human neurons, which contribute to cellular maturation. In dorsally patterned human cortical organoids, disrupting the same interaction impairs the generation of excitatory neurons from cortical progenitors, placing this requirement within human corticogenesis itself. These findings thus establish QR2 as a *bona fide reader* of H3Q5ser, an interaction that plays essential roles during human neurodevelopment, and one that may contribute to pathological states associated with aberrant 5-HTergic signaling in the brain and beyond.

## RESULTS

### Identification of QR2 as an H3Q5ser binding protein

To initially assess binding interactions between the H3Q5ser modification and nuclear proteins, we performed H3 N-terminal peptide pulldown assays using biotinylated H3_1-18_Q5ser *vs.* H3_1-18_ unmodified, followed by Streptavidin capture and mass spectrometry, to enrich for/identify proteins from HeLa nuclear extracts that display potentiated binding to the serotonyl mark. Following LC-MS/MS analysis, we observed that numerous proteins displayed either modestly attenuated or potentiated binding to H3Q5ser *vs.* H3 unmodified, some of which were consistent with previous studies from our group and others (e.g., attenuated – KDM5A/B^15^; potentiated – WDR5^13,18^). However, one protein – the Quinone reductase 2 enzyme (QR2; aka NQO2 or N-ribosyldihydronicotinamide: quinone dehydrogenase 2) – was found to display a considerably stronger interaction with the H3Q5ser peptide *vs.* other proteins identified in our analysis (**Figure 1A**; **Data S1**); note that QR2’s paralog, QR1, was not found to interact with the H3Q5ser peptide (**Figure 1A**, **inset**), indicating that these two evolutionarily related proteins may have diverged to carry out distinct functions in cells. Next, given previously observed roles for H3Q5ser in guiding gene expression programs within neural cells specifically^14,21^, we developed an orthogonal photoaffinity-based chemical probe approach (probe 1; derived from an H3_1-15_Q5ser peptide in which the Ala7 residue was replaced by a diazirine-containing photoreactive amino acid; **Figure S1A**), to covalently capture H3Q5ser binding proteins upon UV irradiation in SK-N-SH cell lysates (a neuroblastoma cell line). This probe also introduced an alkyne-containing amino acid at its C-terminus to enable bioorthogonal conjugation of fluorescence tags (for visualization) or biotin (for isolation) to target proteins. An unmodified probe C (**Figure S1A**) was developed in tandem as a negative control. To further profile potential binding proteins of H3Q5ser, whole-cell lysates were extracted from SK-N-SH cells and were photo-crosslinked with probe 1 and probe C. Proteins captured by the probes were conjugated to biotin via click chemistry, followed by affinity purification and trypsin digestion. The resulting peptides were labeled with Tandem Mass Tag (TMT) reagents and pooled for subsequent quantitative mass spectrometry analysis (**Figure 1B**). Like our results from HeLa cells, the QR2 enzyme, but not QR1, exclusively stood out with a high TMT ratio (**Figure 1C; Data S2**; validated via immunoblotting in **Figure 1D**). Importantly, QR2’s binding to H3Q5ser was also validated via peptide pulldowns from human brain nuclear extracts (**Figure 1E**), indicating that such interactions are preserved in endogenous brain tissues. We next verified that interactions between QR2 and H3Q5ser are specific, demonstrating that recombinant full-length QR2, but not QR1, can be robustly labeled by serotonylated probe 1, but not the unmodified probe C (**Figure 1F**). We subsequently performed biotinylated peptide pulldown assays, followed by immunoblotting, to compare recombinant QR2’s binding to an unmodified H3 peptide *vs.* those containing H3K4me3, H3Q5ser, or H3K4me3Q5ser (all H3_1-18_). These results indicated that while QR2 does not bind to H3 unmodified or H3K4me3, it does bind to both H3Q5ser and H3K4me3Q5ser peptides with roughly equal affinity (**Figure 1G**, **top**). Given that additional H3 monomainylation marks have also recently been identified, such as H3Q5 dopaminylation (dop)^10–12^ and histaminylation (his)^13^, we next compared QR2 binding to H3 peptides containing H3Q5ser *vs.* H3Q5dop *vs.* H3Q5his *vs.* H3 unmodified, which revealed that QR2 selectively interacts with H3Q5ser (**Figure 1G**, **bottom**). To confirm the strength of QR2’s selective binding to H3Q5ser (in the presence or absence of adjacent H3K4me3), we performed isothermal titration calorimetry (ITC) to determine the K_D_ of these interactions. Consistent with our *in cellulo* and *in vitro* peptide pulldown assays, we observed that QR2 binds to H3Q5ser with moderately high affinity (3 µM), an interaction that was unaffected by the presence of H3K4me3 (H3K4me3Q5ser; 3 µM) (**Figure 1H**; **Table S1**). Importantly, our ITC results similarly indicated that QR2 does not bind H3 unmodified or H3K4me3 peptides (**Figure 1H**), nor does it bind H3Q5dop or H3Q5his PTMs (**Figure 1I**; **Table S1**). Finally, using our orthogonal cross-linking approach, we observed that probe 1-induced labeling could be effectively competed off by H3Q5ser and H3K4me3Q5ser peptides, but not by H3 unmodified, H3K4me3, H3Q5dop, or H3Q5his (**Figure S1B-C**).

**Figure 1.**
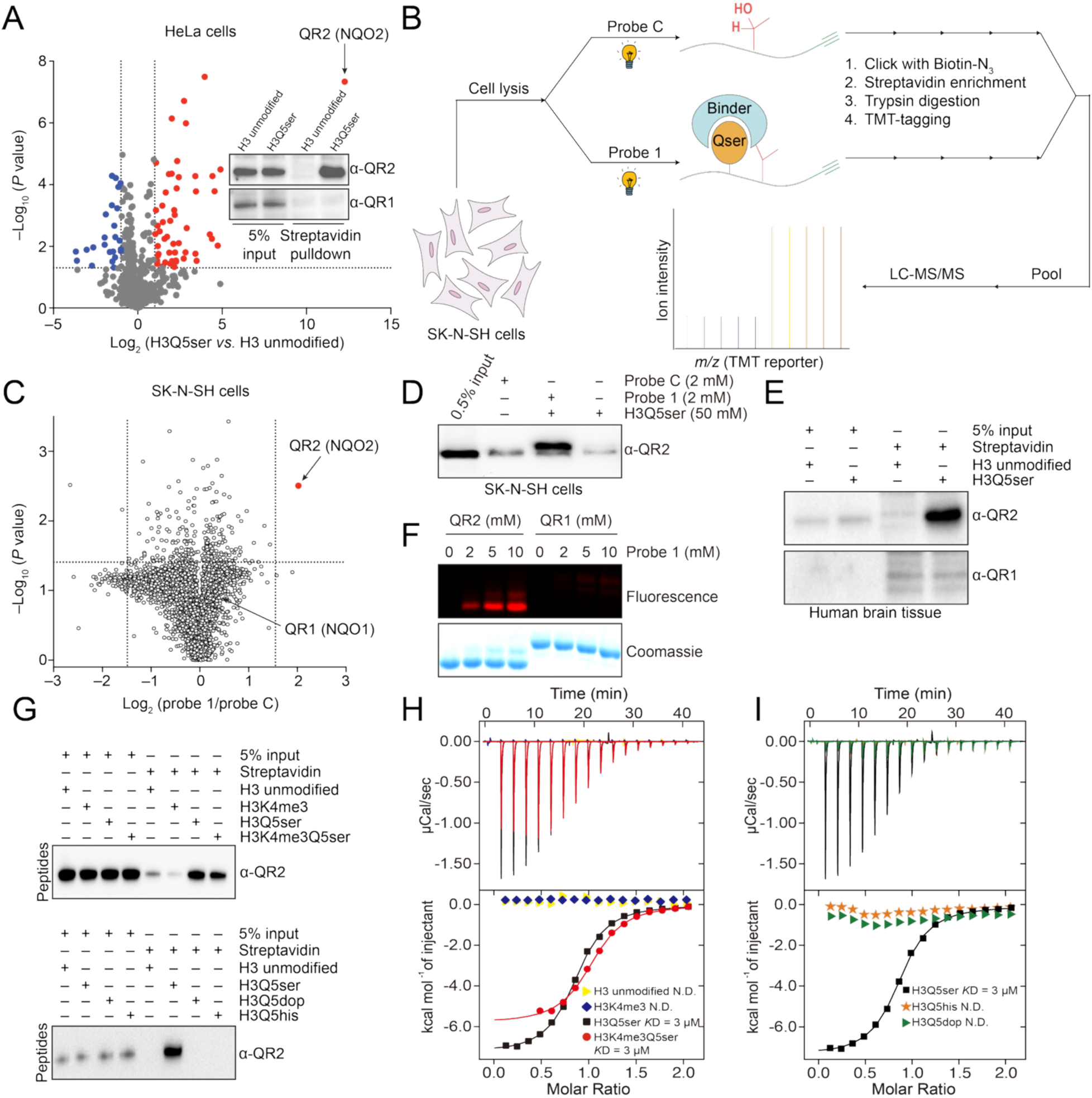
Identification and characterization of QR2 as an H3Q5ser binding proteins. (**A**) Volcano plots of H3_1-18_Q5ser *vs.* H3_1-18_ unmodified binders from HeLa nuclear extracts identified by Streptavidin capture and LC-MS/MS (*n* = 4; –log_10_ *P* value < 0.05; Log_2_ fold difference > 1). Inset: immunoblotting validation of QR2, but not QR1, binding to H3Q5ser in HeLa cells. (**B**) Workflow of TMT-based chemical proteomic profiling of H3Q5ser binding proteins. (**C**) Volcano plot of proteins captured by serotonylated probe 1 *vs.* unmodified probe C from SK-N-SH cell lysates. Statistical analysis was conducted using an unpaired two-tailed Student’s *t*-test. Cutoffs: significance *P* < 0.05; TMT ratio Log_2_ (probe 1/probe C) > 1.5 (*n* = 5). (D) Photo-crosslinking pulldown validation of QR2 binding to H3Q5ser in SK-N-SH cell lysates. (E) H3_1-18_Q5ser *vs.* H3_1-18_ unmodified peptide pulldowns from human brain nuclear extracts, followed by immunoblotting for QR2 or QR1. (**F**) Recombinant QR2, but not its paralog QR1, could be labeled by probe 1. (**G**) Peptide pulldowns with recombinant QR2, followed by immunoblotting for QR2. (**H-I**) Titration and fitting curves of peptides titrated into QR2. K_D_ values are provided. See **Table S1** for ITC statistics. All immunoblotting experiments repeated 3X. See **Figure S14** for uncropped blots.

To date, QR2 has been characterized almost exclusively as a cytosolic flavoprotein that uses FAD as its cofactor to catalyze two-electron reduction of quinones^34^. Its endogenous physiological function, however, remains unresolved: QR2 is inert toward NAD(P)H, instead depending on the rare co-substrate NRH, and comparative evolutionary analysis of the two quinone reductases has led to the proposal that QR2 drifted toward pseudoenzyme function^36^ – consistent with our observation that QR1, which retains canonical reductase activity, does not bind H3Q5ser (**Figure 1A,C**). Of relevance here, QR2 is an indoleamine-binding protein: melatonin and structurally related indoleamines, including 5-HT, occupy the FAD-proximal substrate pocket of the QR2 homodimer and inhibit its reductase activity^37,38^. QR2 thus harbors a structurally defined pocket capable of accommodating the same indoleamine moiety that is transamidated onto H3Q5 during serotonylation. Yet, although QR2 is expressed across subcellular compartments, no nuclear function has been ascribed to it. Genetic variants of *NQO2* have been identified in humans, linking dysregulation of its function to tumorigenesis and neurodegenerative disorders^45^. Additionally, studies in *Nqo2* deficient mice have indicated roles for this protein in the regulation of cognitive behaviors^46^. While our data indicate that QR2 binds to H3Q5ser in neural cells, including in human brain tissues, whether such interactions contribute to previously observed neural phenotypes remains unknown.

### Molecular recognition of H3Q5ser by QR2

Given that QR2 binds to H3Q5ser directly, we next wished to elucidate the way this molecular recognition occurs. To do so, we performed x-ray crystallography to capture the QR2-H3Q5ser complex containing QR2’s coenzyme FAD. These analyses demonstrated that H3Q5ser inserts within a binding pocket near the homodimerization interface of QR2’s chain A and chain B (**Figure 2A-B**; **Table S2**), with rotation of the complex revealing full electron density tracing of the H3Q5ser peptide from H3 Arg (R) 2 to H3 Ala (A) 7 (**Figure 2C**; **Figure S2A**). Further zooming into this binding pocket, we were able to determine that Tyr (Y) 104 of QR2’s chain A, the indole ring of 5-HT, and FAD form π-π stacking interactions, while QR2’s chain A Trp (W) 105 and Phe (F) 106, along with QR2’s chain B F126 and F178, collectively form a hydrophobic aromatic cage to facilitate the specific binding pocket for H3Q5ser recognition (**Figure 2D**). This binding requires the formation of critical contacts between the H3Q5ser peptide and QR2, with the main chain of the H3Q5ser peptide being surrounded by multiple hydrophobic interactions and several pairs of hydrogen bonds (**Figure S2B**). Of note, the *apo* form of QR2 lacking the co-factor FAD was observed to completely lose its H3Q5ser binding activity (**Figure S2C**), highlighting the FAD-dependent nature of H3Q5ser recognition by QR2. Importantly, given that QR2 can bind to H3Q5ser in both the absence and presence of H3K4me3, we next performed x-ray crystallography to capture the QR2-H3K4me3Q5ser complex, again containing QR2’s coenzyme FAD. These data demonstrated that QR2’s molecular recognition of the dual PTM follows a similar logic to its binding of H3Q5ser alone, with H3K4me3 projecting outward from the QR2-H3Q5ser binding pocket (**Figure S2D-G**; **Table S2**); such findings provide further support to our previous observations that the presence of H3K4me3 in the context of H3Q5ser does not impact the strength of QR2’s interaction with the serotonyl mark.

**Figure 2.**
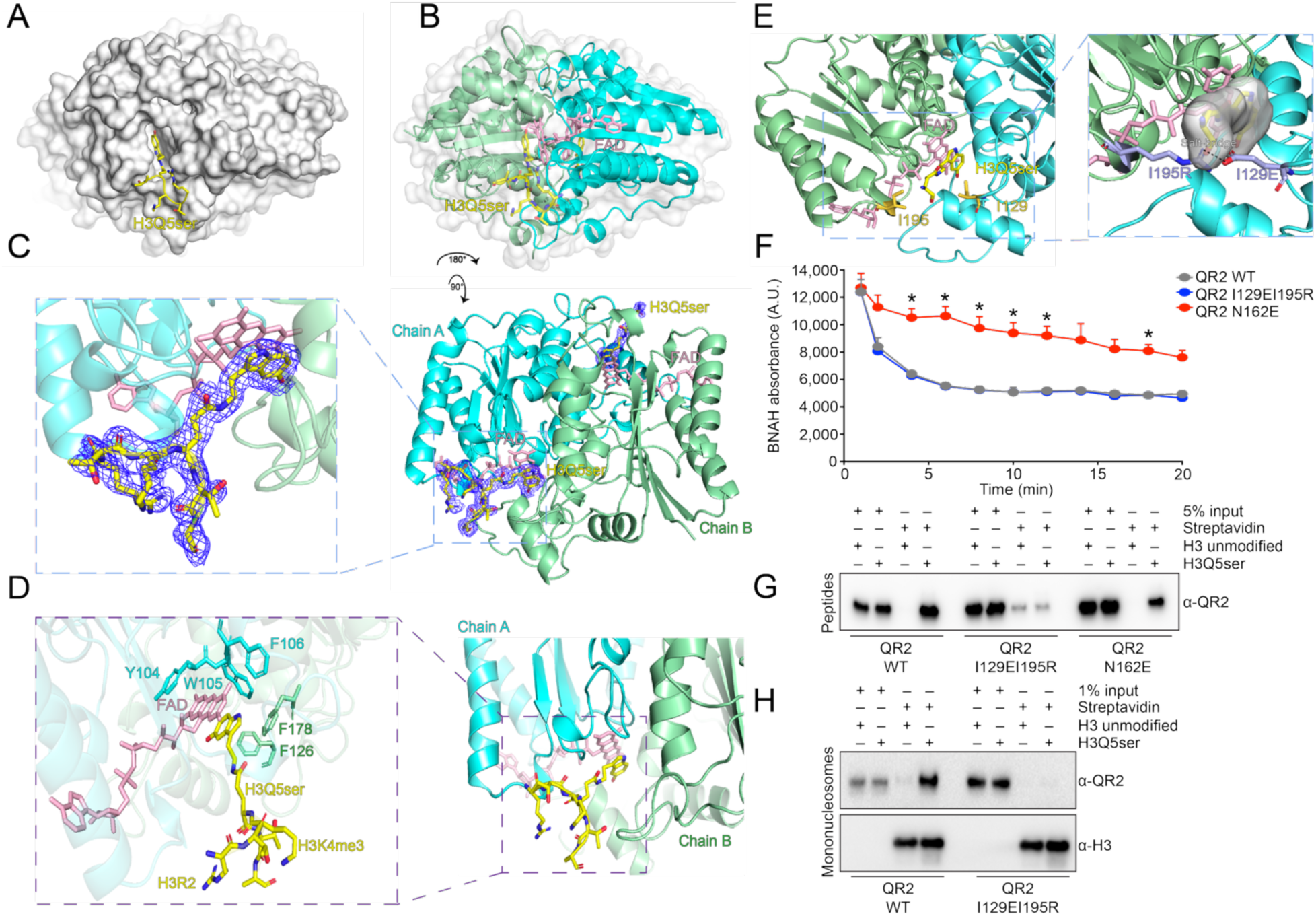
Molecular recognition of H3 Gln5 serotonylation by QR2. (**A**) Structure of the QR2-H3Q5ser complex. H3 residues are shown as yellow sticks, QR2 is shown as a white surface. The H3Q5ser residue inserts into the binding pocket of QR2. (**B**) Ribbon structure of the QR2-H3Q5ser complex. Ribbon shows detailed construction of QR2, H3 residues are shown as yellow sticks, coenzyme FAD is shown as pink sticks. (**C**) After rotating by a certain angle, as indicated in the diagram, the electron density of the H3Q5ser peptide is displayed, shown as a blue mesh. Chain A and Chain B in the QR2 dimer are respectively labeled. (**D**) Details of the QR2-H3Q5ser complex structure are shown. H3 residues are shown as yellow sticks, coenzyme FAD is shown as pink sticks. The key amino acid residues in Chain A (cyan) and Chain B (green) are labeled. Chain A Y104, FAD, and the serotonin ring form π-π stacking interactions, while Chain A W105, F106, Chain B F126, and F178 collectively form a hydrophobic aromatic cage, constituting the specific pocket for recognizing H3Q5ser. See **Table S2** for x-ray crystallography data collection and refinement statistics. (**E**) Molecular design of mutants to disrupt binding to H3Q5ser. The key amino acid residues I129 and I195 are labeled in orange. The mutants I129E and I195R are displayed as purple sticks. I129E and I195R are predicted to form a salt bridge via structural modeling, preventing the H3Q5ser peptide from inserting into the internal pocket of QR2. (**F**) BNAH enzymatic assay comparing wildtype *vs.* I129EI129R (H3Q5ser binding deficient) *vs.* N162E (catalytically deficient) QR2. *n* = 3. Two-way RM ANOVA [main effects: time (F_1.908, 11.45_ = 107.2, *P*<0.001), QR2 mutation (F_2,6_ = 21.20, *P*=0.0019), and QR2 mutation x time (F_3.816, 11.45_ = 4.884, *P*=0.0161); post hoc: Dunnett’s MC test (*p<0.05; QR2 N162E *vs.* wildtype)]. Data shown as mean ± SEM. (**G**) Peptide pulldown validations of mutant *vs.* WT QR2 abrogation of binding on H3Q5ser. (**H**) Mononucleosome pulldown validations of mutant *vs.* WT QR2 abrogation of binding on H3Q5ser. All immunoblotting experiments repeated 3X. See **Figure S14** for uncropped blots.

To provide orthogonal evidence for how QR2 interacts with the H3Q5ser peptide, we additionally mapped peptide binding sites on the protein surface using our photo-crosslinking strategy^47^ (**Figure S3A-C**), which revealed major crosslinks at residues located around the Q5ser-binding pocket of QR2 when mapped back to our co-crystal structure (see **Figure S4** and **Table S1** for further ITC validations of QR2’s molecular recognition of H3Q5ser). Notably, probes bearing the photo-crosslinker N-terminal (T3; probe 2) *vs.* C-terminal (A7; probe 3) to Q5ser converged on an overlapping set of contact residues (E153, E199), while each additionally captured distinct surfaces (**Figures S3B-C**), consistent with the defined peptide orientation across the QR2 dimer interface observed crystallographically. Given that such photo-crosslinking was performed in solution, where both the protein and peptide exist in a freely moving state, these binding site mapping results complement and further support our crystallographic study, indicating that the interaction surface captured in the crystal reflects the predominant mode of QR2–H3Q5ser engagement in solution rather than an artifact of crystal packing.

Finally, to determine whether unique interactions observed in the QR2-H3Q5ser binding pocket are indeed critical for QR2’s molecular recognition of the mark, we focused our attention on two specific residues that contribute to hydrophobic interactions when QR2 is bound to H3Q5ser, Ile (I) 129 and I195. To disrupt QR2 binding to H3Q5ser, we next generated mutations in QR2 (I129EI195R) that were predicted – via structural modeling – to abrogate binding through the introduction of a salt bridge (**Figure 2E**). These mutations were found to have no impact on QR2’s endogenous enzymatic functions (*vs.* QR2 Asn (N) 162 to E); **Figure 2F**), while fully attenuating QR2’s binding to H3Q5ser, both in the context of peptides (**Figure 2G**; **Figure S5A and S5C**; **Table S1**) and mononucleosomes (**Figure 2H**; **Figure 5D**). QR1 showed no selective binding to H3Q5ser-containing mononucleosomes (**Figure S5D**), extending to chromatin the paralog specificity observed with peptides. Given that QR2 has also been reported to bind free 5-HT, we additionally performed ITC assessments to explore the impact of QR2 I129EI915R mutations on its binding to free 5-HT, where we observed negligible effects (**Figure 5B**). Notably, wildtype QR2 bound the H3Q5ser peptide and free 5-HT with indistinguishable enthalpies (−11.16 ± 0.07 *vs.* −11.18 ± 0.24 kcal mol⁻¹) but different binding entropies (−8.41 *vs.* −13.5 cal mol⁻¹ deg⁻¹; **Table S1**), indicating that the ∼13-fold higher affinity for the peptide (452 nM *vs.* 5.78 µM) is entropic in origin rather than reflecting additional enthalpic contacts. This is consistent with the serotonin moiety supplying the bulk of the binding energy through the aromatic cage resolved in our co-crystal structure (**Figure 2D**), with the histone backbone contributing affinity principally by reducing the entropic cost of binding, as expected if it buries additional apolar surfaces.

The residues mediating *reading* and catalysis line the same pocket but are separable (**Figure 2F-2G**), so we asked how each has fared over evolution. Across 16 QR2 orthologues spanning lamprey to human, pairwise identity ranges from 54% to 99%, and 32% of positions (75 of 231) are invariant, arguing against pseudogene-like decay (**Figure S5E-F; Data S3**). Among the three positions we manipulated, Ile129 is the most constrained: it is identical to human in 11 of 16 species and hydrophobic or aromatic in all sixteen, consistent with a packing requirement at the face that contacts the serotonyl moiety. Position 195 is an invariant isoleucine across the primates but varies widely among other vertebrates, and Asn162 alternates with serine throughout the tree, including within primates (**Figure S5G**). Because strong conservation and an intact flavin scaffold are expected of any flavoprotein, and because *reading* and catalysis depend on residues lining a single pocket, sequence conservation alone cannot specify which activity is under selection; that separation rests instead on our binding- and catalysis-dead mutants.

### QR2 binding to H3Q5ser contributes to cellular gene expression across species and cell-types

Since our structural analyses indicated that specific hydrophobic interactions within the QR2-H3Q5ser binding pocket can be perturbed to prevent QR2-H3Q5ser recognition without impacting other aspects of QR2’s functions, we next aimed to exploit these QR2 I129EI195R mutations in cells to explore potential roles for QR2-H3Q5ser binding in the regulation of mammalian gene expression. To do so, we first examined whether QR2 co-enriches with H3(K4me3)Q5ser genome-wide in HeLa cells, a simple genetically modifiable tissue culture system, which has previously been shown to contain the H3 serotonyl mark owing to the presence of 5-HT supplemented to the media via serum^16^. We performed CUT&RUN-seq^48^ using antibodies recognizing H3K4me3Q5ser (validated previously^14^), QR2, and RNA Pol II, which demonstrated that QR2 co-enriches with H3K4me3Q5ser genome-wide, largely at permissive genes displaying co-enrichment with RNA Pol II (**Figure S6A-B**; **Data S4**). Given such co-enrichment, we next performed CRISPR-Cas9 editing to knockdown endogenous QR2 in HeLa cells, followed by genetic complementation (under the endogenous *NQO2* promoter) with either FLAG-HA-tagged QR2 wildtype or QR2 I129EI195R (**Figure S6C**); note that complementation with the mutant form of QR2 did not have any impact on QR2’s subcellular localization, which is both cytoplasmic and nuclear **(Figure S6D**). Importantly, employing H3_1-18_Q5ser *vs.* H3_1-18_ unmodified peptide pulldowns in HeLa nuclear extracts from wildtype *vs.* mutant complemented cells, we first verified that our approach worked, demonstrating that biotinylated H3Q5ser peptides efficiently pulldown wildtype, but not mutant, QR2 in knockdown/complemented cells (**Figure S6E**). Next, we performed CUT&RUN-seq for H3K4me3Q5ser, QR2, and RNA Pol II in wildtype *vs.* QR2 knockdown (no complementation) *vs.* FLAG-HA-QR2 *vs.* FLAG-HA-QR2 I129EI195R complemented cells, demonstrating that QR2 knockdown results in reduced co-enrichment of QR2 at H3K4me3Q5ser/RNA Pol II positive loci, an effect that was fully rescued with wildtype, but not mutant, QR2 complementation (**Figure S6F-H**; **Data S5**). Finally, to assess whether QR2’s binding to H3Q5ser plays functional roles in guiding gene expression in HeLa cells, we performed RNA-seq in wildtype *vs.* QR2 knockdown (no complementation) *vs.* QR2 (wildtype or mutant) complementation cells, which demonstrated that the vast majority of genes displaying differential expression between QR2 knockdown *vs.* wildtype cells were efficiently rescued by complementation with wildtype, but not mutant, QR2 (**Figure S6I-L**, **Data S6**).

To ensure that QR2-H3Q5ser interactions are indeed conserved across species and/or cell-types, we next assessed QR2’s subcellular localization in mouse brain (**Figure S7A**) and primary neurons (mouse cerebellar granule neurons (cGNs), cultured in serum to provide a 5-HT source; **Figure S7B**), where we found that – as in HeLa cells – QR2 is distributed throughout both cytoplasmic and nuclear compartments. ChIP-seq for H3(K4me3)Q5ser, QR2, and RNA Pol II in cGNs further revealed that QR2 largely co-localizes with H3(K4me3)Q5ser genome-wide at permissive genes co-enriched by RNA Pol II (**Figure S7C-E**; **Data S7**). We additionally found that loci determined to be activity-dependent in response to KCl-mediated neuronal depolarization displayed concordant regulation (primarily induction) of H3Q5ser, QR2, and RNA Pol II enrichment (**Figure S7F-H; Data S7**). To further assess whether activity-dependent regulation of H3Q5ser-QR2 co-enrichment plays causal roles in the regulation of depolarization-induced gene expression (focused here on immediate early genes (IEGs)), we performed two complementary sets of analyses. (1) following transduction of cGNs with lentiviruses expressing either FLAG-HA-tagged H3.3 WT or H3.3 Q5A ± KCl (note that since neurons can only actively incorporate replication-independent H3.3 into postmitotic chromatin^49^, expression of H3.3Q5A effectively functions as a dominant negative to reduce levels of H3Q5 monoaminylations^10,13,14,21^), we performed chromatin immunoprecipitations (ChIPs) to enrich for exogenously expressed H3.3 (wildtype or mutant), followed by re-ChIPs for either H3Q5ser or RNA Pol II. In doing so, we found that while depolarization resulted in significantly increased enrichment of both H3Q5ser and RNA Pol II at IEG promoters in H3.3 wildtype-expressing cells, such enrichment was lost following transduction with the H3.3Q5A mutant (**Figure S7I-J**); these reductions in H3Q5ser and RNA Pol II in the mutant expressing cells similarly corresponded to a loss of IEG expression (**Figure S7K**). (2) To examine whether disrupting QR2 binding to H3Q5ser similarly results in a loss of KCl-mediated IEG induction, we employed recently described QR2 inhibitors, yb800 and yb537, which have been shown to bind QR2 within the QR2-H3Q5ser binding pocket near QR2 I129^40^. Both QR2i inhibitors were found to reduce QR2 binding to H3Q5ser *in vitro* (**Fig S7L**), and yb800 was found to significantly reduce the induction of IEGs in cGNs following depolarization (**Fig S7M**).

### H3Q5ser and QR2 track human excitatory neuron maturation and reproduce the fetal cortical landscape

QR2 binding to H3Q5ser shapes gene expression in HeLa cells and cGNs, including activity-dependent transcription, but whether these interactions guide human neuronal maturation was untested. We had shown previously that H3Q5ser is robustly induced in response to human 5-HTergic neuronal differentiation, and that disrupting H3Q5ser levels in immortalized 5-HTergic rat cells results in differentiation deficts^14^; however, we had yet to conclusively demonstrate whether the serotonyl mark, or its *reader* interactions, causally contribute to human neurodevelopmental processes. As such, and given that H3Q5ser has been found to be widely distributed across neuronal cell-types in brain^14^, we turned to a rapid single-step human neuronal induction system in which hiPSCs expressing the transcription factor Neurogenin-2 (NGN2) are efficiently differentiated into excitatory neurons^50^ (**Figure S8A**). Co-culture with mouse astrocytes drove robust maturation programs over the course of only 35 days *in vitro* (DIV): transcriptomes separated by timepoint along PC1 (66.4% of variance; **Figure S8B**), differentially expressed genes resolved into sequential waves (**Figure S8C**), and canonical markers moved from progenitor through intermediate to mature-neuron identity (**Figure S8D; Data S8**). Cultures were maintained in 5-HT-containing serum, which sustains H3 serotonylation in this system. To place these neurons on a developmental scale, we applied a transcriptomic staging procedure based on the BrainSpan developmental transcriptome^51^. Benchmarking it against human fetal cortical samples of known gestational age recovered clinical age in the early specimens (Spearman r = 0.84; mean absolute error 0.8 weeks; **Figure S8E; Data S10; Table S3**). Applying the identical, validated procedure to the NGN2 time course placed DIV12 through DIV35 at approximately 16 to 19 post-conceptional weeks, advancing monotonically with differentiation day, with DIV0 and DIV6 excluded as pre-neural (**Figure S8F**). Both the mark and its *reader* were present in these neurons and increased with maturation. H3K4me3Q5ser and QR2 were each detected by immunofluorescence in MAP2-positive DIV35 neurons (**Figure S8G**). CUT&RUN-seq profiling across the same window showed that H3K4me3Q5ser enrichment rose from DIV0 to DIV35, and that QR2 – nearly absent at DIV0 – co-enriched at those same loci by DIV35 when signal was anchored on H3K4me3Q5ser peaks (**Figure S8H; Data S9**). H3K4me3Q5ser and QR2 were additionally co-detected by chromogenic immunohistochemistry (IHC) in human fetal cortex (**Figure S8I**), establishing that the two are present together in the primary tissue that these cultures model.

We next asked whether NGN2-induced neurons recapitulate the H3K4me3Q5ser landscape of the developing human brain, using fetal cortex H3K4me3Q5ser peaks as a common reference set. Although the two datasets were generated independently and differed markedly in absolute signal amplitude, independently normalized enrichment profiles were similarly distributed around fetal peak centers (**Figure S8J**). The correspondence held without reference to signal magnitude: across 32,453 fetal loci, peak-level enrichment ranks correlated strongly between fetal cortex and wild-type NGN2 neurons (Spearman ρ = 0.87) and 86.3% of loci in the strongest fetal quartile also fell in the strongest NGN2 quartile – 3.45-fold over the 25% expected by chance (**Figure S8K; Data S11**). NGN2-induced neurons therefore reproduce the relative genomic distribution of fetal H3K4me3Q5ser despite differences in absolute intensity. Distribution alone, however, does not show that the mark tracks transcription, so we asked, in fetal cortex itself, whether the promoters gaining H3K4me3Q5ser across fetal cortical maturation were the same ones gaining transcriptional output. All 11,164 genes carrying a promoter H3K4me3Q5ser peak were ranked twice – by the direction and magnitude of their late-versus-early change in promoter occupancy (late, gestational weeks 37 and 41; early, weeks 18–23; **Table S3**), and by the direction and significance of their change in expression – and the two rankings were compared by RRHO; they agreed strongly (**Figure S8L; Data S10 and S11**). Overlap peaked at the most-gained, most-increased ends of both lists, where 1,351 of the top 2,756 H3K4me3Q5ser genes, and the top 2,438 upregulated genes, were shared, against 602 expected by chance (2.24-fold; p < 5 × 10⁻⁴) – importantly, they remained elevated well beyond this point rather than being confined to a small set of extreme genes. Because the late group comprised two specimens, the expression contrast was additionally fitted with gestational week as a continuous covariate, which returned a concordant ranking.

Cell-type enrichment of the 1,351 concordant genes, tested against all promoter-marked expressed genes, resolved them as predominantly glial (**Figure S8M; Data S12**). Microglial references – including brain macrophages, which share the microglial marker set – and astrocytic references were the strongest terms: microglia of embryonic prefrontal cortex ranked first (140 of 476 genes; 2.43-fold; adjusted p = 1.4 × 10⁻³⁷), with the remaining glial terms at 1.6–2.2-fold (adjusted p ≤ 4.1 × 10⁻⁸). A brain neuronal reference was also enriched, though weakly and close to the significance threshold (27 of 172 genes; 1.3-fold; adjusted p = 0.034). This ordering follows from what the contrast measures rather than from the mark itself. Microglia and astrocytes enter and expand in cortex across the second half of gestation, whereas neurons are already present at earlier timepoints, so genes whose promoters gain H3K4me3Q5ser in step with rising transcription in bulk tissue are preferentially glial. Because H3K4me3Q5ser increased at ∼99% of promoter peaks genome-wide, it is the concordance with expression, not the gain of the mark, that resolves this identity. These data derive from bulk tissue and therefore reflect the emergence and expansion of those lineages across gestation; establishing that H3K4me3Q5ser regulates these genes within a single cell-type would require cell-type-resolved profiling. The neuronal arm is addressed below, where the same analysis in a purely neuronal population returns synaptic and axon-guidance programs.

Finally, we asked whether the coupling between gain of mark and gain of transcription similarly operates in the NGN2 system. Ranking genes by their DIV35-versus-DIV0 change in promoter H3K4me3Q5ser and by their change in expression showed concordant enrichment at the gained, upregulated ends of both lists (**Figure S8N**), and genes gaining both were enriched for chemical synaptic transmission, nervous system development, and axon guidance (**Figure S8O; Data S13**). NGN2-induced neurons therefore faithfully reproduce not only the genomic distribution of fetal H3K4me3Q5ser but its coupling to transcriptional output, giving us a tractable system in which to disrupt that coupling genetically.

Using CRISPR-Cas9-mediated gene editing in the parental hiPSC line WTC11^52^, we generated multiple homozygous knock-in clones targeting the *NQO2* (QR2 I129EI195R; disrupting H3Q5ser binding while preserving enzymatic functions), *TGM2* (C277A; catalytically inactivating its monoaminylase functions), and *H3F3B* (Q5A and Q5N; rendering ∼50% of H3 in post-mitotic neurons incapable of serotonylation). These lines enabled us to directly assess roles for H3Q5ser and QR2-H3Q5ser interactions during human neuronal development. Note that while two genes – *H3F3A* and *H3F3B* – encode for the H3.3 variant in humans, we selectively targeted *H3F3B* to achieve partial disruption of serotonylation while preserving neuronal viability^53^. Correct targeting was confirmed by Sanger sequencing, and genomic integrity was verified by copy-number profiling (**Figure S9A-F**). In differentiated neurons at DIV35, H3.3 Q5A and Q5N mutations resulted in significantly reduced levels of H3K4me3Q5ser relative to wildtype isogenic controls, with similar attenuation observed in QR2 I129EI195R mutant neurons (**Figure S9G**), suggesting that QR2 binding to H3Q5ser may help to stabilize the mark against TG2-mediated erasure. Notably, the QR2 mutation did not alter overall protein expression levels in differentiated neurons (**Figure S9H**). Peptide pulldown assays using H3_1-18_Q5ser *vs.* H3_1-18_ unmodified peptides further demonstrated robust binding of wildtype, but not mutant, QR2 to the H3 serotonylation mark (**Figure S9I**). QR2 present at normal levels but unable to bind therefore still lowers H3K4me3Q5ser, implicating reader occupancy in protecting the mark from TGM2-mediated erasure, which we confirmed directly (**Figure S10A-B**).

To anchor these analyses in primary tissue, we first profiled H3K4me3Q5ser and QR2 by CUT&RUN-seq in human fetal cortex, where the two co-enriched genome-wide, including at neuronal genes such as SYT1 (**Figure 3A; Data S11; Table S3**). Employing CUT&RUN-seq, we then profiled enrichment patterns for H3K4me3Q5ser, QR2, and RNA Pol II across wildtype *vs.* mutant NGN2-induced neurons at DIV35. Similar to that observed in HeLa cells and mouse cGNs, we found that QR2 displays robust co-enrichment with H3K4me3Q5ser at permissive loci (co-enriched for RNA Pol II) in wildtype human neurons (**Figure 3B and 3G**; **Data S14**); however, this co-enrichment was severely attenuated across all four mutant lines (**Figure 3C-G**), indicating that both direct (Q5A and Q5N) and indirect manipulations of the mark or its binding interactions with QR2 (QR2 I129EI195R and TG2 C277A) are sufficient to disrupt H3Q5ser-QR2 co-enrichment genome-wide. In addition, consistent with our findings in mouse cGNs, we observed that such perturbations are also sufficient to reduce RNA Pol II localization at H3K4me3Q5ser-marked loci. Corresponding RNA-seq analyses placed these chromatin changes against the developmental program they serve. The transcriptional changes produced by QR2 I129EI195R were discordant with those separating late from early human fetal cortex, indicating that disrupting recognition of the mark opposes the direction of the endogenous developmental program (**Figure 3H**). The four mutant lines nonetheless converged on one another: pairwise RRHO comparisons showed strong concordance amongst them, whether the mark itself, its *writer* or its *reader* had been targeted (**Figure 3I-L**; **Figure S9J-K**; **Data S8 and S15**). Indeed, differential expression analysis identified 1,193 protein-coding genes displaying overlapping and consistent dysregulation across the four mutant lines (**Figure 3M-O; Data S16**), with these overlapping genes found to be enriched for gene ontologies related to neuronal maturation (e.g., axonogenesis, nervous system development, synapse organization, and so on; **Figure 3P**; **Data S17**). Finally, since both disruption of H3Q5ser levels (H3.3 Q5A *vs.* WT transduction) and inhibition of QR2 binding to H3Q5ser (yb800 *vs.* vehicle) were found to disrupt activity-dependent gene expression in mouse cGNs, we next assessed the potential impact of perturbing H3Q5ser (Q5N) or QR2 binding to H3Q5ser (QR2 I129EI195R) in differentiated human neurons in response to KCl-mediated depolarization. Consistent with our studies in mouse cGNs, we observed that both manipulations were sufficient to attenuate activity-dependent gene regulation in NGN-induced neurons (**Figure S9L**).

**Figure 3.**
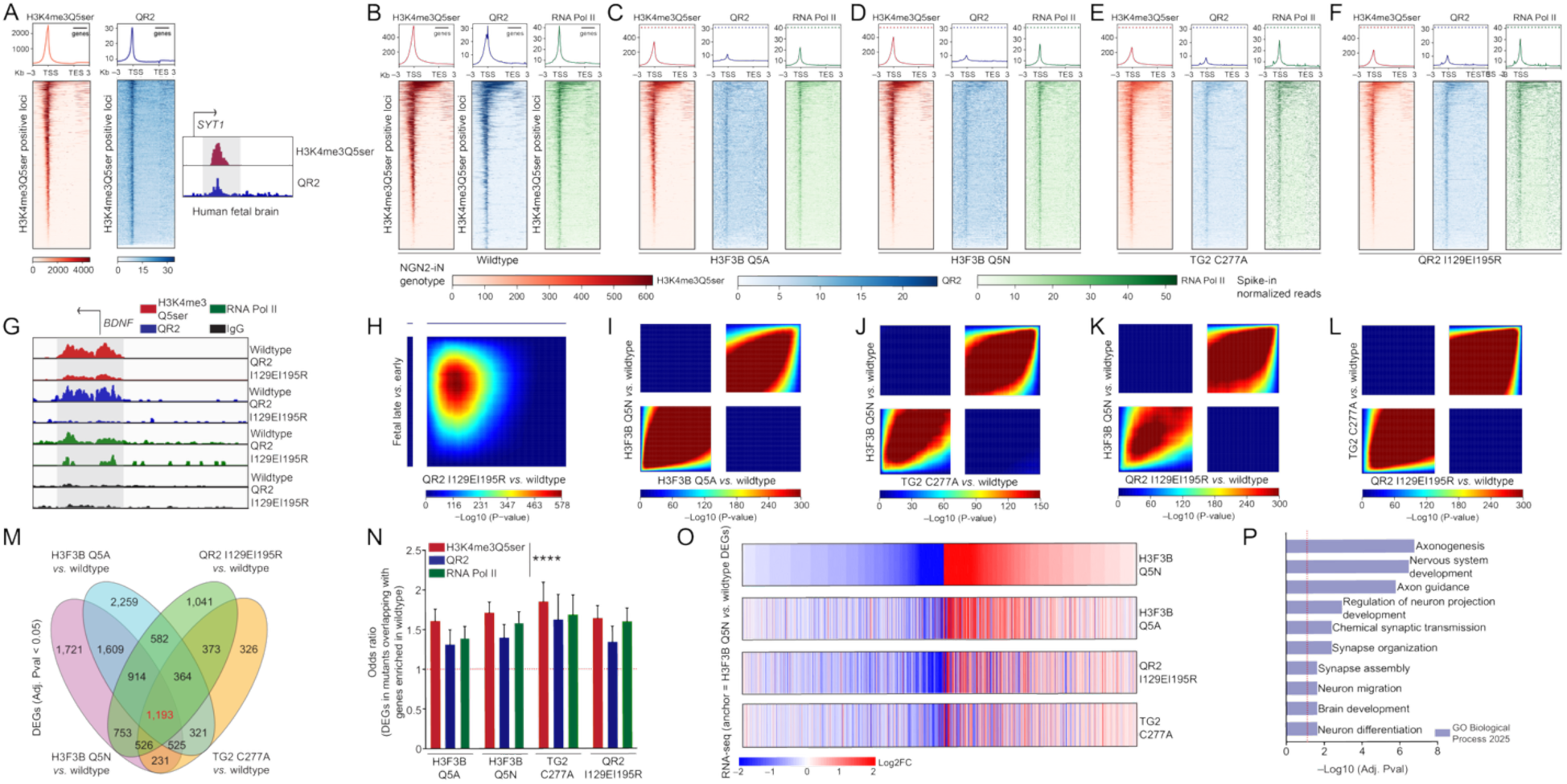
QR2-H3Q5ser interactions contribute to human neurodevelopmental gene expression. (**A**) QR2 co-localizes with H3K4me3Q5ser genome-wide in human fetal cortex. Average metagene profiles (top) and heatmaps (bottom) of H3K4me3Q5ser (red) and QR2 (blue) CUT&RUN signal across H3K4me3Q5ser-positive genes in human fetal cortex, scaled from transcription start site (TSS) to transcription end site (TES) with ±3 kb flanks and ordered by decreasing H3K4me3Q5ser signal, with the same gene order applied to both (*n* = 8; peak enrichment (spike-in normalized read coverage) anchored on H3K4me3Q5ser signal). Bottom, representative aggregated CUT&RUN tracks at the neuronal gene *SYT1* showing co-enrichment of H3K4me3Q5ser and QR2 over the promoter (shaded); arrow, TSS and direction of transcription. CUT&RUN-seq heatmaps of H3K4me3Q5ser, QR2, and RNA Pol II enrichment in (**B**) wildtype, (**C**) H3F3BQ5A, (**D**) H3F3BQ5N, (**E**) TG2 C277A, and (**F**) QR2 I129RI195R hiPSC neurons at DIV 35 (*n* = 3 for all; peak enrichment (spike-in normalized read coverage) anchored on wildtype signal for each line). For B-F, dashed lines represent wildtype signal. (**G**) IGV tracks for H3K4me3Q5ser, QR2, and RNA Pol II at the *BDNF* locus in wildtype *vs.* QR2 I129EI195R hiPSC neurons. For each antibody, scales are consistent between wildtype and QR2 mutants. **(H)** The QR2-I129EI195R mutation opposes the normal human cortical maturation program. RRHO2 between the transcriptional effect of the QR2-I129EI195R mutation in NGN2 induced neurons (mutant *vs.* wildtype; x-axis) and normal human fetal cortical maturation (Fetal Late *vs.* Early; y-axis). Genes were ranked independently in each contrast, the single strong overlap hotspot (peak −log₁₀P ≈ 578) indicates that the QR2-mutant transcriptome is discordant with normal maturation — genes normally increased across fetal Late-versus-Early development are preferentially reduced in QR2-I129EI195R neurons — showing that disrupting the QR2–H3Q5ser interaction derails the normal human cortical maturation program. RRHO analyses comparing gene expression patterns (RNA-seq) between (**I**) H3F3B Q5A *vs.* wildtype and H3F3B Q5N *vs.* wildtype, (**J**) H3F3B Q5N *vs.* wildtype and TG2 C277A *vs.* wildtype, (**K**) H3F3B Q5N *vs.* wildtype and QR2 I129EI195R *vs.* wildtype, and (**L**) TG2 C277A *vs.* wildtype and QR2 I129EI195R *vs.* wildtype demonstrating concordance across mutant lines (*n* = 3 for all). (**M**) Venn diagram displaying the number of overlapping differentially expressed genes (DEGs) between wildtype and mutant hiPSC neurons (FDR < 0.05). (**N**) Bar plot of odds ratio analysis (Fisher’s exact text) examining the overlap of DEGs across mutant lines *vs.* wildtype and enrichment of H3K4me3Q5ser, QR2, and RNA Pol II in wildtype neurons at these same genes (\*\*\*\**P*<0.0001). (**O**) Heatmaps of genes identified as being differentially expressed in H3F3B Q5N *vs.* wildtype neurons (anchor; FDR < 0.05) across all mutant lines. (**P**) Go Biological Process 2025 enrichment for the 1,193 overlapping DEGs identified in panel **M** (FDR<0.05).

### QR2-H3Q5ser interactions are required for human neuronal physiological maturation

Given our observations that disruption of either H3Q5ser itself or its interactions with QR2 produces similar alterations in gene expression programs associated with human neuronal maturation, we next investigated the functional and physiological consequences of these perturbations in mature NGN2-induced neurons. To determine whether QR2-H3Q5ser disruption affects structural maturation, we quantified neuronal morphology at DIV35 using sparse GFP labeling of wildtype and mutant neurons. While the total number of neurites were found to be reduced only in select lines *vs.* wildtype (H3.3Q5N, TG2 C277A; **Figure 4A and 4C**), all four mutants displayed significantly reduced neurite outgrowth (**Figure 4B-C**), suggestive of deficits in dendritic maturation. We next examined synaptic architecture to determine whether disruption of QR2-H3Q5ser signaling impairs synaptic connectivity. Synapses were quantified by co-localization of the presynaptic marker SYN1 and the postsynaptic marker PSD95 (**Figure 4D and 4F**). All four mutant lines exhibited a significant reduction in synaptic density, measured as double-positive puncta relative to wildtype neurons. In addition, puncta size was significantly decreased across all mutants (**Figure 4E-F**), suggesting that disruption of H3Q5ser or its interaction with QR2 impairs synaptic maturation. Finally, to explore the impact of these perturbations, we performed multielectrode array (MEA) recordings in mature wildtype *vs.* mutant neurons. Consistent with structural and synaptic deficits observed above, all four mutant lines exhibited significant reductions in firing rates (**Figure 4G and 4K**), spiking activity (**Figure 4H and 4K**), and burst frequency (**Figure 4I and 4K**), with comparatively modest effects on burst duration (**Figure 4J and 4K**). Similar impairments were observed following pharmacological inhibition of QR2 in wildtype NGN2-induced neurons (**Figure S9M**), supporting a specific role for QR2 activity in neuronal network maturation. Together, these findings demonstrate that the H3Q5ser-dependent recruitment of QR2 to active chromatin is required for proper neuronal gene expression, as well as structural and functional maturation of human neurons.

**Figure 4.**
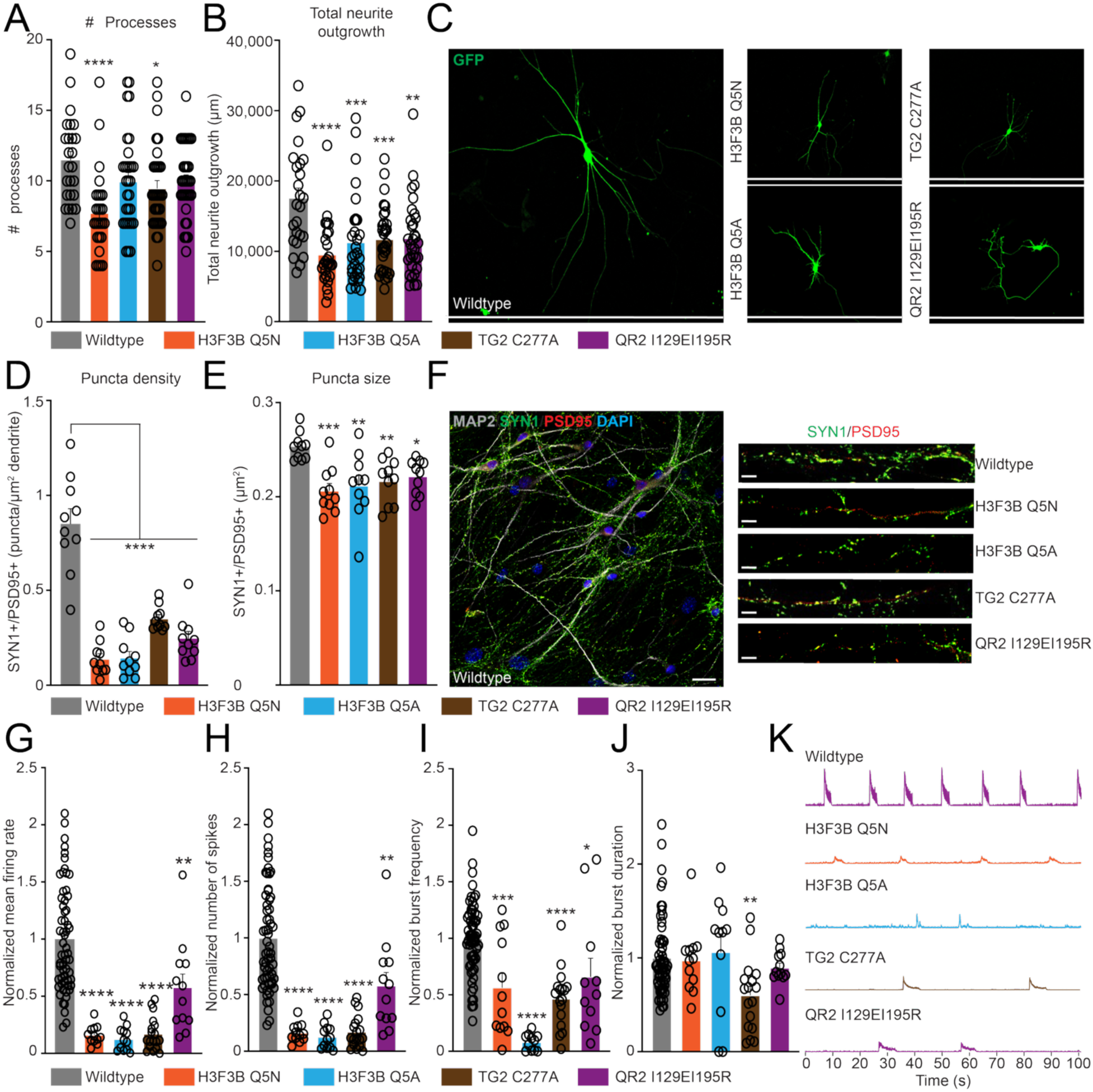
QR2-H3Q5ser interactions are required for human neuronal physiological maturation. (**A**) Quantification of the number of neuronal processes in wildtype *vs.* mutant hiPSC neurons [*n* = 25 wildtype cells, 26 H3F3B Q5N cells, 30 H3F3B Q5A cells, 31 TG2 C277A cells, and 32 QR2I129EI195R cells; one-way ANOVA, F_4,139_ = 5.105, *P*=0.0007; post hoc: Dunnett’s MC test – H3F3B Q5N (\*\*\*\**P*<0.0001) and TG2 C277A (\**P*=0.0497)]. (**B**) Quantification of total neurite outgrowth in wildtype *vs.* mutant hiPSC neurons [*n* = 25 wildtype cells, 26 H3F3B Q5N cells, 30 H3F3B Q5A cells, 31 TG2 C277A cells, and 32 QR2I129EI195R cells; one-way ANOVA, F_4,139_ = 7.222, *P*<0.0001; post hoc: Dunnett’s MC test – H3F3B Q5N (\*\*\*\**P*<0.0001), H3F3B Q5A (\*\*\**P*=0.0003), TG2 C277A (\*\*\**P*=0.0008), and QR2 I129EI195R (\*\**P*=0.0011). (**C**) Representative images of GFP labeled neurons from wildtype *vs.* mutant hiPSC neurons (scale = 625 µm). (**D**) Quantification of SYN1+/PSD95+ co-puncta density in wildtype *vs.* mutant hiPSC neurons [*n* = 10; one-way ANOVA, F_4,45_ = 43.69, *P*<0.0001; post hoc: Dunnett’s MC test **** *P*<0.0001. (**E**) Quantification of SYN1+/PSD95+ co-puncta size in wildtype *vs.* mutant hiPSC neurons [*n* = 10; one-way ANOVA, F_4,45_ = 5.723, *P*=0.0008; post hoc: Dunnett’s MC test – H3F3B Q5N (\*\*\**P*=0.0003), H3F3B Q5A (\*\**P*=0.0015), TG2 C277A (\*\**P*=0.0056), and QR2 I129EI195R (\**P*=0.0208). (**F**) Representative images of co-puncta labeling from wildtype *vs.* mutant hiPSC neurons (scale = left panel 25 µm; right panel 15 µm). MEA-based quantification of: (**G**) normalized mean firing rate [*n* = 60 wildtype cells, 12 H3F3B Q5N cells, 12 H3F3B Q5A cells, 18 TG2 C277A cells, and 12 QR2I129EI195R cells; one-way ANOVA, F_4,109_ = 32.19, *P*<0.0001; post hoc: Dunnett’s MC test – **** *P*<0.0001, ** *P*=0.0020]; (**H**) normalized spike number [*n* = 60 wildtype cells, 12 H3F3B Q5N cells, 12 H3F3B Q5A cells, 18 TG2 C277A cells, and 12 QR2I129EI195R cells; one-way ANOVA, F_4,109_ = 32.19, *P*<0.0001; post hoc: Dunnett’s MC test – **** *P*<0.0001, ** *P*=0.0020]; (**I**) normalized burst frequency [*n* = 60 wildtype cells, 12 H3F3B Q5N cells, 12 H3F3B Q5A cells, 18 TG2 C277A cells, and 12 QR2I129EI195R cells; one-way ANOVA, F_4,106_ = 22.28, *P*<0.0001; post hoc: Dunnett’s MC test – **** *P*<0.0001, *** *P*=0.0004, * *P*=0.0121]; and (**J**) normalized burst duration [*n* = 60 wildtype cells, 12 H3F3B Q5N cells, 12 H3F3B Q5A cells, 18 TG2 C277A cells, and 12 QR2I129EI195R cells; one-way ANOVA, F_4,107_ = 3.323, *P*=0.0132; post hoc: Dunnett’s MC test – ** *P*=0.0032]. (**K**) Representative MEA traces from wildtype *vs.* mutant hiPSC neurons. Data shown as mean ± SEM.

### QR2 binding to H3Q5ser co-recruits POU2F1 to loci important for neuronal maturation

Although our data indicate that QR2-H3Q5ser interactions are required for neuronal gene expression and maturation, the chromatin mechanisms underlying this effect remained unclear. We therefore next investigated how QR2 binding to H3Q5ser promotes transcriptional activation. Previous studies had shown that QR2 binds to free 5-HT, thereby inhibiting its endogenous reductase activities^37,38^. Therefore, we assessed whether QR2 binding to H3Q5ser results in similar disruptions to its enzymatic capacity. We performed QR2 enzymatic activity assays *in vitro* (again using BNAH as an electron donor) with purified wildtype QR2 *vs.* QR2 I129EI195R following pre-incubation with non-biotinylated H3_1-18_Q5ser *vs.* H3_1-18_ unmodified peptides. Consistent with its inhibition by free 5-HT, we found that wildtype QR2 displayed significantly attenuated quinone reductase activity in the presence of H3Q5ser *vs.* H3 unmodified peptides; note that such pre-incubation with H3Q5ser peptides had no effect on the QR2 mutant, which cannot bind to the mark (**Figure 5A**).

**Figure 5.**
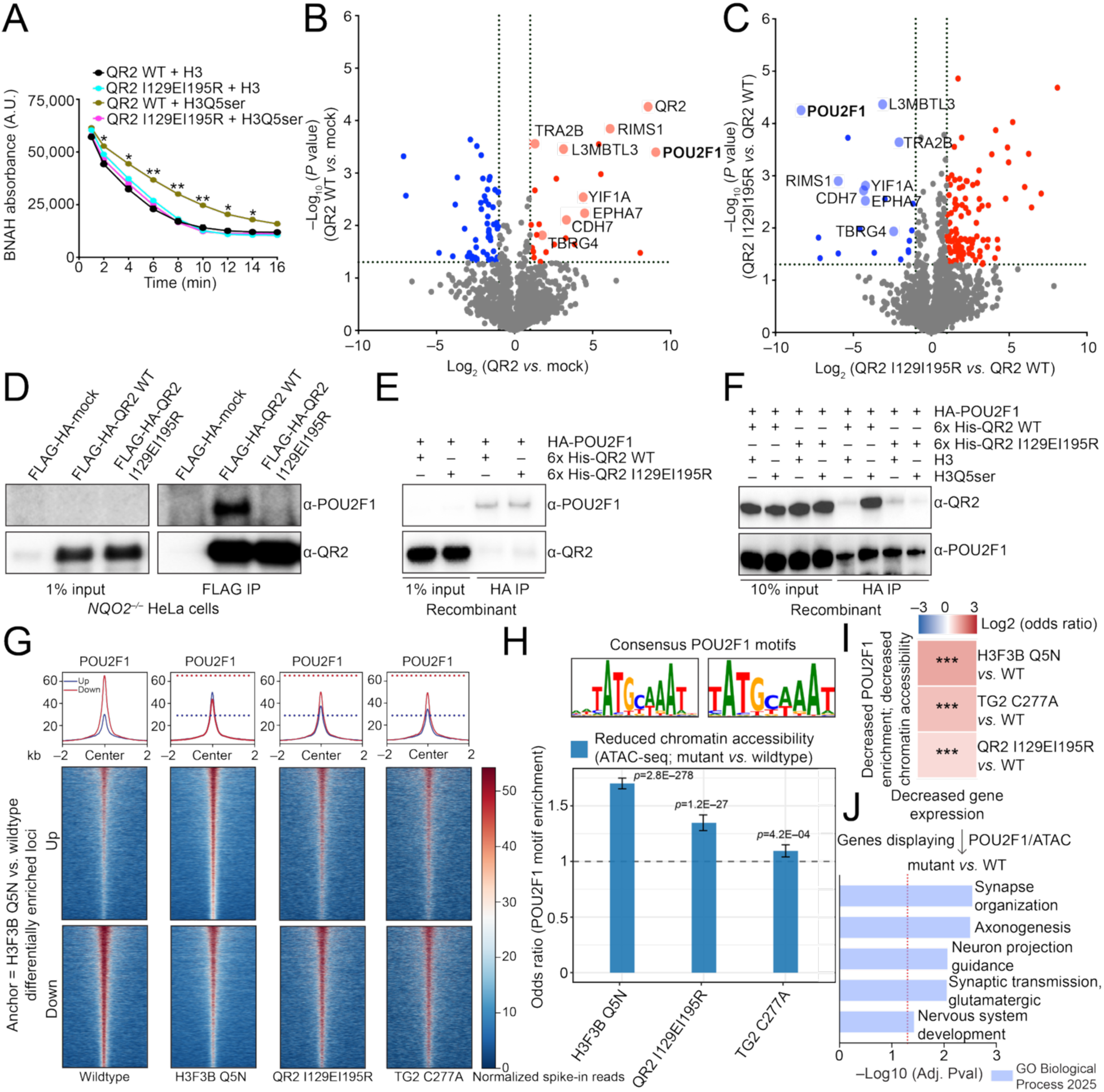
QR2 binding to H3Q5ser co-recruits POU2F1 to loci associated with neuronal maturation. (**A**) BNAH enzymatic assay comparing wildtype *vs.* I129EI129R QR2 following spike-in with either H3 unmodified or H3Q5ser peptides. *n* = 3. Two-way RM ANOVA [main effects: time (F_2.088,16.70_ = 22145, *P*<0.0001), treatment (F_3,8_ = 28.33, *P*=0.0001), and treatment x time (F_6.264,16.70_ = 84.04, *P*<0.0001); post hoc: Dunnett’s MC test (\**P*<0.05, ** *P*<0.01; QR2 wildtype H3Q5ser *vs.* QR2 wildtype H3 unmodified)]. Volcano plots of (**B**) FLAG-HA-tagged QR2 wildtype binders (*vs.* mock) from HeLa nuclear extracts (*NQO2* knockdown background) identified by immunoprecipitation and LC-MS/MS (*n* = 3; –log_10_ *P* value < 0.05; student’s t-test difference > 1), and (**C**) FLAG-HA-tagged QR2 I129EI195R binders (*vs.* FLAG-HA-tagged QR2) from HeLa nuclear extracts (*NQO2* knockdown background) identified by immunoprecipitation and LC-MS/MS (*n* = 3; –log_10_ *P* value < 0.05; student’s t-test difference > 1). (**D**) Immunoblotting validation of QR2-POU2F1 co-immunoprecipitation by FLAG-HA-tagged QR2 wildtype, but not FLAG-HA-tagged QR2 I129EI195R, from HeLa nuclear extracts (*NQO2* knockdown background). (**E**) *in vitro* QR2-POU2F1 co-immunoprecipitation experiment in the absence of H3 peptides. (**F**) *in vitro* QR2-POU2F1 co-immunoprecipitation experiment in the presence of H3Q5ser *vs.* H3 unmodified peptides. (**G**) CUT&RUN-seq heatmaps of POU2F1 enrichment at sites of differential enrichment for POU2F1 in mutant *vs.* wildtype hiPSC neurons (*n* = 3 for all; anchor: H3F3B Q5N *vs.* wildtype). Dashed lines represent wildtype signal. (**H**) Odds ratio analysis (Fisher’s exact test) comparing overlap between POU2F1 consensus motifs displaying loss of enrichment in mutant *vs.* wildtype hiPSC neurons and loss of chromatin accessibility (via ATAC-seq) at these same loci. P values are indicated. (**I**) Odds ratio analysis (Fisher’s exact test) comparing overlap between genes displaying loss of POU2F1 enrichment, reduced chromatin accessibility, and decreased gene expression in mutant *vs.* wildtype hiPSC neurons. *** *P*< 0.001. (**j**) Go Biological Process 2025 enrichment for genes identified in panel **i** (FDR<0.05). Data shown as mean ± SEM. All immunoblotting experiments repeated 3X. See **Figure S14** for uncropped blots.

Given the impact of QR2 binding to H3Q5ser on its enzymatic functions, we next hypothesized that QR2 may act to recruit additional co-factors to chromatin when bound to the serotonyl mark, which may, in turn, facilitate the PTM’s impact on permissive gene expression. For this, we revisited our established QR2 knockdown/complementation HeLa cell lines, where we next performed FLAG immunoprecipitations to enrich for QR2 (complemented wildtype QR2 *vs.* QR2 I129EI195R) in chromatin fractions, followed by LC-MS/MS analysis to identify QR2 binding partners whose interactions are dependent on the protein’s ability to engage with H3Q5ser. Following mass spectrometry, we identified numerous proteins that displayed significantly enriched binding to wildtype QR2 in the context of chromatin (**Figure 5B; Data S18**), with a number of these interactions found to be significantly attenuated in the QR2-H3Q5ser binding-dead mutant line (**Figure 5C; Data S18**). Among these candidate factors was POU2F1 (aka OCT1; **Figure 5B-5D**), a broadly expressed POU-domain transcription factor that is dispensable for self-renewal but engages developmental loci during differentiation, where it is required for lineage-appropriate gene induction^54^. To assess whether QR2 binds POU2F1 directly in the absence of a chromatin context in which H3Q5ser is present, we purified both QR2 (wildtype and mutant, 6X-His-tagged) and full length POU2F1 (HA-tagged), and performed co-immunoprecipitation assays *in vitro*. Interestingly, while QR2 and POU2F1 did not appear to interact directly in the absence of H3Q5ser (**Figure 5E**), they were found to interact following spike-in of an H3Q5ser peptide (*vs.* unmodified; **Figure 5F**), suggestive of a potential allosteric mode of recognition through which QR2 binding to the mark may facilitate structural re-organization of the complex to allow for POU2F1 binding/recruitment to chromatin.

To examine this possibility in NGN2-induced neurons, we then performed CUT&RUN-seq for POU2F1 in wildtype *vs.* mutant cells. In addition to demonstrating that POU2F1indeed co-enriches with H3K4me3Q5ser and QR2 at TSSs of permissive, RNA Pol II-marked genes (**Figure S11A**; **Data S19**), these results identified a large number of loci that were enriched for POU2F1 in wildtype neurons but displayed differential enrichment following H3Q5ser depletion or in response to disrupting QR2’s recognition of H3Q5ser (**Figure 5G**; **Figure S11B**; **Data S19**). Importantly, for those loci displaying loss of POU2F1 enrichment in mutant neurons – sites that largely mapped to consensus POU2F1 motifs – we observed that loss of chromatin accessibility at these sites (via ATAC-seq) was strongly correlated (**Figure 5H**; **Data S20**), along with a decrease in expression of corresponding genes (**Figure 5I**). Finally, for those loci displaying concomitant reductions in POU2F1 enrichment, attenuated chromatin accessibility, and decreased expression in the mutant lines, we found that these genes were highly enriched for gene ontologies related to human neuronal development and maturation, including pathways related to synapse organization, axonogenesis, glutamatergic synaptic transmission, etc. (**Figure 5J**; **Data 21**). In sum, our data suggest that during human neuronal differentiation and maturation, QR2 binds to H3Q5ser, leading to the recruitment of important lineage specific transcription factors, such as POU2F1, which, in turn, results in altered chromatin accessibility and activation of genes important for neuronal development.

### QR2-H3Q5ser interactions are required for progenitor-to-neuron transitions during human corticogenesis

NGN2 induction imposes neuronal identity directly on pluripotent cells, and the phenotypes described above therefore report on the maturation of an already-specified neuron. Whether QR2-H3Q5ser interactions are also required earlier, during the generation of neurons from cortical progenitors, cannot be asked in that system. We therefore generated dorsally patterned cortical organoids from isogenic wildtype and QR2 I129EI195R hiPSCs, using a protocol in which forebrain identity is established by dual Wnt and TGF-β inhibition over the first eighteen days and dorsal telencephalic fate then emerges without exogenous patterning cues^55^ ^56^. We profiled these organoids by single-nucleus multiome-seq, capturing the transcriptome and chromatin accessibility from the same nuclei, at one and three months – stages in this model that correspond to progenitor amplification and peak excitatory neuron diversity, with outer radial glia and diversified projection neuron classes emerging between them (**Figure S12A**)^56^. Immunostaining detected H3K4me3Q5ser and QR2 in organoids of both genotypes at one and three months (**Figure S12B**), confirming that the mark and its *reader* are expressed across the window profiled. These 1-month organoids showed expected dorsal cortical organization at the tissue level: transcripts of the dorsal telencephalic determinants *EMX1* and *EMX2*^57^ were detected by RNAscope (**Figure S12C**), and SOX2+ ventricular-zone-like regions were bordered by CTIP2+ neuronal territories (**Figure S12D**).

Joint clustering of RNA and ATAC modalities resolved nineteen populations (**Figure S12E**) that we grouped into seven broad classes – radial glia, cortical hem, Cajal–Retzius/preplate neurons, cortical excitatory neurons, interneurons, outer radial glia/astroglial progenitors, and mesenchymal-like cells – on the basis of canonical markers (**Figures S12F** and **S13**). Chromatin accessibility at marker loci was concordant with expression, most clearly at *RSPO2* in the cortical hem, *TBR1* in Cajal–Retzius/preplate neurons, *GAD2* in interneurons, and *AQP4* in outer radial glia/astroglial progenitors (**Figure S12G**). Each class carried a distinctive accessible-chromatin landscape of 21,000–38,000 peaks (**Figure S12H**), with the largest pairwise overlaps observed between developmentally adjacent populations (**Figure S12I**). Linking peaks to genes recovered cell-type-restricted regulatory modules in which accessibility and expression covaried (**Figure S12J**), including linked elements at the dorsal telencephalic determinants *EMX1* and *EMX2* (**Figure S12K**). Accessibility at these loci was restricted to the dorsal lineage: signal was present in radial glia and the cortical hem, was sustained in Cajal–Retzius/preplate neurons and cortical excitatory neurons, and was markedly lower in the interneuron and mesenchymal-like classes (**Figure S12K**), consistent with a pallial origin for the excitatory lineage.

To establish that these populations correspond to their primary counterparts, we transferred labels from a single-cell atlas of mid-gestation human neocortex^58^ excluding oligodendrocyte precursor, microglial, endothelial, and pericyte reference types, which are absent in one- to three-month organoids. The cortical classes mapped to their expected primary populations: cortical hem to ventricular radial glia (0.67), outer radial glia/astroglial progenitors to outer radial glia (0.81), cortical excitatory neurons to intermediate progenitors and deep-layer excitatory neurons (0.34 and 0.38), Cajal–Retzius/preplate neurons to deep-layer excitatory classes (0.47 and 0.35), and interneurons to caudal ganglionic eminence-derived interneurons (0.72), with mean prediction scores across classes of 0.53 to 0.79 (**Figure S12L**). The broad radial glia class was distributed across outer radial glia, ventricular radial glia, and cycling progenitors rather than resolving to one of them, as previously reported for organoid apical radial glia at early stages in this model^56^. The mesenchymal-like and mixed populations did not map to any single primary class, consistent with their exclusion from the analyses above. Together with the expected dorsal cortical composition, the transcriptional fidelity to human fetal cortex, and the tissue-level organization described above, the dorsal restriction of *EMX1* and *EMX2* accessibility establishes that these organoids recapitulate a dorsal cortical program.

Cellular composition shifted from progenitor- to neuron-dominated between one and three months, and diverged between genotypes (**Figure 6A**). At one month, organoids of both genotypes were dominated by dorsal radial glia and the other dorsal progenitor classes. By three months, wildtype organoids had acquired RORB+ excitatory neuron subtypes and outer radial glia/astroglial progenitors; in QR2 I129EI195R organoids, these populations were poorly represented, and immature interneuron classes instead accounted for the largest share of cells. These interneurons expressed *GAD1*, *GAD2,* and the telencephalic determinant *FOXG1* but lacked detectable *NKX2-1* and *LHX6*, and were assigned by label transfer to caudal rather than medial ganglionic eminence interneurons (0.72 versus 0.18; **Figure S12L**). They are therefore dorsally derived telencephalic interneurons resembling caudal ganglionic eminence-derived cells, not ventrally derived contaminants, consistent with the interneuron populations described in this organoid system and with the dorsal origin of a subset of human cortical interneurons^56^. Their increased representation in the mutant reflects the depletion of excitatory neurons rather than increased interneuron production. Disrupting QR2-H3Q5ser binding therefore did not change which cell types the organoids could produce, but changed their relative output.

**Figure 6.**
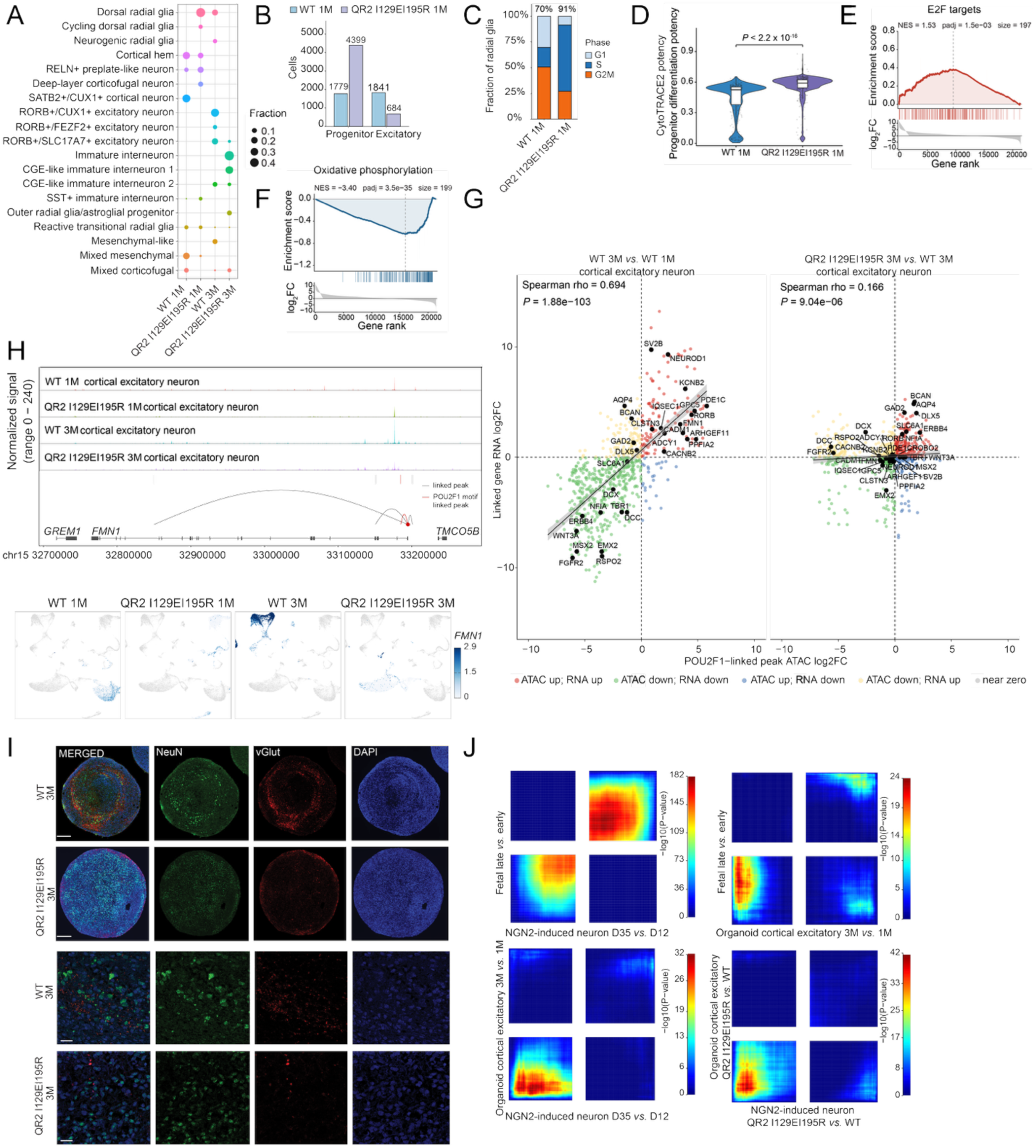
QR2–H3Q5ser interactions are required for human cortical neuron development in a human organoid model. (**A**) Cell-type composition of wild-type (WT) and QR2 I129EI195R organoids at 1 and 3 months (1M, 3M). Dot area shows the fraction of cells in each group (columns sum to 1). Colour shows lineage, as in Figure S13B. CGE, caudal ganglionic eminence. (**B**) Numbers of dorsal progenitors (radial glia and cortical hem) and cortical excitatory neurons at 1M, based on the broad annotation in **Figure S12F**. (**C**) Cell-cycle phase of 1M radial glia (Seurat CellCycleScoring). The percentage of cycling (S + G2/M) cells is shown above each bar. G1 includes G0. (**D**) CytoTRACE 2 differentiation potency of 1M dorsal progenitors; higher values indicate a less differentiated state. Box, median and interquartile range; points, a random subsample of 350 cells per genotype. n = 1,779 (WT) and 4,399 (mutant) cells; Mann–Whitney U test; Cliff’s δ = 0.61; Δmedian = 0.07 (95% CI 0.06–0.07). (**E** and **F**) Pseudobulk GSEA of Hallmark gene sets in radial glia, mutant versus WT, with genes ranked by log₂ fold change (left, higher in mutant). (**E**) E2F targets. (**F**) Oxidative phosphorylation. NES, normalized enrichment score; padj, adjusted p value. Full results are in **Table S22**. (**G**) Change in expression versus change in accessibility for genes linked to POU2F1 motif-containing peaks in cortical excitatory neurons (positive peak–gene links, score ≥ 0.10). Left, WT 3M versus 1M; right, mutant versus WT at 3M. Black points, curated lineage genes; line, linear fit with 95% CI. Spearman ρ and p are shown. Groups with <50 cells were excluded. (**H**) Top, ATAC coverage at the *FMN1* locus in cortical excitatory neurons, with peak– gene links below; the POU2F1 motif-containing linked peak is in red. Bottom, *FMN1* expression on the joint UMAP. (**I**) Immunostaining for NeuN and vGluT1 with DAPI in 3M organoids. Scale bars, 200 μm (top) and 20 μm (bottom). (**J**) Rank–rank hypergeometric overlap between differential expression signatures, left to right and top to bottom: NGN2-induced neurons (DIV35 versus DIV12) against fetal cortex (late versus early); Organoid excitatory neurons (3M versus 1M) against fetal cortex (late versus early); NGN2-induced neurons (DIV35 versus DIV12) against organoid excitatory neurons (3M versus 1M); NGN2-induced neurons against organoid excitatory neurons, mutant versus WT.

That divergence was already evident at one month in the balance between dorsal progenitors and the excitatory neurons to which they give rise. Mutant organoids contained roughly 2.5-fold more dorsal progenitors (broad radial glia and cortical hem classes; 4,399 versus 1,779 nuclei) than wildtype, yet 2.7-fold fewer cortical excitatory neurons (684 versus 1,841), so that the excitatory-to-progenitor ratio fell from 1.03 to 0.15, a 6.7-fold reduction (**Figure 6B**). An excess of progenitors accompanied by a deficit of their progeny is the signature of a transitional block rather than of neuronal loss after specification, and two further measurements describe the state in which the retained progenitors are held.

Mutant radial glia were more frequently cycling (91% *vs.* 70% in S or G2/M), and the shift was specifically into S phase (65% *vs.* 19%), with a reciprocal fall in G2/M (27% *vs.* 51%; (**Figure 6C**) – a redistribution within the cycle rather than a uniform increase in proliferation. Mutant radial glia were also assigned systematically higher differentiation potency by CytoTRACE 2^59^, which infers potency from learned gene-set signatures rather than from hand-curated stemness markers; that is, they were transcriptionally less differentiated than their wildtype counterparts (median 0.59 *vs.* 0.53; Δmedian = 0.07, 95% CI 0.06–0.07; Cliff’s δ = 0.61; Figure **6D**). A curated stemness-gene module gave a concordantly large effect (Cliff’s δ = 0.77), indicating that the difference does not depend on the CytoTRACE 2 model. These two readouts are not independent – potency scores of this kind rise with proliferative expression, and the genotypes differ in cycling fraction – so we take them as two descriptions of a single retained progenitor state rather than as separate lines of evidence.

To identify the programs that distinguish the retained mutant progenitor pool, we performed gene set enrichment analysis on pseudobulk profiles of radial glia (excluding the cortical hem population) from mutant and wildtype organoids at one month, ranking genes by the TMM-normalized log₂ fold change between genotypes (**Data S22**). Two Hallmark sets were significantly enriched among transcripts higher in the mutant, and both relate to the cell cycle: E2F targets (NES = 1.53, adjusted p = 1.5 × 10⁻³, 197 genes; **Figure 6E**) and the G2M checkpoint (NES = 1.49, adjusted p = 4.0 × 10⁻³, 192 genes; **Data S22**). Because E2F activity promotes the G1-to-S transition, this is consistent with the S-phase enrichment of mutant radial glia in **Figure 6C**. The most strongly depleted set was oxidative phosphorylation (NES = −3.40, adjusted p = 3.5 × 10⁻³⁵, 199 genes; **Figure 6F**). Mitochondrially encoded transcripts were excluded before ranking, so this signal reflects nuclear-encoded genes. Depletion was broad, with 43 of the 50 Hallmark sets returning a negative enrichment score (25 of them significant), so we do not interpret the depleted sets further. Together with continued cycling (**Figure 6C**) and higher inferred potency (**Figure 6D**), these data suggest that mutant radial glia at one month remain in cycle rather than progressing toward neuronal differentiation, consistent with a delay at the progenitor-to-neuron transition. By three months, the mutant progenitor pool is itself nearly exhausted (**Figure 6A**). Across the two timepoints, the phenotype is therefore better described as delayed and ultimately failed neurogenesis with progenitor depletion than as a stable expansion of the progenitor pool. Because each genotype is represented by a single pooled sample, none of these comparisons carries a between-sample estimate of variance. We therefore describe the per-cell potency difference (**Figure 6D**) by its effect size and interval, as a measure of magnitude rather than significance, and treat the enrichment p values, computed by gene permutation on one pseudobulk profile per genotype, as measures of how coherently a set is ranked within that contrast. These differences remain to be confirmed in additional organoid batches.

We next asked how loss of mark recognition affects the regulatory elements through which QR2 acts. Because QR2 associates with POU2F1 in an H3Q5ser-dependent manner (**Figures 5D-F**), we examined positively correlated peak-to-gene links at POU2F1 motif-containing elements in cortical excitatory neurons. Across normal maturation, the accessibility of these elements and the expression of their linked genes were strongly coupled (Spearman ρ = 0.69; **Figure 6G, left**): maturation and synaptic genes, among them *NEUROD1*, *RORB*, *KCNB2*, *GPC5*, *PDE1C*, *SV2B*, and *CADM1*, gained both accessibility and expression, while early cortical and progenitor genes, among them *WNT3A*, *RSPO2*, *EMX2*, *FGFR2*, *MSX2*, *TBR1,* and *DCX*, lost both. POU2F1-linked elements therefore open and close in register with the normal progenitor-to-neuron transition. In the mutant this coupling largely collapsed: across the same links, accessibility and expression changes between mutant and wildtype organoids at three months were nearly flat (ρ = 0.17), and the regression slope flattened relative to wildtype maturation (**Figure 6G, right**; **Data S23**). Maturation genes no longer moved coherently; instead, the links that did gain accessibility were associated with interneuron and glial identity – *GAD2*, *DLX5*, *ERBB4*, *SLC6A1*, *AQP4*, *BCAN*, and *NFIA*. POU2F1-linked elements are thus uncoupled from the excitatory maturation trajectory in the mutant, and partly redirected toward non-excitatory programs, which matches the arrested maturation and the and the shift in cellular output seen in the composition and potency analyses. Because POU2F1 physically associates with QR2, this decoupling offers a candidate regulatory route from loss of mark recognition to the excitatory neuron deficit.

*FMN1* illustrates this pattern at a single locus. A distal POU2F1 motif-containing element linked to the *FMN1* promoter gained accessibility in wildtype cortical excitatory neurons between one and three months, accompanied by induction of *FMN1* in those cells; in mutant organoids neither the accessibility gain nor the induction occurred (**Figure 6H**). *FMN1* encodes a formin required for neurogenin 3-induced dendritogenesis and synaptogenesis: in cultured mouse hippocampal neurons, overexpressing the Fmn1 Ib isoform increases primary dendrite number and glutamatergic synaptic input, and knocking *Fmn1* down prevents neurogenin 3 from producing either effect^60^. Its function in human cortical neurons has not been tested directly, and we present it here as one of a set of POU2F1-linked maturation genes that fail to be induced when the mark can no longer be read. These regulatory deficits were mirrored by the cellular output of the tissue. At three months, wildtype organoids contained abundant NeuN+ neurons with dense vGluT1+ puncta organized into cortical-plate-like territories, whereas mutant organoids showed markedly reduced NeuN and vGluT1 immunoreactivity despite comparable nuclear density (**Figure 6I**).

Finally, we asked whether the maturation program disrupted in organoids is the same one engaged in two-dimensional culture and *in vivo*. By RRHO, genes changing with maturation in NGN2 neurons (DIV35 *vs.* DIV12) overlapped strongly and concordantly with those separating late from early human fetal, as did genes changing in organoid cortical excitatory neurons between one and three months; the two *in vitro* maturation programs agreed with one another (**Figure 6J**). Most informatively, the transcriptional consequences of QR2 I129EI195R were themselves concordant across the two systems (**Figure 6J**), indicating that the same *reader*-dependent program is disrupted whether neuronal identity is imposed by NGN2 or arises through endogenous corticogenesis.

## DISCUSSION

Here, we explored the possibility that in addition to its previously described roles as a modifier of H3K4me3-mediated *reader* interactions and permissive gene expression, H3Q5ser may recruit additional dedicated binding proteins that facilitate its impact on transcription within mammalian cells. Employing proteomic-, structural-, molecular-, epigenomic-, and cellular-based approaches, we found that the quinone reductase enzyme, QR2, is a *bona fide reader* of H3Q5ser, both *in vitro* and *in cellulo*, and that this interaction can occur in the absence or presence of adjacent H3K4me3. A structure-guided pair of mutations that abrogates this binding, without affecting QR2’s expression, subcellular localization, or catalysis, dysregulated neuronal gene expression and produced deficits in neuronal morphology, synaptic connectivity and electrophysiology; in dorsally patterned cortical organoids, the same allele caused dorsal progenitors to accumulate at the expense of cortical excitatory neurons, implicating these interactions in the progenitor-to-neuron transition as well. While QR2 is long established as a cytosolic quinone reductase, its binding to H3Q5ser attenuates its enzymatic activities *in vitro*; however, such binding then allows for the recruitment of additional transcription factors, such as POU2F1, which play key roles in promoting gene expression programs that are required for appropriate patterns of neuronal differentiation and maturation.

QR2 recognizes H3Q5ser by a mechanism unlike that of the *readers* characterized to date. The TAF3 PHD finger engages the serotonyl group through a CH–π contact with a proline stabilized by a conserved tryptophan^17^, and WDR5 accommodates it in a shallow hydrophobic pocket, combining a hydrogen bond with van der Waals contacts from three aromatic side chains^18^; in neither case does a cofactor participate. In QR2, the serotonyl indole instead stacks against the flavin ring of the bound FAD, and apo-QR2 does not bind H3Q5ser (**Figure S2C**), placing the cofactor itself within the *reading* surface. Redox cofactors in chromatin-associated enzymes are catalytic rather than recognitional: with LSD1 (an FAD-dependent demethylase), for example, the flavin oxidizes methylated Lys4^61^ and H3Q5ser is excluded from that active site by steric clash^15^. The modification that blocks one flavoenzyme is accommodated in the catalytic pocket of another. A cofactor contributing directly to the recognition of a histone modification is, to our knowledge, without precedent among histone PTM *readers*, raising the possibility that cofactor-binding enzymes may also serve as *readers* for H3Q5dop, H3Q5his, and other aromatic marks.

While our data establish that QR2’s interactions with H3Q5ser in chromatin support permissive gene transcription, particularly within neurons, it remains unclear to what extent these interactions account for QR2’s previously documented roles within cells. Remarkably little is known about QR2^34^: it is not known which endogenous quinones, if any, QR2 reduces in cells; the only co-substrate it uses efficiently, NRH, has no established endogenous source; and rather than detoxifying reactive oxygen species, as might be expected of a reductase, QR2 appears to generate them, such that its inhibition reduces oxidative and metabolic burden and reverses Alzheimer’s disease phenotypes in mice^40^. Its canonical quinone-reductase activity is therefore unlikely to represent a major cellular function *in vivo*. Even so, QR2 is not a decaying gene: across 16 orthologues spanning lamprey to human, it remains strongly conserved and retains its FAD-binding scaffold (**Figure S5E-G**), indicating continued purifying selection for a maintained role rather than pseudogene-like decay. More tellingly, the two functional surfaces diverge sharply in their conservation: position 129 is hydrophobic or aromatic in all sixteen orthologues and position 195 is the human isoleucine throughout the primate lineage, whereas the catalytic residue 162 cycles freely among asparagine, serine and arginine, even amongst mammals (**Figure S5F**). Because the cofactor is itself part of the *reading* surface, conservation of the FAD-binding scaffold is evidence for *reader* function and not catalysis alone; the functional separation of the two rests, in addition, on our binding- and catalysis-dead mutants. This pattern maps onto the history QR2 shares with its paralog: QR1 and QR2 arose from a gene duplication approximately 460 million years ago, after which QR2 lost efficient use of NAD(P)H – a divergence since described as neofunctionalization toward a conditional pseudoenzyme^35,36^. QR1 also differs from QR2 at the active-site positions that form the serotonyl-reading surface, including residues 129 and 195, and carries an additional C-terminal domain absent in QR2, so the same divergence that altered QR2’s co-factor use plausibly created its capacity to read H3Q5ser – and explains why the interaction is specific to QR2. Chromatin reading, rather than quinone reduction, may therefore be a – or perhaps the – principal conserved function of QR2 in cells.

Four genetically distinct perturbations converge on the same outcome in 2D NGN2-induced neurons. The H3F3B Q5A and Q5N knock-ins remove the acceptor residue, the TGM2 C277A inactivates the enzyme that installs the mark, and the QR2 I129EI195R removes the protein that reads it; each of the four reduced neurite outgrowth, synaptic density and SYN1/PSD95 puncta size, along with firing, spiking, and burst frequency on multielectrode arrays (**Figures 4B–I and 4K**). That four alleles acting at three different steps of one axis produce a single phenotype argues that the phenotype reports on the axis rather than on any individual allele. TGM2 C277A is particularly informative in this respect, since it acts on the *writer* rather than on the Q5 side chain, separating loss of monoaminylation from any effect of substituting the residue immediately adjacent to Lys4.

Induced neuronal models impose identity rather than letting it emerge, so they can report on how a neuron matures but not on whether it is made. By placing the same separation-of-function mutation in a dorsally patterned organoid, we were able to ask whether *reading* H3Q5ser is required earlier during the generation of neurons from cortical progenitors, it is. QR2 I129EI195R organoids accumulate dorsal progenitors while producing fewer excitatory neurons, so the defect is not a failure to specify a cortical progenitor – pallial identity is intact by composition, by accessibility at *EMX1* and *EMX2*, and by correspondence to primary fetal classes – but a failure to convert those progenitors into neurons. The mark is therefore required at two separable points in the same lineage: the progenitor-to-neuron transition, which the organoid exposes, and the subsequent maturation of the neuron, which the induced model exposes. That the same *reader*-dependent program is disrupted in two systems that reach neuronal identity by different routes is the most informative result. Induced neurons bypass the progenitor stage entirely, while organoids pass through it. Yet both maturation programs track the direction of change in human fetal cortex, and, in both, the mutation drives the transcriptome counter to that direction (**Figures 3H and 6J**). Agreement between these two models that reach neuronal identity through different routes places the transcriptional program on which QR2-H3Q5ser acts within the cortical excitatory lineage itself rather than in either culture system.

The mechanism that we propose runs through chromatin. Across normal maturation, POU2F1 motif-containing elements and their linked genes move together, opening at maturation and synaptic genes and closing at early cortical ones; in the mutant, that coupling is lost, and the elements that do gain accessibility belong to interneuron and glial programs rather than to excitatory ones. Because QR2 associates with POU2F1 in an H3Q5ser-dependent manner, the simplest interpretation is that losing the mark releases POU2F1-associated elements from the excitatory maturation trajectory. Two caveats bound that assumption: the octamer motif is shared across the POU family, and class III factors, such as POU3F2 and POU3F3, are the ones that are conventionally associated with the cortical excitatory program, so motif enrichment alone cannot assign these elements to POU2F1; the physical interaction, not the motif, carries that weight. Separately, POU2F1 is a documented stress-responsive factor whose target selectivity shifts after oxidative and genotoxic stress^62,63^ and whose loss shifts cells away from glucose metabolism and toward mitochondrial activity^62^. A factor with those properties sits at the junction of the chromatin and metabolic observations in our data, which is a reason to treat them as potentially coupled rather than as independent lines of evidence.

A further consequence of QR2 occupancy is protection of the mark itself. H3Q5 monoaminylation has no dedicated *eraser*; TGM2 acts as *writer*, *eraser*, and *exchanger* at the same residue^13^, and its activity on chromatin is dictated by steric accessibility rather than by primary sequence or pre-existing modifications^16^, so a stably bound *reader* would be expected to occlude the enzyme from its substrate. Consistent with this, H3K4me3Q5ser levels in QR2 I129EI195R neurons were reduced to a degree comparable to that seen in the H3.3Q5A and Q5N lines (**Figure S9G**), consistent with QR2 occupancy stabilizing the mark against TG2-mediated erasure (**Figure S10A-B**). *Reader*-mediated protection of a histone modification from its own *writer*–*eraser* has, to our knowledge, no direct precedent, although within the H3 tail itself, Q5 serotonylation protects adjacent H3K4me3 from demethylation^15^. QR2 binding therefore appears to stabilize the very mark it *reads*, providing a feed-forward mechanism by which a transient neurotransmitter signal could be converted into a more persistent chromatin state.

Beyond development, QR2 has been studied in two other settings, and in both, the same ambiguity applies. In the mature brain, QR2 protein and transcript are elevated in the hippocampus and frontal cortex of Alzheimer’s disease patients^41–43^, and human *NQO2* variants associate with cognitive decline and neuropsychiatric disease^45^. Rodent studies establish the direction of that relationship: reducing QR2 expression improves novel memory^44^, and cholinergic and dopaminergic pathways that suppress QR2 gate novel memory formation^42,64,65^. Outside the brain, QR2 is expressed across human tissues – heart, lung, kidney and skeletal muscle, and abundantly in liver, where QR1 levels are low– and constitutive knockout in mice causes myeloid hyperplasia^66,67^ and other malignancies^68^. Critically, every one of these manipulations was global, or used inhibitors such as yb537 or yb800 that also abrogate QR2 binding to H3Q5ser, so none can resolve which phenotypes reflect QR2’s enzymatic functions and which reflect its role in H3Q5ser-mediated gene expression. A separation-of-function allele of the kind described here offers a route to separating them, and given H3 serotonylation’s broad distribution across tissues, we posit that QR2’s binding to the mark plays a prominent, and perhaps evolutionarily conserved, role beyond the nervous system.

## Supporting information

Data S5_H3K4me3Q5ser_QR2

Data S5_RNA Pol II

Data S11_H3K4me3Q5ser

Data S20_ATAC_H3F3BQ5A_vs_WT_annotated_peaks

Data S20_ATAC_H3F3BQ5N_vs_WT_annotated_peaks

Data S20_ATAC_I129EI195R_vs_WT_annotated_peaks

Data S20_ATAC_TG2C277A_vs_WT_annotated_peaks

## Key resources table

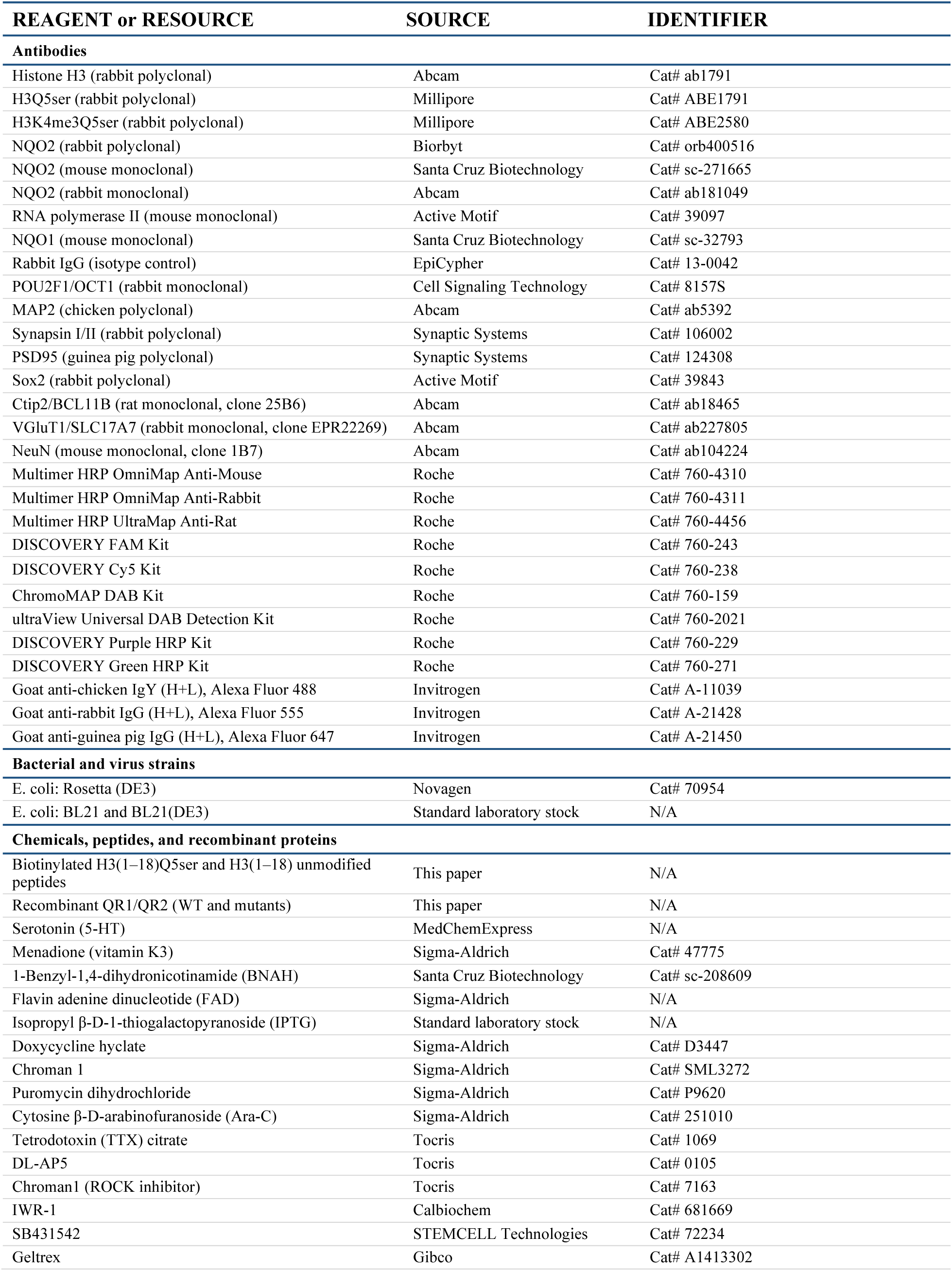
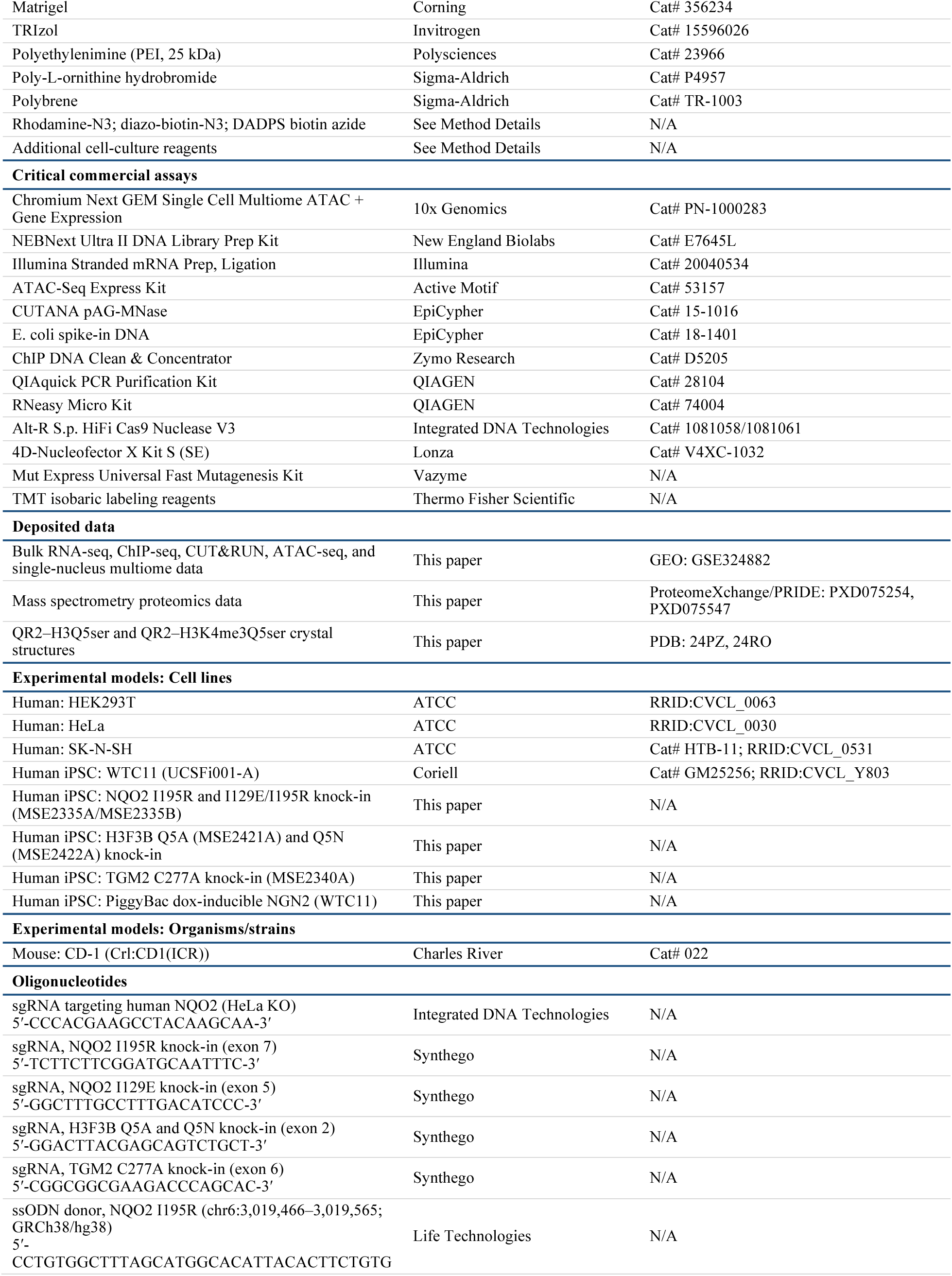
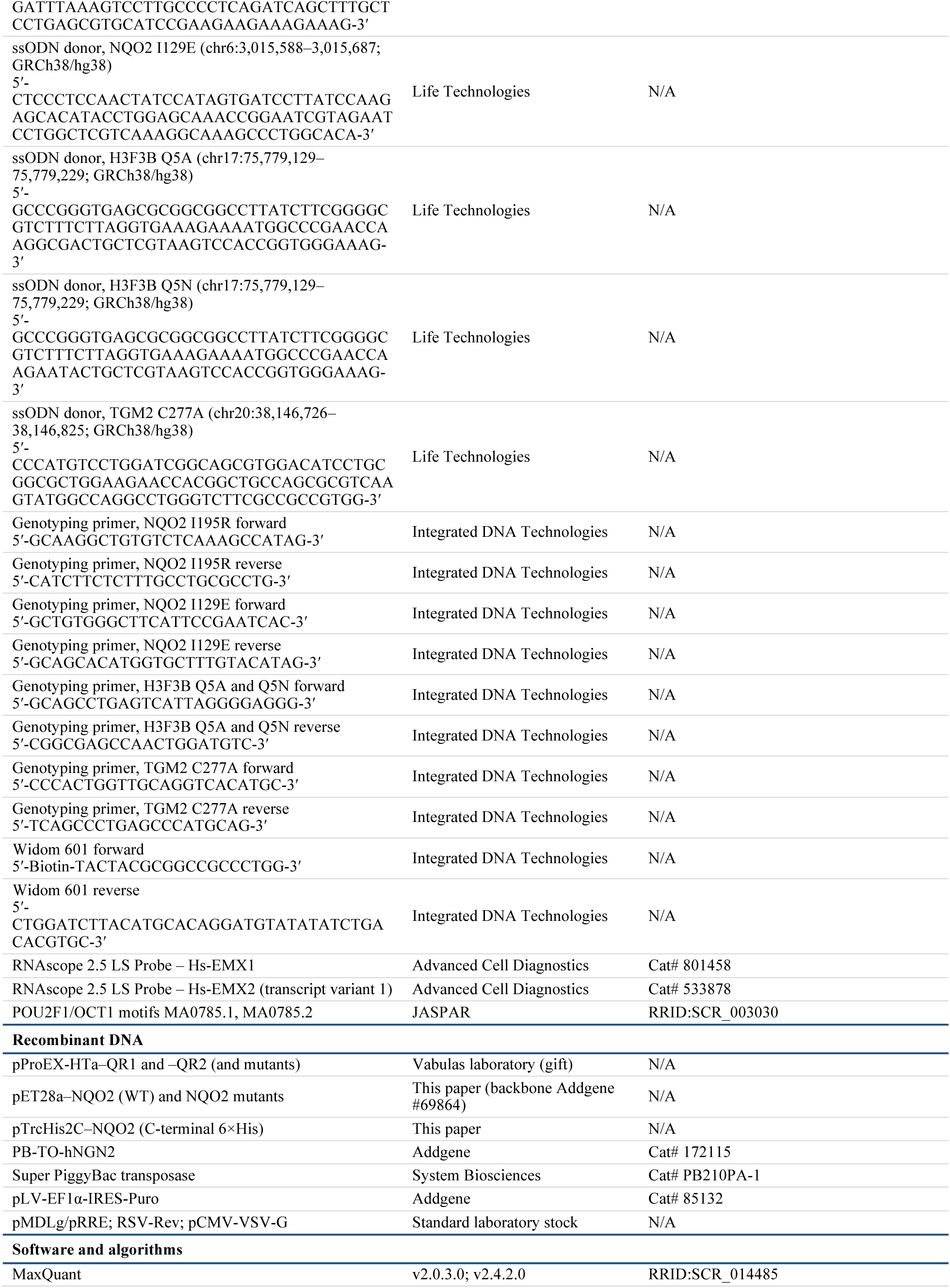
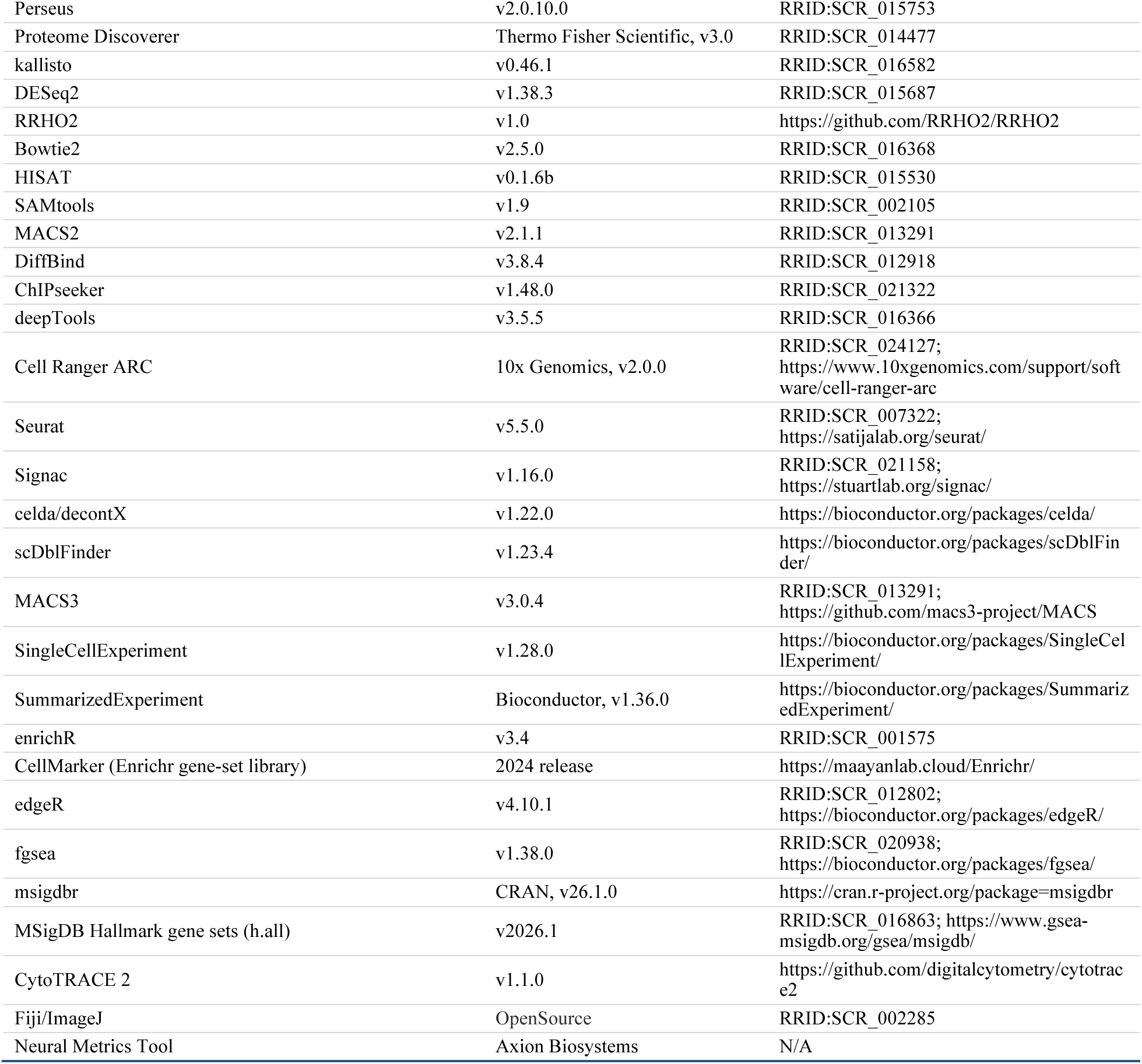

### Resource availability

#### Lead contact

Further information and requests for resources and reagents should be directed to and will be fulfilled by the lead contact, Ian Maze.

#### Materials availability

Plasmids and genome-edited iPSC and HeLa cell lines generated in this study are available from the lead contact under a completed Materials Transfer Agreement.

#### Data and code availability

- All next-generation sequencing data (bulk RNA-seq, ChIP-seq, CUT&RUN, ATAC-seq, and single-nucleus multiome) have been deposited at GEO; mass spectrometry proteomics data have been deposited at ProteomeXchange/PRIDE; and atomic coordinates and structure factors have been deposited at the Protein Data Bank. All are publicly available as of the date of publication. Accession numbers are listed in the key resources table.
- Any additional information required to reanalyze the data reported in this paper is available from the lead contact upon request.

## Experimental model and study participant details

### HEK 293T and HeLa cell culture

HEK 293T and HeLa cell lines were obtained from the American Type Culture Collection (ATCC) and grown in DMEM supplemented with 10% FBS with 100 μg ml^−1^ streptomycin and 100 U ml^−1^ penicillin. Cells were maintained at 37 °C in a 5% CO_2_, 95% humidified incubator.

### Culturing of SK-N-SH cell

SK-N-SH cells (American Type Culture Collection, HTB-11) were cultured in DMEM supplemented with 10% fetal bovine serum (FBS), 100 U/mL penicillin, and 100 μg/mL streptomycin. Cells were maintained in a humidified 37 °C incubator with 5% CO_2_.

### Human iPSC maintenance

The WTC11 human pluripotent stem cell (hPSC) line (hPSCreg UCSFi001-A)^52^ was obtained from Coriell Institute for Medical Research, GM25256. Cells were maintained on six-well tissue culture plates coated with Geltrex diluted in DMEM/F-12 and cultured in StemFlex medium. Cultures were maintained at 37 °C in a humidified incubator with 5% CO₂, with medium changes performed every other day. Cells were passaged at approximately 80% confluency using 0.5 mM EDTA in DPBS at room temperature. Detached cells were resuspended in StemFlex medium supplemented with 5 nM Chroman1 and reseeded at split ratios adjusted according to cell density. The following day, the medium was replaced with fresh StemFlex medium lacking Chroman1, and cultures were subsequently maintained using this routine passaging protocol.

### Generation of human *NQO2*, *H3F3B*, and *TGM2* knock-in iPSC lines

Homozygous single- and double-knock-in human iPSC lines were generated in the WTC11 background using CRISPR–Cas9–mediated homology-directed repair (HDR). Briefly, 1 × 10⁶ WTC11 iPSCs were electroporated using a 4D-Nucleofector system (Lonza; program V4XP-3024) with preassembled ribonucleoprotein complexes containing 10 µg S. pyogenes Cas9 nuclease (IDT, 1081058), 8 µg synthetic single-guide RNA (sgRNA; Synthego, CRISPRevolution sgRNA EZ Kit), and 100 pmol single-stranded oligodeoxynucleotide (ssODN) donor template (Life Technologies). Following electroporation, cells were replated on Geltrex-coated plates and cultured in mTeSR Plus medium. Five days after electroporation, cells were dissociated with Accutase and single cells were sorted into 96-well plates using a WOLF benchtop microfluidic cell sorter (Nanocellect) in mTeSR Plus supplemented with CloneR. After 13 days of clonal expansion, genomic DNA was extracted from individual clones using QuickExtract DNA Extraction Solution (Lucigen, QE090500). Correct HDR events were identified by locus-specific PCR using GoTaq DNA Polymerase (Promega) followed by Sanger sequencing.

For *NQO2*, HDR was first used to introduce the I195R mutation in exon 7. Clone MSE2335A was identified as a correctly edited homozygous knock-in clone, whereas clone MSE2335B served as an isogenic wild-type control. Clone MSE2335A was subsequently subjected to a second round of CRISPR–Cas9 editing to introduce the I129E mutation in exon 5, generating a homozygous double-mutant line. For *H3F3B*, HDR was used to introduce the Q5A mutation in exon 2, yielding clone MSE2421A. A separate HDR strategy targeting the same exon was used to generate the Q5N mutation, and clone MSE2422A was used for downstream experiments. For *TGM2*, HDR was used to introduce the C277A mutation in exon 6, and clone MSE2340A was identified as a correctly edited homozygous knock-in line and used for subsequent analyses. Genomic integrity and normal karyotype were confirmed for all edited clones by SNP array analysis. Sequences of sgRNAs, ssODN donor templates, and PCR primers used for CRISPR–Cas9–mediated HDR are provided in the key resources table.

### Genomic integrity of human iPSC lines

The WTC-11 donor human iPSC line was tested for genomic integrity at passage 36 using SNP-array technology (Global Diversity Array v1.0 BeadChip, Illumina). All the edited clones were also tested at passage 4 after the genome editing. No detection of CNV larger than 1.5 Mb or AOH larger than 3 Mb were detected on somatic chromosomes. The typical WTC-11 deletion of 2.9 Mb on Yp11.2 was detected in all clones. This deletion is known to be present in the donor, from whom the cell line was derived^52^.

### Generation of PiggyBac-integrated hPSC lines

For stable transgene integration, WTC11 hPSCs were seeded at a density of 3–4 × 10⁵ cells per well in six-well tissue culture plates in StemFlex medium supplemented with Chroman1 and cultured overnight. On the following day, cells were transfected with a PiggyBac transgene plasmid encoding doxycycline-inducible human NGN2 (PB-TO-hNGN2; Addgene #172115) together with Super PiggyBac transposase (System Biosciences, PB210PA-1) using Lipofectamine Stem Transfection Reagent. Transfection complexes were prepared by separately diluting 8 μL of Lipofectamine Stem Transfection Reagent in 125 μL Opti-MEM I reduced-serum medium and mixing 500 ng of transposase plasmid and 1 μg of PB-TO-hNGN2 plasmid in 125 μL Opti-MEM I. The two solutions were then combined and incubated for 10 min at room temperature to allow complex formation. A total of 250 μL of the transfection mixture was added dropwise to cells pre-equilibrated in 2 mL of warm Opti-MEM I. Following a 4 h incubation at 37 °C, 2 mL of warm StemFlex medium was added, and cells were cultured overnight. On the subsequent day, the medium was replaced with 1.5 mL of fresh StemFlex medium. Selection was initiated on day 3 by switching to StemFlex medium supplemented with 2 μg mL⁻¹ puromycin and maintained for 4–6 days, with medium changes every other day. To ensure uniform selection and eliminate false-positive cells, cultures were passaged once in puromycin-containing StemFlex medium prior to further expansion.

### Mice

CD1 mice (Charles River 022) were housed in specific-pathogen-free (SPF) conditions and environmentally controlled animal facilities with a 12-hour light-dark cycle and were given unrestricted access to food and water. All animal protocols and experiments were approved by the Institutional Animal Care and Use Committee (IACUC) of ISMMS. All experiments were performed following strict IACUC guidelines.

### Human brain tissue

Postmortem human brain frozen cortex tissues were obtained from patients who had enrolled in and provided consent for the brain donation program through the Neuropathology Brain Bank & Research CoRE at Mount Sinai, RRID: SCR_027565. These samples were collected according to ethical guidelines and institutional review board (IRB) approval, ensuring the privacy and dignity of the donors while supporting ongoing neuropathological research. All specimens obtained were de-identified. Brain specimens were dissected by expert neuroanatomists into defined blocks. Nuclei isolation and peptide pulldown assays were conducted, as previously described. Two male fetal brain frontal cortex samples of 32 weeks and 35 weeks were used for H3K4me3Q5ser and QR2 immunohistochemistry evaluation. An additional cohort of fresh-frozen postmortem human brain tissue (frontal cortex) was obtained from the Iowa Neuropathology Resource Laboratory (INRL) at the University of Iowa (project #12425051106). Eight de-identified cases spanning mid-gestation through the early postnatal period were analyzed: six second-trimester fetal specimens with gestational ages of 18–23 weeks, together with two perinatal cases (37 weeks’ gestation/19 days postnatal and 41 weeks’ gestation/1 day postnatal). The cohort included five male and three female donors, and postmortem intervals ranged from 8 to 41 hours; case-level information is provided in Table S3. All samples were received fully de-identified.

## METHOD DETAILS

### Cell culture materials

DMEM (Gibco, 10569044); fetal bovine serum (FBS; Gibco, A5670801); penicillin–streptomycin (Gibco, 15140122); Basal Medium Eagle (Gibco, 21010046); HyClone bovine growth serum (Cytiva, SH30541.03); GlutaMAX™ supplement (Gibco, 35050061); potassium chloride (KCl; Invitrogen, AM9640G); tetrodotoxin citrate (TTX; Tocris, 1069); DL-2-amino-5-phosphonopentanoic acid (DL-AP5; Tocris, 0105); Geltrex™ matrix (Gibco, A1413302); DMEM/F-12 (Gibco, 11320033); StemFlex™ medium (Gibco, A3349401); EDTA (Invitrogen, AM9260G); Dulbecco’s phosphate-buffered saline (DPBS; Gibco, 14190144); Chroman 1 (Sigma, SML3272); doxycycline hyclate (Sigma, D3447); Opti-MEM™ I reduced-serum medium (Gibco, 31985070); Lipofectamine™ Stem transfection reagent (Invitrogen, STEM00015); Lipofectamine™ 3000 transfection reagent (Invitrogen, L3000008); puromycin dihydrochloride (Sigma, P9620); Accutase (STEMCELL Technologies, 07920); Neurobasal™ medium (Gibco, 21103049); Neurobasal™-A medium (Gibco, 10888022); B-27™ supplement (Gibco, 17504044); cytosine β-D-arabinofuranoside (Ara-C; Sigma, 251010); calcium chloride (CaCl₂; Sigma, 21115); magnesium chloride (MgCl₂; Sigma, 63069); HEPES (Gibco, 15630080); N2 supplement (Gibco, 17502048); insulin (Sigma, I0516); Hank’s balanced salt solution (HBSS; Gibco, 14175103); papain (Worthington Biochemical Corporation, NC9597281); HyClone™ Cosmic Calf Serum (Cytiva, SH30087.04); sodium pyruvate (Gibco, 11360070); MEM non-essential amino acids solution (NEAA; Gibco, 11140050); 2-mercaptoethanol (Sigma, M3148); Bambanker cryopreservation medium (FUJIFILM Wako, 302-14681); 0.25% trypsin–EDTA (Gibco, 25200056); polyethylenimine (PEI, 25 kDa; Polysciences, 23966); poly-L-ornithine hydrobromide (Sigma, P4957); polybrene (Sigma, TR-1003); mTeSR™ Plus medium (STEMCELL Technologies, 100-0276); CloneR™ supplement (STEMCELL Technologies, 05888).

### Preparation of HeLa nuclear extracts

HeLa cells were cultured in DMEM to confluence, harvested, and washed with ice-cold DPBS. Nuclei were isolated by hypotonic lysis using a Dounce homogenizer (20 strokes) in hypotonic buffer containing: 10 mM HEPES (pH 7.5), 1.5 mM MgCl₂, 10 mM KCl, 0.5 mM DTT, 1 mM PMSF, and a protease inhibitor cocktail. Nuclei were collected by centrifugation and lysed in high-salt buffer containing: 50 mM HEPES (pH 7.5), 10% (v/v) glycerol, 500 mM KCl, 5 mM MgCl₂, 0.1% NP-40, 0.5 mM DTT, 1 mM PMSF, and protease inhibitor cocktail for 30 min at 4 °C with rotation. Lysates were clarified by centrifugation at 13,000 rpm for 10 min at 4 °C, and the resulting supernatant was dialyzed overnight at 4 °C against 2 L of dialysis buffer containing: 50 mM Tris-HCl (pH 8.0), 5% (v/v) glycerol, 150 mM KCl, 2 mM MgCl₂, 0.1% NP-40, and 1 mM DTT. Following dialysis, extracts were transferred to fresh tubes and further clarified by centrifugation at 13,000 rpm for 20 min at 4 °C. The final supernatant was used for peptide pulldown assays.

### H3 peptide pulldown assay

For each pulldown assay, 100 μg of biotinylated histone peptide (H3_1-18_Q5ser *vs.* H3_1-18_ unmodified) was incubated overnight at 4 °C with rotation with 100 μL of prewashed immobilized streptavidin beads (Dynabeads™ M-280 Streptavidin, Invitrogen, 11206D) in DPBS containing 0.01% Triton X-100. For each immunoprecipitation, 40 μL of a 50% peptide–bead slurry was incubated with dialyzed HeLa nuclear extract for 4 h at 4 °C with rotation. Peptide–protein complexes were collected by centrifugation at 1,000 rpm for 1 min, and beads were washed four times with dialysis buffer followed by four washes with detergent-free dialysis buffer (50 mM Tris-HCl (pH 8.0), 150 mM KCl, 2 mM MgCl₂, and 1 mM DTT). For western blot analysis, beads were boiled for 8 min in 30 μL of denaturing sample buffer and proteins were resolved by SDS– PAGE.

### Sample preparation for mass spectrometry

For *liquid chromatography tandem mass spectrometry analysis (LCMS)*, proteins bound to histone peptides were reduced (10mM Dithiothreitol) and alkylated (30mM iodoacetamide) followed by partial trypsinization (10ng/uL, Promega). elution. After 4h at room temperature, the supernatant was transferred to new vial and re-trypsinized overnight. Digestion was halted by addition of 10% trifluoroacetic acid (TFA; Thermo Scientific, 85183 and peptides were solid phase extracted prior to being analyzed by direct loading reversed-phase nano–LC–MS/MS using a Fusion Lumos mass spectrometer operated in high/high acquisition mode. Petptides were separated using a packed-in-emitter 12cm/75um ID/3um C18 column (Nikkyo Technos Co., Ltd. Japan). Peptides were eluted using a gradient delivered at 300nL/min increasing from 2% Buffer B (0.1% formic acid in 80% acetonitrile) / 98% Buffer A (0.1% formic acid) to 35% Buffer B / 65% Buffer A, over 70 minutes (EasyLC 1200, Thermo Scientific). All solvents were LCMS grade (Optima, Fisher Scientific).

### Western blotting

Proteins were electrophoresed on 4–12% NuPAGE Bis-Tris protein gels (Invitrogen) and transferred to nitrocellulose membranes. Efficient transfer was always confirmed with direct blue staining (0.1% stock aqueous solution in milliQ water) before being incubated with primary antibodies overnight at 4 °C (all membranes were blocked in 5% milk or 5% BSA depending on the primary antibody used). Membranes were then washed and incubated with peroxidase-labelled secondary antibodies (1:2,000–1:10,000, depending on the primary antibody used). Bands were detected using SuperSignal™ West Femto Maximum Sensitivity Substrate (Thermo Scientific, 34095) and imaged on a ChemiDoc Imaging System (Bio-Rad) under chemiluminescent detection. Molecular weight markers were visualized using the colorimetric (Ponceau S) imaging setting of the ChemiDoc system. Chemiluminescent images were acquired using fixed exposure settings and are presented in the main figures. Composite images merging chemiluminescent signals and molecular weight marker images were generated for the supplementary materials to display molecular weight references.

### Preparation of SK-N-SH lysates

Harvested SK-N-SH cells were resuspended in hypotonic buffer (10 mM HEPES, pH 7.5, 2 mM MgCl_2_, 0.1% Tween-20, 20% glycerol, 2 mM PMSF, and Roche Complete EDTA-free protease inhibitors) and lysed on ice using a glass Dounce homogenizer (pestle B, 30 strokes). The homogenate was centrifuged at 16,000 × *g* for 15 min at 4 °C. The supernatant was collected and the pellet was resuspended in high-salt buffer (50 mM HEPES, pH 7.5, 420 mM NaCl, 2 mM MgCl_2_, 0.1% Tween-20, 20% glycerol, 2 mM PMSF, and Roche Complete EDTA-free protease inhibitors) and homogenized with a Dounce homogenizer (pestle B, 3 × 30 strokes). The suspension was centrifuged at 16,000 × *g* for 15 min at 4 °C. Supernatants from two portions were pooled and clarified again at 21,000 × *g* for 15 min at 4 °C. Protein concentration was determined using the Bradford assay (Bio-Rad).

### Photo-crosslinking experiments

Photoaffinity probes in the absence or presence of different concentrations of competitors were incubated with recombinant proteins (1 µM) or cell lysate (1.0-1.5 mg/mL) in binding buffer (150 mM NaCl, 50 mM HEPES, pH 7.5, 0.05% Tween 20, 20% glycerol) for 10 min at 4 °C. Then, the samples were irradiated at 365 nm (Spectrolinker XL-1000) for 20 min in 96-well (recombinant proteins, 75 µL each well) or 6-well plate (cell lysate, 1-2 mL each well) on ice.

### Copper(I)-catalyzed azide–alkyne cycloaddition (CuAAC)

To the prepared photo-cross-linking samples, 100 µM rhodamine-N_3_ for in-gel fluorescence (10 mM stock in DMSO) or diazo-biotin-N_3_ for pulldown (5 mM stock in DMSO) was added, followed by 1 mM TCEP (freshly prepared 50 mM stock in H_2_O), 100 µM TBTA (10 mM stock in DMSO), and finally the reactions were initiated by the addition of 1 mM CuSO_4_ (freshly prepared 50 mM stock in H_2_O). The reactions were incubated for 1 h at room temperature with regular vortexing. The reactions were quenched by adding 5 volumes of ice-cold acetone and placed at −20 °C overnight to precipitate proteins.

### In-gel fluorescence scanning

For in-gel fluorescence, acetone-precipitated protein pellets were collected by centrifugation at 6,000 × *g* for 5 min at 4 °C, washed twice with ice-cold methanol, and air-dried for 10 min. Pellets were resuspended in 1 × LDS sample buffer (Invitrogen) containing 50 mM DTT and heated at 95 °C for 10 min before separation by SDS–PAGE. Fluorescently labeled proteins were visualized using an Amersham ImageQuant 800 (excitation 535 nm, emission 580 nm).

### Enrichment of biotinylated proteins

For enrichment of biotinylated proteins, acetone-precipitated pellets were centrifuged at 3,500 × *g* for 5 min at 4 °C, washed twice with ice-cold methanol, and air-dried for 10 min. Protein pellets were dissolved in PBS containing 4% SDS, 20 mM EDTA, and 10% glycerol by vortexing and heating, then diluted with 1 × PBS to a final SDS concentration of 0.5%. High-capacity streptavidin agarose beads (Thermo Fisher Scientific) were added, and samples were rotated for 1.5 h at room temperature to capture biotinylated proteins. The beads were stepwise washed by 1 × PBS with 0.2% SDS (5 times), 6 M urea in 1 × PBS with 0.1% SDS (5 times) and 250 mM NH_4_HCO_3_ with 0.05% SDS (5 times). The enriched proteins were eluted by incubating the beads with 25 mM Na_2_S_2_O_4_, 250 mM NH_4_HCO_3_, and 0.05% SDS for 1 h. The eluted proteins were dried with SpeedVac.

For mass spectrometry samples, before washing by 250 mM NH_4_HCO_3_ with 0.05% SDS, beads were incubated in 10 mM DTT, 6 M urea in 1 × PBS with 0.1% SDS at 37 °C for 1 h and then were added 30 mM iodoacetamide at room temperature for another 30 min in dark to reduce and alkylate cysteines.

### TMT-based quantitative proteomics

SpeedVac samples were redissolved in 100 μL water. Proteins were precipitated by sequential addition of 400 μL methanol, 100 μL chloroform, and 300 μL water, followed by centrifugation at 16,000 × *g* for 2 min; the upper aqueous phase was removed, and the protein interphase was washed with 600 μL pre-chilled (−20 °C) methanol and centrifuged again under the same conditions. The resulting protein pellets were air-dried and reconstituted in 50 mM triethylammonium bicarbonate (TEAB, pH 8.0), then digested with 0.2 μg MS-grade trypsin at 37 °C for 12 h with shaking, followed by an addition of 0.1 μg trypsin for 2 h to ensure complete digestion. Peptide concentrations were measured using the Pierce Quantitative Fluorometric Peptide Assay (Thermo Fisher Scientific), and peptides in 80 μL (∼4 mg) of 50 mM TEAB were labeled with TMT reagents (Thermo Fisher Scientific) dissolved in 20 μL acetonitrile for 1 h at room temperature. Labeling reactions were quenched with 4 μL of 5% hydroxylamine for 15 min at room temperature, after which labeled peptides were combined and dried in SpeedVac, followed by desalting using Sep-Pak Vac tC18 cartridges (Waters), and dried again. Peptides were fractionated on an XBridge Peptide BEH C18 column (300 Å, 3.5 μm, 2.1 mm × 250 mm) using a 2–80% acetonitrile gradient in 10 mM ammonium bicarbonate (pH 10.0) over 50 min, dried, and reconstituted in 0.1% formic acid for LC-MS/MS.

### Binding-site mapping sample preparation for QR2

For binding-site mapping, recombinant QR2 (1 μM) was incubated with probes 2 and 3 (5 μM each) in 3 mL of binding buffer (50 mM HEPES, 150 mM NaCl, 2 mM MgCl_2_, 0.1% Tween-20, 20% glycerol, pH 7.5) for 10 min at 4 °C, followed by UV irradiation at 365 nm for 20 min. After crosslinking, acid-cleavable DADPS biotin-azide (100 μM) was added together with TCEP (1 mM) and TBTA (100 μM), and CuAAC was initiated by adding 1 mM CuSO_4_ and incubating for 1.5 h at room temperature before quenching with four volumes of ice-cold acetone and overnight precipitation at −20 °C. Pellets were collected by centrifugation (3,500 × *g*, 5 min, 4 °C), washed twice with ice-cold methanol, air-dried, and dissolved in 1 × PBS containing 4% SDS, 20 mM EDTA, and 10% glycerol, then diluted to 0.5% SDS and incubated with high-capacity streptavidin agarose beads (Thermo Fisher Scientific) for 1.5 h at room temperature. Beads were washed sequentially with PBS + 0.2% SDS, PBS + 0.1% SDS + 6 M urea, and resuspended in 6 M urea in 100 mM NH_4_HCO_3_. Then, the samples were reduced with DTT (10 mM, 1 h, room temperature), alkylated with iodoacetamide (30 mM, 30 min, dark, room temperature), followed by washed and resuspended in 1 M urea in 100 mM NH_4_HCO_3_ for on-bead digestion with trypsin (1 μg, 16 h, 37 °C). SC-OH–labeled peptides were eluted twice with 100 μL 2% formic acid (30 min each), followed by two washes with 50% acetonitrile + 1% formic acid. All eluates and washes were pooled, dried in a SpeedVac, resuspended in 200 μL 0.5% acetic acid, and desalted using StageTips for LC-MS/MS.

### LC–MS/MS analysis and data analysis

The TMT samples were analyzed on an Orbitrap Ascend Tribrid mass spectrometer coupled to a Vanquish™ Neo UHPLC system (Thermo Fisher Scientific). Peptides were first loaded onto a PepMap 100 C18 trapping cartridge (5 µm, 300 µm × 5 mm) and subsequently separated on a PepMap RSLC C18 analytical column (2 µm, 75 µm × 50 cm). Data acquisition was performed using Thermo Xcalibur 4.3 software. Electrospray ionization was achieved by applying a voltage of 1,800 V through the PEEK junction at the inlet of the microcapillary column. Peptides were separated with a 4–40% acetonitrile gradient in 0.1% formic acid over 140 min at a flow rate of 250 nL/min. Data were acquired in data-dependent acquisition (DDA) mode optimized for TMT-based quantitation. Full MS1 scans were collected in the Orbitrap at a resolution of 120,000 over an m/z range of 400–1,600 Th, with an automatic gain control (AGC) target of 4E5 and a maximum injection time (IT) of 50 ms. The most intense precursor ions above the defined intensity threshold were sequentially selected for MS2 fragmentation using collision-induced dissociation (CID) with a collision energy of 30%, activation time of 10 ms, and an isolation width of 0.7 m/z. MS2 spectra were acquired in the ion trap at turbo scan rate, with an AGC target of 1E4 and IT of 35 ms. For TMT reporter ion quantitation, synchronous precursor selection (SPS) MS3 scans were performed on up to 20 MS2 fragment ions using higher-energy collisional dissociation (HCD) with a normalized collision energy (NCE) of 55%. MS3 spectra were acquired in the Orbitrap at a resolution of 30,000, with an AGC target of 1E5 and IT of 200 ms. Dynamic exclusion was set to 60 s to prevent repeated analysis of the same precursor ions, and charge states of 2–6 were considered for fragmentation. Proteome Discoverer (v3.0) was used to search the raw data against the UniProt *Homo sapiens* (UP000005640). Fixed modifications included cysteine carbamidomethylation and TMT-tag labeling on lysine side chains and peptide N-termini. Variable modifications included methionine oxidation, protein N-terminal acetylation, and methionine loss. Mass error tolerances were set to 20 ppm for MS1 spectra, 0.1 Da for MS2 spectra, and 20 ppm for MS3 spectra. A maximum of two missed cleavages was allowed. The false discovery rate (FDR) for both peptides and proteins was set at 0.01. Missing values were imputed using the low-abundance sampling method. Quantification was based on five biological replicates.

Binding-site mapping samples were analyzed on a Q Exactive PLUS mass spectrometer (Thermo Fisher Scientific) coupled to an Ultimate 3000 RSLCnano system. Peptides were first loaded onto a PepMap 100 C18 trapping cartridge (5 μm, 300 μm × 5 mm) and then separated on a PepMap RSLC C18 analytical column (2 μm, 75 μm × 25 cm). Data acquisition was performed using Thermo Xcalibur v4.3 software. Peptides were eluted with a 4–40% acetonitrile gradient in 0.1% formic acid over 90 min at a flow rate of 300 nL/min. Electrospray ionization was achieved by applying 1,800 V through a PEEK junction at the column inlet.

MS acquisition was conducted in data-dependent acquisition mode. Survey scans (m/z 300–1,800) were acquired in the Orbitrap at a resolution of 70,000, followed by HCD fragmentation of the top 20 most intense precursors (isolation window ±1.5 m/z, resolution 17,500). Automatic gain control (AGC) targets were set to 1 × 10^6^ ions for MS1 and 5 × 10^4^ ions for MS2, with maximum ion injection times of 30 ms and 50 ms, respectively. The normalized collision energy for HCD was 27%. Singly charged ions, ions with >8 charges, and unassigned charge states were excluded from MS2 selection.

### Vector construction and recombinant QR2 protein purification

Human *NQO2* cDNA was initially cloned into the pUC19 vector. Site-directed mutagenesis was performed to generate the I129E/I195R *NQO2* mutant. All constructs were verified by Sanger sequencing. Wildtype and mutant *NQO2* inserts were subsequently subcloned into the pET28a expression vector (Addgene #69864) to generate pET28a–*NQO2*, pET28a–*NQO2*(I129EI195R) and pET28a–*NQO2*(N162E) respectively. Recombinant wildtype and mutant QR2 proteins were expressed in *Escherichia coli* BL21 cells. Protein expression was induced with 0.5 mM isopropyl β-D-1-thiogalactopyranoside (IPTG) at 16 °C overnight in LB medium. Cells were harvested and resuspended in lysis buffer containing: 500 mM NaCl, 20 mM Tris-HCl (pH 7.5), and 10% (v/v) glycerol. Cells were lysed and clarified by centrifugation, and the supernatant was applied to a HisTrap affinity column (GE Healthcare). The column was washed with five column volumes of lysis buffer, and bound proteins were eluted with buffer containing: 150 mM NaCl, 20 mM Tris-HCl (pH 7.5), and 500 mM imidazole. Purified proteins were buffer-exchanged into storage buffer (150 mM NaCl, 20 mM Tris-HCl, pH 7.5), concentrated to approximately 10 mg mL⁻¹, aliquoted, and stored at −80 °C.

### Vector construction and recombinant QR2 purification

Human NQO2 cDNA was first cloned into pUC19, and site-directed mutagenesis was used to generate the I129EI195R and N162E mutants; all constructs were verified by Sanger sequencing. Wildtype and mutant inserts were then subcloned into pET28a (Addgene #69864) to give pET28a– NQO2, pET28a–NQO2(I129EI195R) and pET28a–NQO2(N162E). Recombinant wildtype and mutant QR2 were expressed in *Escherichia coli* BL21 (DE3), induced with 0.5 mM isopropyl β-D-1-thiogalactopyranoside (IPTG) at 16 °C overnight in LB medium. Cells were harvested, resuspended in lysis buffer (20 mM Tris-HCl, pH 7.5, 500 mM NaCl, 10% (v/v) glycerol), lysed and clarified by centrifugation, and the supernatant was applied to a HisTrap column (GE Healthcare). The column was washed with five column volumes of lysis buffer and bound protein eluted in 20 mM Tris-HCl, pH 7.5, 150 mM NaCl, 500 mM imidazole. Purified protein was buffer-exchanged into storage buffer (20 mM Tris-HCl, pH 7.5, 150 mM NaCl), concentrated to approximately 10 mg mL⁻¹, aliquoted and stored at −80 °C.

### QR1 and QR2 expression and purification

Plasmids encoding human QR1 and QR2 with an N-terminal hexahistidine (6×His) tag in pProEX-HTa were a gift from R. M. Vabulas. Point mutations were introduced with the Mut Express Universal Fast Mutagenesis Kit (Vazyme) and verified by Sanger sequencing. Full-length proteins were expressed in *Escherichia coli* Rosetta (DE3) in LB medium, induced with 0.2 mM IPTG at 16 °C overnight. Cells were lysed by sonication or French press in lysis buffer (1× PBS, pH 7.4, 20 mM imidazole, 1 mM PMSF, 0.5× Roche cOmplete EDTA-free protease inhibitor cocktail), and lysates were clarified at 25,000 × g for 30 min at 4 °C. The supernatant was bound to Ni–NTA resin for 2 h at 4 °C and washed with 1× PBS followed by wash buffer (1× PBS, pH 7.4, 20 mM imidazole). Bound flavin was stripped by washing and incubating the resin in denaturing buffer (1× PBS, pH 7.4, 2 M urea, 2 M KBr), repeated until no FAD was detectable in the supernatant by fluorescence (excitation 450 nm, emission 530 nm), then reloaded by overnight incubation in 1× PBS, pH 7.4, containing 20 mM FAD. Protein was eluted in 1× PBS, pH 7.4, 500 mM imidazole and polished by size-exclusion chromatography on a Superdex 75 column (Cytiva) in 50 mM HEPES, pH 7.4, 150 mM NaCl. QR2 mutants were purified identically.

### Protein preparation for X-ray crystallography and ITC

Human *NQO2* was subcloned into the pTrcHis2C expression vector (Thermo Fisher Scientific) to generate a construct encoding a C-terminal hexahistidine (6×His) tag. Point mutations in *NQO2* were introduced using QuikChange site-directed mutagenesis and verified by Sanger sequencing. Full-length QR2 proteins were expressed in *Escherichia coli* BL21 (DE3) cells. Protein expression was induced with 0.2 mM isopropyl β-D-1-thiogalactopyranoside (IPTG) at 16 °C overnight in LB medium. Cells were harvested by centrifugation, resuspended in buffer A (150 mM NaCl, 20 mM Tris-HCl, pH 7.5, 10 mM imidazole), and lysed using an EmulsiFlex-C3 high-pressure homogenizer (Avestin). Clarified lysates were incubated with Ni–NTA affinity resin for purification.

To remove bound flavin cofactors, the resin-bound protein was washed with buffer B (buffer A supplemented with 2 M KBr and 2 M urea), followed by overnight incubation in buffer A containing 10 mM flavin adenine dinucleotide (FAD) to reload the cofactor. Eluted protein was further purified by anion-exchange chromatography using a Q HP column (Cytiva) and size-exclusion chromatography on a Superdex 75 column (Cytiva). All QR2 mutant proteins were purified using the same procedure. H3 peptides were prepared as previously described^13^ or synthesized commercially (Beijing SciLight Biotechnology).

### Isothermal titration calorimetry

Isothermal titration calorimetry (ITC) experiments were performed at 25 °C using a MicroCal PEAQ-ITC instrument (Malvern Instruments). The sample cell was loaded with 200 μL of QR2 protein (100 μM), and titrated with 17 sequential injections of histone peptide or serotonin (5-HT; MedChemExpress) at a concentration of 1 mM. Titration data were analyzed using Origin 7.0 software and fitted to a single-site binding model. ITC statistics are provided in **Table S1**.

### Crystallization, data collection, and structure determination

Crystallization trials were performed using the sitting-drop vapor diffusion method at 18 °C. Purified QR2 protein (10 mg mL⁻¹) was preincubated with H3Q5ser or H3K4me3Q5ser peptides at a 1:10 molar ratio in buffer containing: 100 mM NaCl and 20 mM Tris-HCl (pH 7.5). Crystals of the QR2–H3Q5ser complex were obtained under multiple reservoir conditions, including mixtures containing polyols, imidazole/MES buffers, hexylene glycol, and polyethylene glycol (PEG) 1000 and PEG 3350 at pH 6.5. Crystals of the QR2–H3K4me3Q5ser complex were obtained under similar conditions, using alternative salt components.

Crystals were cryoprotected and flash-frozen in liquid nitrogen prior to data collection. X-ray diffraction data were collected at the Shanghai Synchrotron Radiation Facility. Diffraction images were indexed, integrated, and scaled using HKL2000. Structures were solved by molecular replacement using MOLREP^69^ from the CCP4 suite, with the QR2 structure (PDB ID: 1QR2) as the search model. Model building and refinement were carried out using COOT^70^ and PHENIX^71^, respectively. Data collection and refinement statistics are provided in **Table S2**.

### Generation of unmodified H3 and H3Q5ser nucleosomes

#### Peptide synthesis

Synthetic peptide α-thioesters used for semisynthesis were manually synthesized on Trityl-OH resin (ChemMatrix). The resin was chlorinated in dry DCM with 12 equivalents of thionyl chloride and 24 equivalents of pyridine (2 x 2 h). Excess thionyl chloride was quenched and washed with a 10% DIPEA/DCM solution. Following the chlorination, hydrazine loading was performed in DMF using 10 equivalents of Fmoc-carbazate, and 10 equivalents of DIPEA (2 x 2 h). Unreacted trityl-chloride resin was then capped with 50 equivalents of methanol in DMF (30 min) and washed extensively. Subsequent peptide synthesis was carried out through iterative cycles of Fmoc deprotection (20% piperidine in DMF with 0.1 M HOBt; 1 x 1 min and 1 x 20 min) and amino acid coupling (5 equivalents of standard Fmoc-protected amino acid, 4.9 equivalents of HATU, and 10 equivalents of DIPEA for 2 × 30 min), with extensive DMF washes between steps. Double couplings were performed when necessary to ensure complete acylation. Serotonylated glutamine residues were introduced by incorporation of Fmoc-L-glutamic acid γ-allyl ester (Chem-Impex; Coupling condition: 2 equivalents of Fmoc-L-glutamic acid γ-allyl ester, 1.9 equivalents of HATU, and 5 equivalents of DIPEA for 2 x 2 h), followed by allyl deprotection and serotonin coupling as previously described^14^. Completed peptides were cleaved and deprotected using 92.5% trifluoroacetic acid (TFA), 2.5% ethanedithiol, 2.5% triisopropylsilane, and 2.5% water, precipitated with diethyl ether, and air-dried. Peptide hydrazides were converted to C-terminal thioesters via the peptide acyl pyrazole intermediates following established protocols^72^. Briefly, 1 mM crude peptide hydrazides were dissolved in 100 mM phosphate buffer (pH = 3) with 6 M guanidinium hydrochloride and 2.5 mM acetylacetone. Excess 4-mercaptophenylacetic acid was then added to the solution, and the suspensions were vigorously shaken overnight at room temperature. The reaction mixtures were centrifuged, and the supernatant containing the products were filtered. The peptide MPAA thioesters were purified on preparative reversed-phase HPLC using a 0–30% solvent B (10% water, 90% acetonitrile with 0.1% TFA) gradient over 60 min.

#### Preparation of recombinant histones

Recombinant human histones H2A, H2B, and H4 were expressed in *E. coli* BL21(DE3) cells and purified with minor modifications to established protocols. Cultures were grown in LB medium at 37 °C to an OD₆₀₀ of ∼0.6 and induced with 1 mM IPTG for 3–4 h. Cells were harvested, resuspended in lysis buffer (20 mM Tris-HCl pH 7.5, 200 mM NaCl, 1 mM EDTA, 1 mM β-mercaptoethanol, 0.1% Triton X-100), and lysed by sonication.

Inclusion bodies were collected by centrifugation, washed twice in lysis buffer lacking detergent, and extracted in resuspension buffer (7 M urea, 10 mM Tris-HCl, 1 mM EDTA, 100 mM NaCl, 1 mM DTT). Clarified extracts were purified by cation-exchange chromatography using a HiTrap SP FF column with a linear NaCl gradient. Histone-containing fractions were pooled and further purified by preparative reversed-phase HPLC.

#### Preparation of truncated histone H3

N-terminally His₆–SUMO–tagged human histone H3(14–135, K14C) was expressed in *E. coli* BL21(DE3). Cells were lysed under denaturing conditions (6 M guanidinium hydrochloride), and the tagged protein was purified by Ni–NTA affinity chromatography. Eluted protein was stepwise dialyzed to remove denaturant and refold the protein. Ulp1 protease was added to cleave the His₆–SUMO tag. Following proteolysis, solubilized truncated histone was purified by preparative reversed-phase HPLC.

#### Semisynthesis of modified histone H3

Full-length histone H3 was generated by native chemical ligation between peptide α-thioesters corresponding to residues 1–13 and truncated H3(14–135, K14C). Ligation reactions were performed in a well-ventilated chemical hood in 200 mM sodium phosphate buffer (pH = 7.0–7.2) containing 6 M guanidinium hydrochloride, 50 mM TCEP, and 2% v/v TFET at room temperature overnight. Completion of ligation was confirmed by HPLC and ESI–MS. The ligation product was purified by preparative reversed-phase HPLC and treated with 2-bromoethylammonium bromide to convert the non-native cysteine at position 14 into a lysine analogue following established protocols^73^. Final products were purified by semi-preparative HPLC and validated by HPLC and ESI–MS.

#### Histone octamer refolding

Histone octamers were assembled using established protocols.^1^ Individual histones were dissolved in unfolding buffer (6 M guanidinium chloride, 20 mM Tris-HCl pH 7.9, 1 mM DTT), quantified by absorbance at 280 nm, and combined at a molar ratio of H3:H4:H2A:H2B = 1:1:1.1:1.1 to a final concentration of 1 mg mL⁻¹. The mixture was dialyzed stepwise into refolding buffer (2 M NaCl, 10 mM Tris-HCl pH 7.9, 1 mM EDTA, 1 mM DTT) at 4 °C. Refolded octamers were purified by size-exclusion chromatography using a Superdex S200 Increase column. Pure fractions were pooled, adjusted to 50% glycerol, and stored at −20 °C.

#### DNA preparation

Biotinylated Widom 601 DNA containing 15-bp overhangs was prepared for mononucleosome assembly. The target fragment was amplified by PCR with Pfu DNA polymerase. A 5′-biotinylated forward primer (5’-Biotin-TACTACGCGGCCGCCCTGG-3’) and an unmodified reverse primer (5’-CTGGATCTTACATGCACAGGATGTATATATCTGACACGTGC-3’) were synthesized and HPLC-purified commercially (IDT). The PCR product was purified by ethanol precipitation. Briefly, sodium acetate (0.1 volumes of 3 M solution, pH 5.3) and ethanol (2.5 volumes) were added to the reaction mixture, and the DNA product was precipitated overnight at −20 °C. Precipitated DNA was collected by centrifugation and resuspended in a minimal volume of nuclease-free water. The sample was further purified using PCR purification silica columns according to the manufacturer’s instructions and eluted in nuclease-free water. DNA concentration was determined by UV absorbance using a NanoDrop spectrophotometer.

#### Nucleosome reconstitution

Nucleosomes were assembled by salt-gradient dialysis. Histone octamers were mixed with 601 DNA in octamer refolding buffer (10 mM Tris-HCl, 2 M NaCl, 0.5 mM EDTA, 1 mM DTT, pH 7.8) and dialyzed stepwise at 4 °C to decreasing the salt concentrations (from 2 M NaCl to 1.4 M KCl, and then to 10 mM KCl overnight via a peristaltic pump) using a Slide-A-Lyzer MINI dialysis device (3.5-kDa cutoff, Thermo Scientific). Following overnight dialysis, the samples were incubated at 37°C for 30 minutes to promote nucleosome repositioning and subjected to two final rounds of dialysis in nucleosome assembly end buffer. Samples were clarified by centrifugation, and nucleosome concentrations were determined by absorbance at 260 nm. Nucleosome quality was assessed by native polyacrylamide gel electrophoresis and ethidium bromide staining.

#### In vitro pulldown assays

*In vitro pulldown assays* were performed using either biotinylated mononucleosomes or biotinylated histone peptides as binding substrates. For each assay, 1 μg of biotinylated substrate was preincubated with 10 μL of streptavidin Dynabeads™ M-280 at 4 °C to allow immobilization. Substrate–bead complexes were then incubated with 1 μg of purified recombinant proteins in 200 μL of binding buffer containing: 150 mM NaCl, 20 mM Tris-HCl (pH 7.5), and 0.1% NP-40 for 1 h at 4 °C with rotation. Following incubation, beads were washed three times with wash buffer containing: 350 mM NaCl, 20 mM Tris-HCl (pH 7.5), and 1% NP-40. Bound proteins were eluted by boiling beads in 30 μL of denaturing sample buffer and analyzed by western blotting.

#### Flag–HA–QR2 immunoprecipitation and mass spectrometry

Flag–HA mock control, Flag–HA–tagged wildtype QR2, or Flag–HA–tagged QR2 I129EI195R mutant HeLa cell lines were cultured in DMEM to confluence. Dialyzed nuclear extracts were prepared as described above.

Flag M2 agarose beads were prewashed with 100 mM glycine (pH 2.7) and equilibrated in binding buffer (pH 7.5). For each immunoprecipitation, 50 μL of Flag M2 agarose beads were incubated with nuclear extract for 4 h at 4 °C with rotation. Beads were washed four times with dialysis buffer followed by four washes with detergent-free dialysis buffer. Bound proteins were processed for mass spectrometry using the same protocol described above.

#### Mass spectrometry data processing and analysis

Mass spectrometry data were searched against the UniProt human protein database using MaxQuant (v. 2.0.3.0 or v. 2.4.2.0)^74^ using full tryptic constraints. Oxidation of methionine residues and protein N-terminal acetylation were specified as variable modifications, and carbamidomethylation of cysteine residues was set as a fixed modification. False Discover Rate (FDR) were set at 1% for both peptides and proteins. The experiment comprised two peptide conditions (H3_1-18_Q5ser *vs.* H3_1-18_ unmodified), each analyzed with four biological replicates. Quantitative data were processed using Perseus v. 2.0.10.0^74^. Reverse database hits and potential contaminants were removed prior to analysis. Log₂-transformed intensity-based absolute quantification (iBAQ) values were median-normalized on a per-sample basis. Proteins were required to be quantified in at least three replicates in at least one condition to be included in downstream analyses. Missing values were imputed using a normal distribution (width = 0.3; downshift = 1.8). Statistical comparisons between unmodified H3 and H3Q5ser conditions were performed using a permutation-based two-sided t-test with an FDR threshold of 0.05.

Raw data were processed using Proteome Discoverer v3.0, with SC-OH (exact mass: 326.1777 Da) specified as a variable modification on any residue. The NQO2 single-protein FASTA file was used for database searching. Default parameters were applied for identification and quantification: parent MS tolerance of 20 ppm, MS/MS tolerance of 20 ppm, minimum peptide length of 6 amino acids, and up to 2 missed cleavages. Peptide quantification was based on peptide-spectrum match (PSM) counts.

#### BNAH fluorescence-based enzymatic activity assay

QR2 enzymatic activity was measured using a fluorescence-based BNAH oxidation assay. Oxidation of the reduced co-substrate 1-benzyl-1,4-dihydronicotinamide (BNAH) (Santa Cruz, sc-208609) by QR2 results in a time-dependent decrease in intrinsic BNAH fluorescence. Reactions were performed in assay buffer containing: 50 mM Tris-HCl (pH 8.0) and 0.1% Triton X-100, supplemented with 100 µM Menadione (K3, Sigma, 47775) and BNAH. Reactions were initiated by addition of recombinant QR2 (100 ng per reaction). Control reactions lacking enzyme were included to account for non-enzymatic BNAH decay. Fluorescence was monitored every 2 min using a SpectraMax iD5 plate reader with excitation at 340–355 nm and emission at 440–460 nm. Enzymatic activity was calculated from the linear decrease in fluorescence intensity over time.

#### In vitro erasure-protection assay

Recombinant histone H3 protein carrying the Q5ser modification (2 µM) was first incubated with QR2 (0–50 µM) in buffer containing 25 mM Tris-HCl (pH 7.5), 10 mM CaCl₂, 10 mM KCl and 100 mM NaCl at 37 °C. After 5 min, TGM2 (0.2 µM) was added, and the reaction was continued for 1 h at 37 °C. The reaction was quenched by the addition of LDS loading buffer, followed by boiling for 10 min and immunoblotting analysis. Band intensities were quantified by densitometry, and data are shown as the mean of three independent experiments (n = 3). Group differences were analyzed by ordinary one-way ANOVA followed by Dunnett’s multiple-comparisons test (alpha = 0.05), with adjusted P values reported. Significance was defined as *P < 0.05, **P < 0.01 and ***P < 0.001; ns, P > 0.05.

#### Comparative analysis of NQO2 orthologs

NQO2 orthologous protein sequences from 16 vertebrate species, spanning sea lamprey (*Petromyzon marinus*) to human (*Homo sapiens*), were obtained from NCBI RefSeq / Ensembl (**Data S3**). Pairwise amino-acid percent identity was computed for each species pair over alignment columns at which both sequences carried a residue. Per-position conservation was assessed across all 16 orthologues, a position being scored invariant when all non-gap residues were identical (32% of positions; 75 of 231). The focal residues (human NQO2 positions 129, 162, and 195) were mapped to alignment columns through the human sequence and classified as identical to the human residue, of the same physicochemical class, or of a different class, with hydrophobic and aromatic residues grouped as a single class because the constraint at these positions is side-chain packing. A maximum-likelihood phylogeny was inferred with phangorn under the LG+Γ model (four rate categories; α = 0.74), optimizing branch lengths on the accepted species topology (held fixed, as a single ∼230-residue protein provides insufficient signal to resolve the shallow nodes) and rooting on *P. marinus*. Conservation analysis and visualization were performed in R v4.4.3 (Biostrings, msa, ape, phangorn, reshape2, patchwork, ggplot2).

#### Lentivirus packaging

HEK293T cells were maintained in DMEM supplemented with 10% fetal bovine serum (FBS), 100 U mL⁻¹ penicillin, and 100 μg mL⁻¹ streptomycin. One day prior to transfection, 10-cm culture dishes were coated with 10 mL of poly-L-ornithine, incubated at 37 °C for 1 h, and washed twice with DPBS. HEK293T cells were dissociated using 0.25% trypsin–EDTA and seeded at a density of 7 × 10⁶ cells per dish in 6 mL of DMEM. After 24 h, the medium was replaced with fresh DMEM and cells were transfected using a polyethylenimine (PEI)-based method. For each dish, a total of 20 μg plasmid DNA (10 μg lentiviral transfer plasmid, 5 μg pMDLg/pRRE, 2.5 μg RSV-Rev, and 2.5 μg pCMV-VSV-G) was diluted in 500 μL DMEM. In parallel, 60 μL of PEI (1 mg mL⁻¹) was diluted in 500 μL DMEM. DNA and PEI solutions were combined, incubated for 20 min at room temperature to allow complex formation, and added dropwise to the cultures. Six hours after transfection, cells were washed once with DPBS and fresh, prewarmed DMEM was added. Viral supernatants were collected at 30 and 46 h post-transfection, filtered through a 0.45-μm membrane, and concentrated by ultracentrifugation at 21,000 rpm for 2 h at 4 °C. Viral pellets were resuspended overnight at 4 °C in DMEM/F12 supplemented with HEPES at a volume 100-fold smaller than the original supernatant. An equal volume of 1 M sucrose in DMEM/F12 was then added, and viral stocks were aliquoted and stored at −80 °C.

#### Knockout and rescue of QR2 expression in HeLa cells

*NQO2* knockout HeLa cell lines were generated using CRISPR–Cas9–mediated genome editing. Alt-R™ S.p. HiFi Cas9 Nuclease V3 (IDT, 1081061) was complexed with an Alt-R CRISPR– Cas9 single-guide RNA (sgRNA) targeting human *NQO2* (5′-CCCACGAAGCCTACAAGCAA-3′; IDT) and delivered into HeLa cells using the SE Cell Line 4D-Nucleofector™ X Kit S (Lonza, V4XC-1032) according to the manufacturer’s instructions. Following nucleofection, single-cell clones were isolated and expanded, and *NQO2* knockout was confirmed by western blot analysis. For rescue experiments, the lentiviral backbone pLV-EF1α-IRES-Puro (Addgene #85132) was used to generate expression constructs encoding either Flag–HA–tagged wild-type *NQO2* or the I129EI195R *NQO2* mutant under the control of the endogenous *NQO2* promoter. Lentiviral particles were produced as described above and used to transduce *NQO2*-knockout HeLa cells in DMEM containing 8 μg mL⁻¹ polybrene. Cells were incubated at 37 °C with 5% CO₂ for 24 h, then passaged and selected in DMEM containing 2.0 µg mL⁻¹ puromycin. Selection medium was replaced every 3–4 days until resistant colonies were established. Re-expression of wildtype or mutant QR2 was verified by western blot analysis.

#### Cerebellar granule neuron culture and depolarization

Cerebellar granule neurons were prepared from postnatal day 7 (P7) CD-1 mouse pups as previously described^75^. Briefly, neurons were cultured *in vitro* in high-potassium medium during days in vitro (DIV) 1–2 consisting of Basal Medium Eagle supplemented with 5% HyClone bovine growth serum, 1× penicillin–streptomycin, 1× GlutaMAX and 25 mM KCl. On DIV 3, the medium was replaced with low-potassium medium containing Basal Medium Eagle supplemented with 5% HyClone bovine growth serum, 1× penicillin–streptomycin, 1× GlutaMAX, and 5 mM KCl. On DIV 5, neurons were pretreated with 1 μM TTX and 100 μM DL-AP5 for 12 h to suppress spontaneous neuronal activity. For depolarization, cultures were switched to high-potassium medium (25 mM KCl) for 1 h, after which cells were collected for downstream analyses.

#### Mouse glial cell preparation

Primary mouse glial cultures were prepared from CD-1 pups (postnatal days P1–P3) in accordance with institutional animal care and use guidelines. Pups were anesthetized by hypothermia and humanely euthanized on ice. Cerebral cortices were dissected in ice-cold HBSS supplemented with 10 mM HEPES, and meninges were carefully removed. Cortical tissue was enzymatically dissociated by incubation with papain (80 µL), activated with 0.5 µM CaCl₂ and 1 µM EDTA, at 37 °C for 15 min. Following digestion, tissue was gently triturated and passed through a 70 µm cell strainer. Cells were pelleted by centrifugation at 500g for 5 min and resuspended in mouse embryonic fibroblast (MEF) medium consisting of 435 mL DMEM, 50 mL HyClone™ Cosmic Calf Serum, 5 mL penicillin–streptomycin, 5 mL sodium pyruvate, 5 mL non-essential amino acids (NEAA), and 4 µL 2-mercaptoethanol. Cells were plated onto 10 cm tissue culture dishes, and the medium was replaced after 24 h. After 5–6 days in culture, glial cells were detached, resuspended in Bambanker freezing medium, and cryopreserved in liquid nitrogen until use.

#### One-step neuronal induction via forced NGN2 expression

Glutamatergic neurons were generated by doxycycline-inducible overexpression of the transcription factor NGN2, as previously described^50^. PiggyBac–NGN2–integrated WTC11 hPSCs were dissociated using Accutase for 5 min at 37 °C and resuspended in N3 induction medium supplemented with Chroman1 and doxycycline (Dox). N3 induction medium was prepared by combining 485 mL DMEM/F-12, 5 mL N2 supplement, 5 mL non-essential amino acids, 5 mL penicillin–streptomycin, and 10 mg insulin. Cells were counted and seeded at a density of 3 × 10⁶ cells per Geltrex-coated 10-cm dish. Approximately 16–18 h after plating (day 1), the medium was replaced with fresh N3 medium containing Dox. On day 5, the medium was replaced with N3 medium supplemented with Dox and 4 µM cytosine β-D-arabinofuranoside (Ara-C). On day 7, cells were dissociated by incubation with Accutase for 20 min at 37 °C and resuspended in Neurobasal-zero medium supplemented with Chroman1. Neurobasal-zero medium was prepared by combining 483 mL Neurobasal medium, 5 mL GlutaMAX, 10 mL B27 supplement, and 2.5 mL penicillin–streptomycin. Cells were then co-cultured with mouse glial cells in NBP medium and plated onto pre-coated six-well plates. NBP medium consisted of Neurobasal-zero supplemented with 5% fetal bovine serum (FBS), Chroman1, and Dox. On day 8, the medium was replaced with Neurobasal-zero containing 2% FBS (NB-2%). On day 14, cultures were transitioned to Neurobasal-A-zero medium containing 1% FBS (NBA-1%). Neurobasal-A-zero medium was prepared by combining 483 mL Neurobasal-A, 5 mL GlutaMAX, 10 mL B27 supplement, and 2.5 mL penicillin–streptomycin. From day 14 onward, 30% of the medium was replaced twice weekly until day 35, at which time neurons were used for functional and molecular analyses.

#### Time-course of neuronal differentiation

For time-course analyses of neuronal differentiation, samples were collected at six developmental stages: undifferentiated hPSCs (day 0), and induced neurons at days 6, 12, 20, 27, and 35 for RNA-sequencing. CUT&RUN profiling was performed on undifferentiated hPSCs (day 0) and mature induced neurons at day 35.

#### Depolarization treatment

For membrane depolarization experiments, neurons were pretreated with 1 µM TTX and 100 µM DL-AP5 for 12 h to suppress spontaneous neuronal activity. Depolarization was then induced by the addition of a high-potassium buffer (170 mM KCl, 2 mM CaCl₂, 1 mM MgCl₂, and 10 mM HEPES) to achieve a final extracellular KCl concentration of 55 mM, as previously described^76^. Untreated control neurons and neurons subjected to 1 h of depolarization were harvested for RNA-sequencing.

#### Immunocytochemistry (ICC)

Cells were washed with DPBS and fixed with 4% paraformaldehyde (PFA) for 5 min at room temperature. Following three washes with DPBS, cells were permeabilized with 0.2% Triton X-100 for 5–10 min at room temperature. Non-specific binding was blocked for 2 h at room temperature in blocking buffer consisting of DPBS supplemented with 4% bovine serum albumin (BSA; Sigma, A9647), 1% fetal bovine serum (FBS), and 0.02% sodium azide (Sigma, 71289). For human iPSC-derived neurons, cells were incubated overnight at 4 °C with primary antibodies diluted in blocking buffer, including anti-MAP2 (1:20,000), anti-Synapsin I/II (1:500), and anti-PSD95 (1:500). For cultured mouse cerebellar granule neurons, cells were incubated overnight at 4 °C with anti-NQO2 (1:1,000) and anti-H3K4me3Q5ser (1:1,000). For mouse brain slices, tissues were incubated with anti-NQO2 under the same conditions. After three washes with DPBS, cells or tissue sections were incubated for 1 h at room temperature in the dark with species-specific secondary antibodies diluted in blocking buffer: Alexa Fluor 488–conjugated anti-chicken IgY (H+L), Alexa Fluor 555–conjugated anti-rabbit IgG (H+L), and Alexa Fluor 647–conjugated anti– guinea pig IgG (H+L) (all at 1:1,000). Samples were washed three times with DPBS, and nuclei were counterstained with DAPI (0.2 ng mL⁻¹; Thermo Fisher Scientific, 62248) during the final wash. Coverslips were mounted using ProLong Gold Antifade Mountant with DAPI (Invitrogen, P36941).

#### Analysis of dendritic arborizations

Immunofluorescence imaging was performed on mature, induced neurons (iNs) following sparse labeling with an hSYN-EGFP expression plasmid introduced by calcium phosphate–mediated transfection. Images were acquired using an EVOS™ M5000 Imaging System equipped with a 20× objective. For each condition, 20 EGFP-positive neurons per biological replicate were analyzed using FIJI. Quantitative morphological analyses included total neurite outgrowth, defined as the summed length of all neurites per neuron; the number of processes, defined as primary neurites emerging from the soma; neurite branch points; and cell body area.

#### Synaptic puncta density analysis

Immunofluorescence analyses were performed on mature, induced neurons stained for microtubule-associated protein 2 (MAP2) to label dendrites and either Synapsin I/II (SYN1) or PSD95 to label pre- or post-synaptic puncta, respectively. Confocal images were acquired using a Leica SP8 inverted confocal microscope equipped with a 40× oil-immersion objective. Z-stack images were collected from ten randomly selected fields per condition, converted to maximum-intensity projections, and analyzed using FIJI software. Cell bodies were masked prior to analysis. Synaptic puncta number was normalized to the total MAP2-positive area to calculate puncta density, or to the mean MAP2 fluorescence intensity to calculate puncta intensity. In addition, puncta size and fluorescence intensity were quantified.

#### Colocalization analysis

Synaptic colocalization was assessed between presynaptic SYN1 and postsynaptic density protein 95 (PSD95). To minimize background and non-synaptic signal, particle size thresholds were applied in FIJI prior to analysis. SYN1-positive puncta were segmented using a size range of 0.05– 3 μm², whereas PSD95-positive puncta were segmented using a size range of 0.05–1 μm². Following thresholding and size filtering, binary masks were generated for each channel, and colocalized puncta were defined as pixels positive for both SYN1 and PSD95. The degree of synaptic colocalization was quantified using Manders’ overlap coefficient, calculated as the fraction of double-positive signal. Identical acquisition settings, thresholding parameters, and size filters were applied across all experimental conditions.

#### Multielectrode array (MEA) recordings

Neuronal network activity was recorded using the Maestro Pro multielectrode array system (Axion Biosystems). Recordings were performed for 10 min per well under controlled conditions (37 °C, 5% CO₂). Signals were acquired at a sampling rate of 12.5 kHz per channel and filtered using a Butterworth band-pass filter (200–3,000 Hz). Both positive- and negative-deflecting spikes were detected using an adaptive threshold set at six times the standard deviation of the baseline noise. Stimulation and pharmacological assays were primarily conducted at day 35 of neuronal differentiation. For inhibitor experiments, baseline activity was recorded prior to compound addition. Inhibitors were prepared as 10× stock solutions and rapidly added to the existing culture medium at 1/9 of the total volume to achieve the desired final concentration (10 µM). Stock solutions were prepared in DMSO, with the final DMSO concentration maintained below 0.1%. Following compound addition, neuronal activity was recorded for an additional 10 min. MEA data analysis was performed using the Neural Metrics Tool software package (Axion Biosystems).

#### Immunohistochemistry (IHC) and immunofluorescence (IF) of postmortem human brain tissue and organoids

Formalin-fixed, paraffin-embedded (FFPE) human brain and organoid tissues were sectioned at 5 µm, mounted onto positively charged glass slides, and baked at 70°C prior to staining. Hematoxylin and eosin (H&E) staining was performed to assess tissue morphology and overall tissue quality. Single- and multiplex immunohistochemistry (IHC) and immunofluorescence (IF) were performed using the Ventana DISCOVERY ULTRA automated staining system (Roche Diagnostics) according to the manufacturer’s instructions at the Neuropathology Brain Bank and Research CoRE (RRID: SCR_027565). For postmortem human fetal brain analyses, cortical tissue from two male cases at 32 and 35 gestational weeks was stained for H3K4me3Q5ser (Millipore, ABE2580) and QR2/NQO2 (Biorbyt, orb400516). Targets were examined by both single-plex chromogenic IHC and duplex chromogenic IHC. For duplex staining, H3K4me3Q5ser and QR2/NQO2 were detected on the same tissue section using the DISCOVERY Purple HRP Kit (Roche, 760-229) and DISCOVERY Green HRP Kit (Roche, 760-271), respectively. For organoid analyses, organoids were fixed in 4% paraformaldehyde, paraffin embedded, sectioned, and analyzed by multiplex IF. Wild-type organoids were compared with the QR2/NQO2 I129E/I195R mutant line using three staining combinations: SOX2 (Active Motif, 39843) together with CTIP2/BCL11B (Abcam, ab18465) at 1 month; NeuN (Abcam, ab104224) together with VGluT1/SLC17A7 (Abcam, ab227805) at 3 months; and H3K4me3Q5ser together with QR2/NQO2 at both 1 and 3 months. Fluorescent detection was performed using species-appropriate secondary detection reagents and the DISCOVERY FAM Kit (Roche, 760-243) and DISCOVERY Cy5 Kit (Roche, 760-238). Fluorescence channels were acquired separately and combined to generate multiplex composite images.

#### RNAscope *in situ* hybridization

The RNAscope assay was performed at the Neuropathology Brain Bank and Research CoRE at Mount Sinai (RRID: SCR_027565) using the Leica BOND RX (LS) automated staining platform. Formalin-fixed, paraffin-embedded (FFPE) organoid sections (4 µm thick). Sections were deparaffinized and pretreated with ER2 buffer pH9 at 88 °C for 15 min. RNAscope in situ hybridization was performed using the RNAscope™ 2.5 LS assay (Cat. No. 332150, Advanced Cell Diagnostics) with the RNAscope™ probe targeting human EMX1 (Hs-EMX1; Cat. No. 801458, ACD Bio) and EMX2 (Hs-EMX2; Cat. No. 533878, ACD Bio). Signal detection was performed using the BOND Polymer Refine Red Detection Kit (DS9390, Leica Biosystems), according to the manufacturer’s instructions. Sections were counterstained with hematoxylin and bluing.

#### Digital whole-slide imaging

Whole-slide digital images were acquired at 40× magnification using the Aperio GT450 high-resolution slide scanner (Leica Biosystems). The digital images are in.scn format and can be visualized with the Aperio ImageScope x64 [v12.4.6.5003] pathology Slide Viewing Software.

#### RNA isolation, library preparation, and sequencing

RNA-seq was performed on HeLa cells, human brain tissues and human iPSC-derived neurons co-cultured with mouse glia across multiple experimental conditions, including genetic perturbations, differentiation stages, and depolarization treatments. All datasets were processed using the same experimental and computational pipelines unless otherwise noted. Cells were lysed in TRIzol reagent (Invitrogen, 15596026), and total RNA was extracted according as follows. Chloroform was added in a 5:1 ratio, and the aqueous phase was isolated. An equal volume of 70% ethanol was added, and the mixture was applied to RNeasy Micro spin columns (Qiagen, 74004). RNA purification was completed following the Qiagen RNeasy Micro Kit protocol, including all optional steps and on-column DNase treatment. RNA was eluted in 15 μL and quantified using a NanoDrop spectrophotometer. Only samples with A260/280 and A260/230 ratios greater than 1.8 were used for downstream applications. For library preparation, 100 ng of total RNA was used as input for the Illumina Stranded mRNA Prep, Ligation kit (Illumina, 20040534). Libraries were amplified with 14 cycles of PCR, and amplified fragments were purified using AMPure XP beads (Beckman Coulter, A63881). Library concentration and fragment size distribution were assessed using a Qubit dsDNA HS assay (Invitrogen, Q33231) and a High Sensitivity D5000 TapeStation assay (Agilent Technologies, 5067-5592, 5067-5593). Libraries were pooled at equimolar concentrations and sequenced on an Illumina NovaSeq X+ platform at the NYU Genome Technology Center. HeLa cell libraries were sequenced to a depth of 50–60 million paired-end reads per sample, whereas libraries generated from human iPSC-derived neurons co-cultured with mouse glia were sequenced to a depth of approximately 100 million paired-end reads (150 bp) per sample to enable robust species-specific read assignment.

#### Differential expression analysis

For HeLa cell RNA-seq and human brain tissue datasets, raw FASTQ files were pseudoaligned and quantified using kallisto (v0.46.1)^77^ against a GRCh38 (hg38) human transcriptome reference derived from Ensembl release 86. For human iPSC-derived neurons co-cultured with mouse glia, reads were pseudoaligned against a combined human–mouse transcriptome reference generated by concatenating Ensembl human (GRCh38, release 86) and mouse (GRCm38/mm10, release 79) transcriptomes, and human-mapped transcripts were retained for downstream analyses. Reads corresponding to the human transcriptome were subsequently extracted for downstream analyses. To reduce noise from lowly expressed transcripts, genes with a total read count of fewer than 15 reads across all samples were excluded. Differential gene expression analysis was performed using DESeq2 (v1.38.3)^78^. Gene-level count matrices were normalized using the DESeq2 median-of-ratios method, and differential expression was assessed using the Wald test. *P* values were adjusted for multiple testing using the Benjamini–Hochberg procedure, with an adjusted *P* value < 0.05 considered statistically significant.

#### Gene overlap analyses

Transcriptome-wide, threshold-free gene expression overlap was visualized using Rank-Rank Hypergeometric Overlap (RRHO) heatmaps generated with the RRHO2 package (v1.0)^79^. Gene lists were ranked by signed p-values, calculated as the log10-transformed nominal p-value multiplied by the sign of the fold change, without applying differential expression thresholds.

#### Cell-type enrichment of the concordant gene set in human developing cortex

Genes in the peak-overlap corner of the rank–rank hypergeometric overlap map (DiffBind fold change > 0 and log2 fold change > 0, i.e. gaining both promoter H3K4me3Q5ser and expression) were tested for cell-type enrichment with Enrichr (enrichR v3.4)^80,81^ against the CellMarker 2024 library^82^. The background was set to the gene universe used in the overlap analysis, comprising all genes carrying a promoter H3K4me3Q5ser peak and detectable expression, rather than the whole genome, since that universe was already restricted to expressed, promoter-marked genes. P values were corrected across terms by the Benjamini–Hochberg procedure, and enrichment magnitude was reported as fold enrichment, the ratio of gene-set to background proportions.

#### Chromatin immunoprecipitation

Untreated and depolarization mouse cultured granule neuron cells (1 × 10⁷ cells per sample) were crosslinked with 1% formaldehyde for 10 min at room temperature and quenched with 125 mM glycine for 5 min. Cells were washed thoroughly prior to lysis and chromatin shearing by sonication, as previously described^14^. Sonicated chromatin was incubated overnight at 4 °C with rotation with antibody-conjugated Dynabeads™ M-280 (7.5 μg antibody per sample). Antibodies used included anti-H3K4me3Q5ser, anti-H3Q5ser, anti-NQO2, and anti–RNA polymerase II. The following day, immunocomplexes were washed, eluted, and reverse-crosslinked, as previously described^14^. After RNase A and proteinase K digestion, DNA fragments were purified using the QIAquick PCR Purification Kit (QIAGEN, 28104).

#### ChIP-seq library preparation and analysis

Purified ChIP DNA was used to prepare sequencing libraries using the NEBNext® Ultra™ II DNA Library Prep Kit for Illumina (NEB, E7645L) following the manufacturer’s protocol. Libraries were sequenced as paired-end reads (50 bp) on an Illumina NovaSeq X+ platform to a depth of approximately 30 million reads per sample. ChIP-seq peak calling and differential analyses were performed as previously described, with minor modifications^14^. Raw sequencing reads were aligned to the mouse reference genome (mm10) using the NGS Data Charmer pipeline with default settings (HISAT v0.1.6b)^83^. Peak calling was performed using MACS2 (v2.1.1)^84^, normalized to the corresponding input controls, with a false discovery rate (FDR) threshold of < 0.01. Differential peak analysis was conducted using pairwise comparisons in the DiffBind package (v3.8.4)^85^ to identify depolarization-induced changes in H3Q5ser, H3K4me3Q5ser, NQO2, and RNA polymerase II (Pol II) occupancy. Identified peaks and differentially enriched regions were annotated to nearby genes or intergenic regions using the ChIPseeker^86^ package in R. Visualization of differential binding profiles was performed using internal functions of DiffBind or deepTools^87^.

### CUT&RUN-seq

#### Cells

Hela cells or human iPSC-derived neurons co-cultured with mouse glia cells or human brain tissues were resuspended in 1 mL of nuclear extract (NE) buffer (20 mM HEPES-KOH, pH 7.9, 10 mM KCl, 0.5 mM spermidine, 0.1% Triton X-100, 20% glycerol and freshly added protease inhibitors (Roche, 11836170001) and passed through a 21-gauge needle 20 times to lyse the cells. Nuclei were pelleted at 1,100*g* for 5 min at 4 °C and the supernatant was discarded. Nuclei were washed again in 1 mL NE buffer and counted. In total, 100,000 nuclei were used per biological replicate.

#### CUT&RUN

CUT&RUN was performed as previously described, with minor modifications^48,88,89^. BioMag Plus Concanavalin A beads (Polysciences, 860573) were prepared by washing 15 μL bead slurry per reaction three times with binding buffer (20 mM HEPES-KOH, pH 7.9, 10 mM KCl, 1 mM CaCl₂, 1 mM MnCl₂) and resuspending the beads in the original volume. Beads (15 μL) were aliquoted into DNA low-bind tubes, and 500 μL nuclear extraction buffer was added. A total of 100,000 nuclei were added to each tube and incubated with beads by end-over-end rotation at room temperature for 10 min. Bead-bound nuclei were washed three times with 1 mL wash buffer (WB; 20 mM HEPES, pH 7.5, 150 mM NaCl, 0.1% Triton X-100, 0.1% Tween-20, 0.5 mM spermidine, 0.1% BSA, freshly supplemented with protease inhibitors). Nuclei were then resuspended in 100 μL antibody buffer (wash buffer supplemented with 2 mM EDTA). Primary antibodies (2 μL per reaction; 1:50 dilution) against H3Q5ser, H3K4me3Q5ser, RNA polymerase II, NQO2, POU2F1, or IgG control were added, and bead-bound nuclei were incubated overnight at 4 °C with gentle horizontal rotation with tubes at a 45 degree angle. The following day, nuclei were washed twice with 1 mL cold wash buffer and resuspended in 50 μL cold wash buffer. Protein A/G–MNase (pAG-MNase; Epicypher, 15-1016; 2.5 μL per reaction) was added, and samples were incubated for 1 h at 4 °C with the same rotation. Nuclei were subsequently washed three times with 1 mL ice-cold wash buffer, followed by one wash with 1 mL low-salt rinse buffer (20 mM HEPES, pH 7.5, 0.5 mM spermidine, 0.1% Tween-20, 0.1% Triton X-100). Bead-bound nuclei were resuspended in ice-cold calcium incubation buffer (3.5 mM HEPES, pH 7.5, 10 mM CaCl₂, 0.1% Tween-20, 0.1% Triton X-100) and incubated on ice (0 °C) for 30 min for MNase digestion. Reactions were quenched by addition of 100 μL of 2× stop buffer (340 mM NaCl, 20 mM EDTA, 5 mM EGTA, 0.1% Tween-20, 0.1% Triton X-100, 25 μg mL⁻¹ RNase A (Thermo Scientific, EN0531), and 0.05 ng per 100 μL *E. coli* spike-in DNA (Epicypher, 18-1401)). Samples were incubated at 37 °C for 30 min without shaking to allow chromatin release and RNA digestion. Beads were captured using a magnetic rack, and the supernatant (200 μL) containing released DNA fragments was collected. DNA was purified using the Zymo ChIP DNA Clean & Concentrator kit (Zymo Research, D5205), eluted in 30 μL, and stored at −20 °C for library preparation.

#### CUT&RUN-seq library preparation and sequencing

Library preparation was performed using the NEBnext Ultra II DNA library kit (E7645L) with multiplexed adapters with minor modifications. CUT&RUN DNA underwent end repair and adapter ligation according to the manufacturer’s protocol (1:15 adapter dilution). DNA was amplified using 16 PCR cycles with an amended 10 s of extension time per cycle to enrich for short fragments. DNA was enriched for 150–350 bp fragments for transcription factor binding events and 150–800 bp for histone markers using Ampure XP beads cleanup. Libraries were quantified with Qubit dsDNA HS assay (Invitrogen, Q33231), and the size distribution was determined by High Sensitivity D5000 TapeStation assay (Agilent Technologies, 5067-5592, 5067-5593). Libraries were pooled at an equimolar concentration and sequenced on the Illumina NovaSeq X+ sequencer by the NYU Genome Technology Center. HeLa cell libraries were sequenced to a depth of 20–30 million paired-end reads per sample, whereas libraries generated from human iPSC-derived neurons co-cultured with mouse glia were sequenced to a depth of approximately 50 million paired-end reads (150 bp) per sample to enable robust species-specific read assignment.

#### CUT&RUN-seq data analysis

CUT&RUN data were analyzed as previously described, with minor modifications^89^. For human iPSC-derived neurons co-cultured with mouse glia, reads were aligned using Bowtie2 (v2.5.0)^90^ to a concatenated reference genome comprising the human genome (GRCh38/hg38), the mouse genome (GRCm38/mm10), and the Escherichia coli genome used as a spike-in control. Paired-end reads were aligned using Bowtie2 (v2.5.0) with the --very-sensitive preset and Phred+33 encoding. Insert size parameters were set to 10–2000 bp (-I 10 -X 700), and dovetailing read pairs were permitted (--dovetail). For HeLa cell samples, reads were aligned to a concatenated reference consisting of GRCh38/hg38 and the *E. coli* genome. Aligned reads were filtered using SAMtools (v1.9)^91^ to retain only uniquely mapped reads with a minimum mapping quality (MAPQ) score of 30. Reads were subsequently separated by species (human, mouse, and *E. coli*) for downstream analyses. For normalization and quantitative comparisons across samples, *E. coli* spike-in DNA was used to calculate a scaling factor for each sample, defined as the median number of uniquely mapped *E. coli* reads across all samples divided by the number of *E. coli* reads in the given sample. This scaling factor was applied to normalize human-aligned read counts and genome coverage tracks. Genome coverage tracks were generated as bigWig files using the deepTools (v3.5.5)^87^ bamCoverage function, and regions with consistently high background signal were excluded using the ENCODE hg38 blacklist (v2). Heatmaps were generated using deepTools (v3.5.5) computeMatrix and plotHeatmap functions in reference-point mode, centered on peak centers. Peak calling was performed using MACS2^84^ with a false discovery rate (FDR) threshold of < 0.01. Differential peak analysis was conducted using DiffBind^85^ by quantifying read counts across consensus peak regions and statistically testing for condition-dependent differences in chromatin occupancy. Identified peaks and differentially enriched regions were annotated, and motif enrichment analyses were performed using the ChIPseeker^86^ package in R.

#### Odds ratio analysis integrating RNA-seq and CUT&RUN

To quantify the association between transcriptional changes and chromatin occupancy, an odds ratio (OR) framework was used to integrate RNA-seq and CUT&RUN datasets. Differential gene expression was assessed for each mutant relative to wild-type (WT) using DESeq2^78^, and differentially expressed genes (DEGs) were defined as genes with an adjusted P value < 0.05 and an absolute log₂ fold-change ≥ 0.5. CUT&RUN peaks were called per biological replicate using MACS2^84^, and differential binding between WT and mutant conditions was assessed using DiffBind^85^. For each CUT&RUN dataset (NQO2, H3K4me3Q5ser, or RNA polymerase II), WT-enriched CUT&RUN peaks were annotated to genes using ChIPseeker^86^ by assigning peaks to the nearest transcription start site (TSS), with promoter regions defined as ±3 kb around the TSS (tssRegion = −3,000 to +3,000). Annotations were performed using TxDb.Hsapiens.UCSC.hg38.knownGene together with the org.Hs.eg.db database. Genes associated with at least one WT-enriched CUT&RUN peak within this window were classified as “bound.” For each mutant and each CUT&RUN dataset, enrichment was tested using a 2 × 2 contingency table constructed across the expressed gene universe (genes passing RNA-seq filtering), stratified by (i) DEG status (DEG vs non-DEG) and (ii) CUT&RUN association (bound vs unbound). Odds ratios, 95% confidence intervals, and two-sided P values were computed using Fisher’s exact test^92^. Odds ratios greater than 1 indicate enrichment of DEGs among genes associated with WT-enriched CUT&RUN signal. Where applicable, P values were adjusted for multiple testing as indicated.

### ATAC-seq

#### Nuclei preparation

Nuclei from induced neurons (iNs) were prepared using the same nuclei isolation procedure as described for CUT&RUN. For each biological replicate, 100,000 nuclei were used as input for ATAC-seq library preparation.

#### ATAC-seq library preparation

ATAC-seq libraries were generated using the ATAC-Seq Express Kit (Active Motif, 53157) according to the manufacturer’s instructions. Libraries were size-selected using SPRIselect Bead-Based Reagent (Beckman, B23318) to enrich for fragments ranging from 200 to 800 bp. Library quality and fragment size distribution were assessed using a High Sensitivity D5000 TapeStation assay (Agilent Technologies; 5067-5592, 5067-5593). Libraries were pooled at equimolar concentrations and sequenced as 150-bp paired-end reads on an Illumina NovaSeq X+ platform at the NYU Genome Technology Center, yielding approximately 60 million reads per sample.

#### ATAC-seq data processing and analysis

Primary ATAC-seq data processing included adapter trimming, alignment to a concatenated human–mouse reference genome comprising the human genome (GRCh38/hg38) and the mouse genome (GRCm38/mm10) using Bowtie2^90^, and removal of mitochondrial reads and PCR duplicates. Peak calling was performed independently for each of the three biological replicates using MACS2^84^, followed by assessment of reproducibility across replicates. Normalization and differential chromatin accessibility analyses were conducted using DiffBind^85^. Transcription factor footprinting analysis was performed using TOBIAS^93^, which corrects ATAC-seq signal for Tn5 transposase sequence bias and local chromatin accessibility prior to footprint inference. Differential transcription factor binding activity was quantified using the BINDetect module by comparing footprint depth and flanking accessibility at motif instances, enabling condition-specific comparison of POU2F1 binding scores between wild-type and mutant samples. To assess whether loss of chromatin accessibility in mutant neurons was associated with loss of POU2F1 (OCT1) binding motifs, we performed motif scanning of ATAC-seq peaks using POU2F1 consensus motifs from the JASPAR database (MA0785.1 and MA0785.2). Genomic coordinates of ATAC-seq peaks were scanned for motif occurrences to classify loci as OCT1 motif–positive (OCT1⁺) or OCT1 motif–negative (OCT1⁻). Differential accessibility analysis was used to identify ATAC peaks showing reduced accessibility in mutant compared with wild-type hiPSC-derived neurons.

#### Dorsally patterned forebrain organoid differentiation

Cortical organoids were generated from two pluripotent stem cell lines — a wild-type control (WT) and QR2 I129EI195R mutant line — using the dorsal forebrain differentiation strategy of paper^55^ with minor adaptations. At the onset of differentiation (day 0), pluripotent colonies were enzymatically dissociated into a single-cell suspension with Accutase (Gibco), and 9,000 cells were seeded per well into V-bottomed, ultra-low-attachment 96-well plates (PrimeSurface; Sumitomo Bakelite) to promote formation of uniform embryoid bodies (EBs). Aggregation was performed in Cortical Differentiation Medium I (CDM I): Glasgow-MEM (Gibco) supplemented with 20% (v/v) Knockout Serum Replacement (Gibco), 0.1 mM non-essential amino acids (Gibco), 1 mM sodium pyruvate (Gibco), 0.1 mM 2-mercaptoethanol (Gibco), and 100 U/mL penicillin–100 µg/mL streptomycin (Corning). To support single-cell survival during reaggregation, the ROCK inhibitor Chroman1 (Tocris Bioscience; 20 µM) was supplied from day 0 to day 6. Dorsal forebrain fate was imposed by simultaneous pathway inhibition between day 0 and day 18, combining the WNT inhibitor IWR1 (Calbiochem; 3 µM) with the TGFβ inhibitor SB431542 (STEMCELL Technologies; 5 µM). On day 18, EBs were transferred to ultra-low-attachment dishes (Corning) and placed on an orbital shaker for the remainder of the culture, with the medium exchanged for CDM II: DMEM/F12 (Gibco) containing 2 mM GlutaMAX (Gibco), 1% (v/v) N-2 supplement (Gibco), 1% (v/v) Chemically Defined Lipid Concentrate (Gibco), 0.25 µg/mL Fungizone (Gibco), and penicillin–streptomycin as above. From day 35, aggregates were moved into spinner-flask bioreactors (Corning) and cultured in CDM III, generated by supplementing CDM II with 10% (v/v) fetal bovine serum (GE Healthcare), 5 µg/mL heparin (Sigma), and 1% (v/v) Matrigel (Corning). Beginning on day 70, organoids were maintained in CDM IV, comprising CDM III additionally supplemented with B-27 (Gibco) and 2% (v/v) Matrigel. For both genotypes, organoids were collected at two differentiation stages — approximately one month (∼day 30) and three months (∼day 90) — to sample an early and a later stage of cortical development.

#### Dissociation of brain organoids, single nucleus multiome library preparation, and sequencing

Single-nucleus multiome profiling was performed across four experimental groups: two genotypes, wild-type (WT) and the mutant line QR2 (I129EI195R), each collected at two differentiation stages, approximately one and three months. For each condition, approximately ten organoids were pooled as a single sample, giving four samples in total. Nuclei were isolated by transferring pooled organoids into freshly prepared, ice-cold nuclei isolation buffer (10 mM Tris-HCl [pH 7.4], 10 mM NaCl, 3 mM MgCl₂, 0.01% [v/v] Triton X-100, 1% [w/v] BSA, 1 mM DTT, 1 U/µL SUPERase-In RNase Inhibitor, 1 U/µL Takara Recombinant RNase Inhibitor, and 1× protease inhibitor in nuclease-free water). Organoids were incubated on ice in this buffer and homogenized with a Dounce homogenizer to release nuclei, and the homogenate was filtered through a 40 µm pluriStrainer Mini strainer to yield a single-nucleus suspension. Nuclei were pelleted by centrifugation, resuspended in chilled Nuclei Buffer, counted, and adjusted to a concentration targeting ∼10,000 nuclei per reaction.

Paired single-nucleus gene expression and chromatin accessibility libraries were generated from each sample with the Chromium Next GEM Single Cell Multiome ATAC + Gene Expression kit (10x Genomics, PN-1000283), following the manufacturer’s protocol (CG000338 Rev G). In brief, nuclei were transposed in bulk to tagment accessible chromatin, then loaded onto a Chromium Next GEM Chip J and partitioned into gel bead-in-emulsion (GEM) droplets on the Chromium instrument, within which transposed DNA fragments and mRNA-derived cDNA from each nucleus acquired a common 16-bp cell barcode. After GEM incubation and cleanup, the barcoded material was pre-amplified by PCR and split to construct the two library types in parallel. ATAC libraries were completed by sample-index PCR with the Single Index Kit N Set A (PN-1000212) and a double-sided size selection; gene expression libraries were built from the amplified cDNA by fragmentation, end repair and A-tailing, adaptor ligation, and sample-index PCR with the Dual Index Kit TT Set A (PN-1000215). PCR cycle numbers followed the input-dependent recommendations of the kit. Final libraries were assessed for fragment-size distribution on the Agilent High Sensitivity D5000 ScreenTape assay (Agilent Technologies, Cat# 5067-5592 and 5067-5593) and quantified with the Qubit 1× dsDNA HS Assay Kit (Invitrogen, Cat# Q33231) before pooling.

Libraries were sequenced on an Illumina NovaSeq X Plus with a 25B 300-cycle flow cell. Gene expression and ATAC libraries were co-sequenced using a 50 (Read 1) × 10 (i7) × 24 (i5) × 100 (Read 2) cycle configuration, which supports both the ATAC read structure (Read 1, 50 bp; i5, 24 cycles encoding the cell barcode) and the gene expression read structure (Read 2, 100 bp). Target sequencing depths were approximately 500 million read pairs per gene expression library and 1 billion read pairs per ATAC library.

#### Single-nucleus multiome processing and quality control

Raw sequencing reads were processed with Cell Ranger ARC (10x Genomics, v2.0.0) against the GRCh38 (hg38) reference to produce per-nucleus gene-expression matrices, ATAC peak matrices and fragment files. All downstream analysis was performed in R (v4.4.3) using Seurat (v5.5.0)^94^ and Signac (v1.16.0)^95^, in a conda-managed environment (conda v26.1.1, Python v3.10.20), with a fixed random seed (1234) throughout. For each sample, gene-expression counts were loaded into a Seurat object and ATAC peak counts, restricted to standard chromosomes, into a Signac ChromatinAssay annotated with hg38 gene models (EnsDb.Hsapiens.v86 v2.99.0, ensembldb v2.30.0^96^, biovizBase v1.54.0, BSgenome.Hsapiens.UCSC.hg38 v1.4.5). Per-nucleus quality metrics were computed — RNA and ATAC fragment counts, transcription start site (TSS) enrichment, nucleosome signal and mitochondrial content — and nuclei were retained at > 500 RNA counts, > 500 ATAC counts, TSS enrichment > 2, nucleosome signal < 2 and < 5% mitochondrial reads. Ambient RNA contamination was estimated per nucleus with decontX (celda v1.22.0)^97^ and doublets were called from the RNA counts with scDblFinder (v1.23.4)^98^; nuclei with an estimated contamination fraction ≥ 0.40, or classified as doublets, were removed. Contamination estimates served only as a quality-control filter, and raw counts were carried into all downstream analysis.

#### Peak calling and construction of a union peak set

For each sample, ATAC peaks were called on the filtered nuclei with MACS3 (v3.0.4) via Signac CallPeaks, retained on standard chromosomes, filtered against the ENCODE hg38 blacklist (v2) and restricted to widths of 20–10,000 bp. Sample-level peaks were combined into a genome-wide union peak set (GenomicRanges v1.58.0, GenomeInfoDb v1.42.0, rtracklayer v1.66.0), re-filtered by width, and each sample was re-quantified against this common peak set with Signac FeatureMatrix before the re-quantified objects were merged into a single dataset.

#### Normalization, integration and clustering

Gene expression was normalized with SCTransform^99^ and reduced by principal-component analysis (30 components). Chromatin accessibility was normalized by term frequency–inverse document frequency, restricted to the top features (top quartile) and decomposed by singular value decomposition, retaining latent semantic indexing components 2–30. The two modalities were integrated by weighted nearest-neighbour (WNN) analysis and visualized by WNN-based UMAP. Nuclei were clustered on the WNN graph with the SLM algorithm across resolutions 0.1–1.0 (clustree v0.5.1, patchwork v1.3.2, future v1.70.0), and a resolution of 0.8 was selected. Cluster marker genes were identified on the SCT assay with FindAllMarkers after PrepSCTFindMarkers (Wilcoxon rank-sum test, positive markers only, log fold-change threshold 0.25), and cell identities were assigned from canonical cortical cell-type markers. Data handling and visualization used dplyr (v1.2.1), readr (v2.2.0) and ggplot2 (v4.0.3).

#### Pseudobulk chromatin-accessibility tracks and peak calling

Single-nucleus ATAC fragments were pooled by genotype × age × broad cell type — wildtype and QR2 I129EI195R at one and three months — and combinations with ≥ 50 nuclei were retained. For each pseudobulk, fragments were restricted to chromosomes in the hg38 chromosome-size file, sorted and indexed (bgzip/tabix). CPM-normalized coverage was generated with BEDTools genomecov (−bg; scale = 10⁶ / total fragments) and converted to bigWig with bedGraphToBigWig. Accessible peaks were called per pseudobulk with MACS3 (v3.0.4) (callpeak −f BED −g hs −−nomodel −−shift −100 −−extsize 200 −−keep-dup all −q 0.05).

#### Cell-type accessible peaks, genomic annotation and peak-set intersections

Within each broad cell type, a peak was defined as accessible if detected at ≥ 1 count in ≥ 5% of that cell type’s nuclei. Peaks were assigned a genomic context from the Signac gene annotation under the priority promoter (± 2 kb of a TSS) > exon > intron > distal. Cross-cell-type sharing was computed from a binary peak × cell-type membership matrix over the six broad cell types and visualized as an Upset plot.

#### Marker-locus accessibility tracks

Aggregated per-cell-type ATAC coverage at canonical marker loci was plotted with Signac CoveragePlot, grouped by broad cell type, with peaks shown and ± 50 kb flanks.

#### Peak-to-gene linkage

Peak-to-gene links were computed on the WNN-integrated object with Signac LinkPeaks, correlating peak accessibility (ATAC assay) with SCT-normalized expression across all nuclei (Pearson correlation; distance 2 Mb; min.cells = 10; n_sample = 100), after per-peak sequence statistics were added with RegionStats against BSgenome.Hsapiens.UCSC.hg38. Links were retained at P < 0.05 and correlation score ≥ 0.1. Because LinkPeaks operates across all cells, a single link set was used for every cell-type analysis. For the paired heatmap, the most significantly correlated peak per gene was retained, and its accessibility together with the linked gene’s expression was row-scaled (z-score, clipped at ± 2) and grouped by cell type.

#### Genome-browser coverage with peak-to-gene arcs

Per-cell-type ATAC coverage was plotted with CoveragePlot over each locus, and retained peak-to-gene links (|score| ≥ 0.1) were drawn as arcs from each linked peak centre to the target-gene TSS, with gene models from EnsDb.Hsapiens.v86 (GRCh38). Strand-aware windows were applied per gene: EMX1, 50 kb upstream and 100 kb downstream; EMX2, 50 kb upstream and 30 kb downstream; all other loci, ± 500 kb.

#### Cell-type composition and excitatory-to-progenitor ratio

Composition was summarised from the broad annotation. For each genotype and age, dorsal progenitors were counted as the union of the radial glia and cortical hem classes and excitatory neurons as the cortical excitatory neuron class; outer radial glia/astroglial progenitors, Cajal– Retzius/preplate neurons, interneurons, mesenchymal-like and mixed classes were tabulated separately and contribute to neither term. The excitatory-to-progenitor ratio is the quotient of these two counts, and the fold difference between genotypes at one month is the ratio of those quotients.

#### Cell-cycle phase

Phase was assigned with Seurat CellCycleScoring using the S and G2/M gene sets in cc.genes.updated.2019, scored on the SCT assay with set.ident = FALSE. Phase fractions were then computed within the radial glia class at one month for each genotype; the cycling fraction is the sum of the S and G2/M fractions. Phase inferred from single-nucleus RNA is a coarse estimate of proliferative state and was not corroborated by a labelling method.

#### Progenitor differentiation potency

Developmental potency was scored with CytoTRACE 2 (v1.1.0)^59,100^ a deep-learning framework in which Gene Set Binary Network modules learn the gene sets associated with each potency category; the released ensemble comprises 19 models trained across 19 datasets from 16 human and mouse tissues, and prediction is made over a fixed panel of 14,271 gene features rather than from hand-curated stemness markers. Raw RNA counts for dorsal progenitors — the radial glia and cortical hem classes — at one month were supplied as a dense expression matrix with species = “human” and is_seurat = FALSE; batch_size, smooth_batch_size and seed were left at their defaults of 10,000, 1,000 and 14. Wild-type and QR2 I129EI195R nuclei were scored together in a single run, so the postprocessing steps — Markov diffusion smoothing, binning within predicted potency categories, and adaptive nearest-neighbour smoothing — drew on the pooled dataset. We report the postprocessed absolute potency score (CytoTRACE2_Score), a continuous value from 0 (differentiated) to 1 (totipotent). Genotypes were compared with a two-sided Wilcoxon rank-sum test. Because the comparison is made across nuclei within a single clonal line per genotype rather than across lines, we treat the effect size rather than the p value as the measure of magnitude, reporting Cliff’s δ computed from the rank-sum statistic as 2U/(n₁n₂) − 1, with |δ| of 0.147, 0.33 and 0.474 taken as the boundaries between negligible, small, medium and large effects, together with the difference in medians and its 95% percentile confidence interval from 2,000 bootstrap resamples drawn within each genotype.

#### Pseudobulk gene set enrichment analysis

Raw RNA counts for the radial glia class at one month, the cortical hem excluded, were summed across nuclei within each genotype to give one pseudobulk profile per genotype. Sex-chromosome genes (XIST, TSIX, RPS4Y1, RPS4Y2, DDX3Y, UTY, KDM5D, EIF1AY, USP9Y, NLGN4Y, TXLNGY, ZFY, PRKY, TMSB4Y) and mitochondrially encoded (MT-) transcripts were removed, and genes with fewer than 10 summed counts across the two profiles were discarded. The remaining counts were TMM-normalised in edgeR and converted to log₂ counts per million with a prior count of 3, and genes were ranked by the difference between the mutant and wild-type profiles. Enrichment was tested against the MSigDB Hallmark collection^101^ with fgsea^102^ using gene sets of 10 to 500 genes, and p values were adjusted across the 50 Hallmark sets by the Benjamini–Hochberg procedure. Because each genotype contributes a single summed profile, the contrast carries no replicate-level dispersion estimate, and the fgsea p values are computed by gene permutation: they describe how coherently a set is ranked within this one contrast rather than significance replicated across lines or samples. The full result is given in Data S22.

#### POU2F1 motif-linked peak-to-gene concordance

Motif matches were added to the ATAC assay with Signac AddMotifs using the JASPAR2022 CORE (Homo sapiens) collection and BSgenome.Hsapiens.UCSC.hg38, and peaks containing a POU2F1 motif were identified. Positive POU2F1-linked peak-to-gene links (score ≥ 0.10) were retained. For each genotype × age × cell type with ≥ 50 nuclei, mean accessibility of POU2F1-linked peaks (ATAC counts) and mean expression of their linked genes (SCT data) were computed from a cell-membership design matrix; log₂ fold changes between conditions were calculated with a 1 × 10⁻⁴ pseudocount, and accessibility and expression changes were scored as concordant when they shared sign.

## ACKNOWLEDGEMENTS

We would like to thank members of the Maze lab for their helpful discussions on this study. This work was partially supported by grants from the National Institutes of Health: R01 MH116900 (I.M.), F32 MH140478 (B.H.W.), F32 MH126534 (J.C.O’C.), R01 NS136448 (M.H.), R21 NS130319 (S.G.M.), as well as funds from the Brain and Behavior Research Foundation (J.C.C.), the Damon Runyon Cancer Research Foundation (DRG-2506-23; J.Z.), the Cindy Silvian Foundation (S.G.M.), and the Howard Hughes Medical Institute (I.M.). Additional support was provided by the National Natural Science Foundation of China (22425702 to X.D.L.; 22577080 to X. L.), the General Research Fund (GRF 17107123, 17310122, 17302524, 17102124, 17104923, and 17311025 to X.D.L.), the Large Research Equipment Fund 2022–2023 of the University of Hong Kong (X.D.L.), and the Guangdong Provincial Project (2023QN10C205 to X.D.L.). All schematics were created with Biorender.com. Experiments were supported by the Stem Cell Engineering Core (RRID:SCR_027503) at the Icahn School of Medicine at Mount Sinai. We thank R. M. Vabulas for providing plasmids for the expression of QR1 and QR2 in *E. coli*.

## AUTHOR CONTRIBUTIONS

M.C., C.Y., X.L., X.D.L. H.L., and I.M. conceived of the study, designed the experiments, and interpreted the data. M.C. performed most biochemical-, molecular-, epigenomic-, and cellular-based assays. X.L. performed chemical proteomics experiments. L.K. performed peptide pulldown assays from HeLa NEs and ChIP/Re-ChIP experiments in cGNs. C.Y., X.L., R.Z., and H.L. performed structural and ITC analyses, and interpreted the data. B.H.W., X.W., J.C.O’C, D.A.V., A.R., and L.S. assisted with bioinformatics analyses. B.C. provided general experimental assistance. X.L., Q.Z., and Y.D., generated H3 peptides (modified and unmodified). J.Z, K.M.C., J.R.S, and T.W.M. generated semisynthetic mononucleosomes (modified *vs.* unmodified). K.R. provided QR2 inhibitors. Z.L. and H.M. performed LC-MS/MS analyses. E.B., R.H., and S.G.M. assisted with hiPSC maintenance and gene editing. C.D.S. and V.F.A. assisted with fetal cortex and organoid IHC. M.C., C.Y., X.L., X.D.L., H.L., and I.M. wrote the manuscript.

## DATA AVAILABILITY

The genomics data generated in this study have been deposited in the National Center for Biotechnology Information Gene Expression Omnibus (GEO) database under the SuperSeries GSE324882. All mass spectrometry-based proteomics data have been deposited to the ProteomeXchange Consortium via the PRIDE partner repository with the datasets identified as PXD075254 and PXD075547. The atomic coordinates and structure factors have been deposited in the Protein Data Bank (PDB) under PDB ID codes 24PZ and 24RO. We declare that the data supporting findings for this study are available within the article and Supplementary Information. Related data including raw microscopy images are available from the corresponding author upon reasonable request. No restrictions on data availability apply.

## CODE AVAILABILITY

Related code is available from the corresponding author on reasonable request.

## ETHICS STATEMENT

All collaborators associated with this work meet the authorship criteria outlined by Cell Press journals. Each author made contributions essential to the design, execution, analysis, and/or interpretation of the study. Roles and responsibilities were defined and agreed upon among collaborators at the outset of, or during, the research.

## Supplementary Information

**Figure S1.**
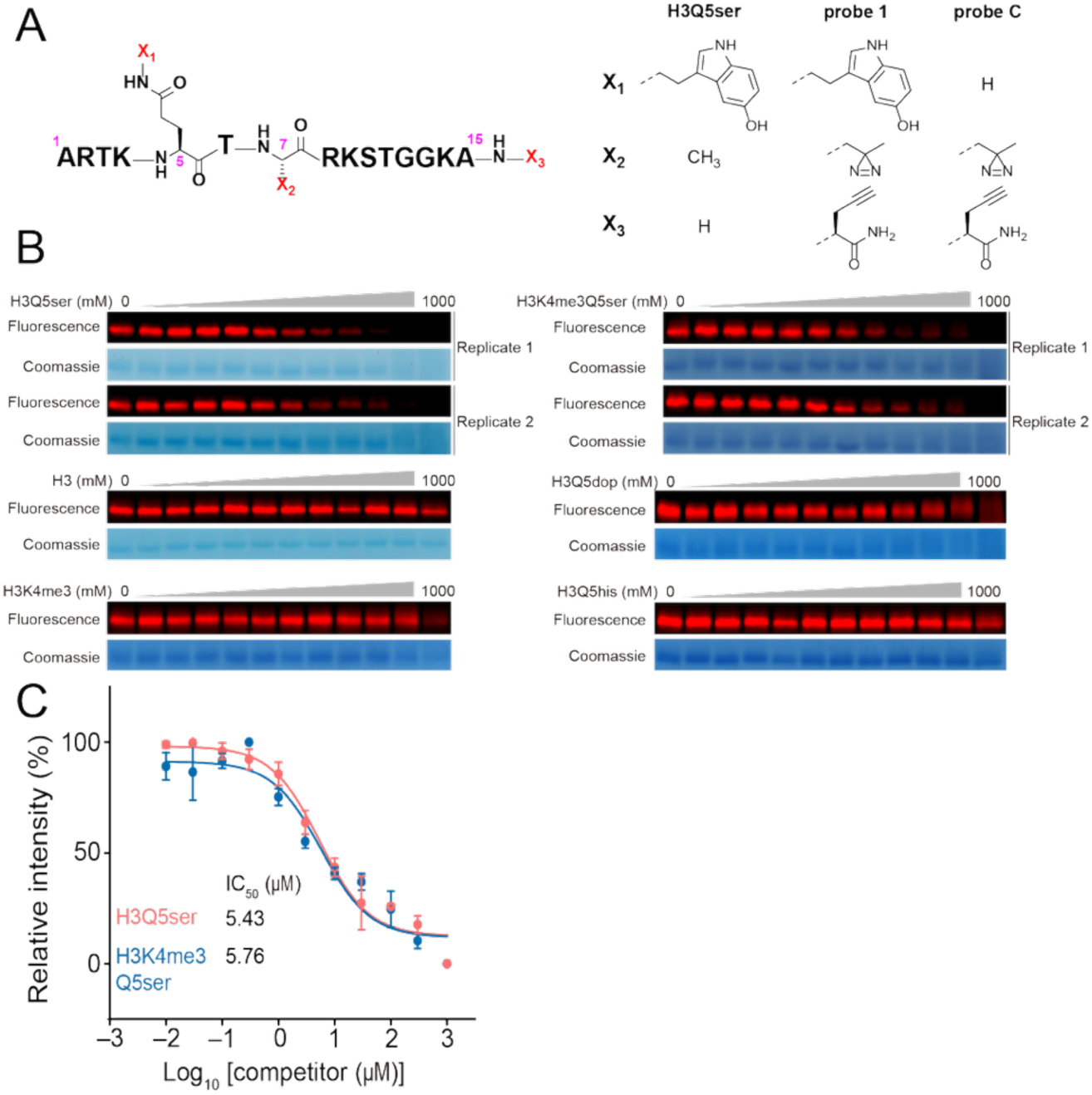
QR2 selectively binds to H3Q5ser, related to Figure 1. (**A**) Chemical structures of H3Q5ser, probe 1, and probe C. (**B**) In-gel fluorescence scanning results of different H3 peptides competing with probe 1-induced labeling of recombinant QR2. (**C**) Determination of IC_50_ for inhibition of probe 1-induced labeling of QR2 by H3Q5ser and H3K4me3Q5ser peptides. The fluorescence intensity of each band was quantified by ImageJ. All curves were normalized between 100 and 0% at the highest and lowest fluorescence intensities, respectively. Data are reported as mean ± SD (*n* = 2). All immunoblotting experiments repeated 3X. See **Figure S14** for uncropped blots.

**Figure S2.**
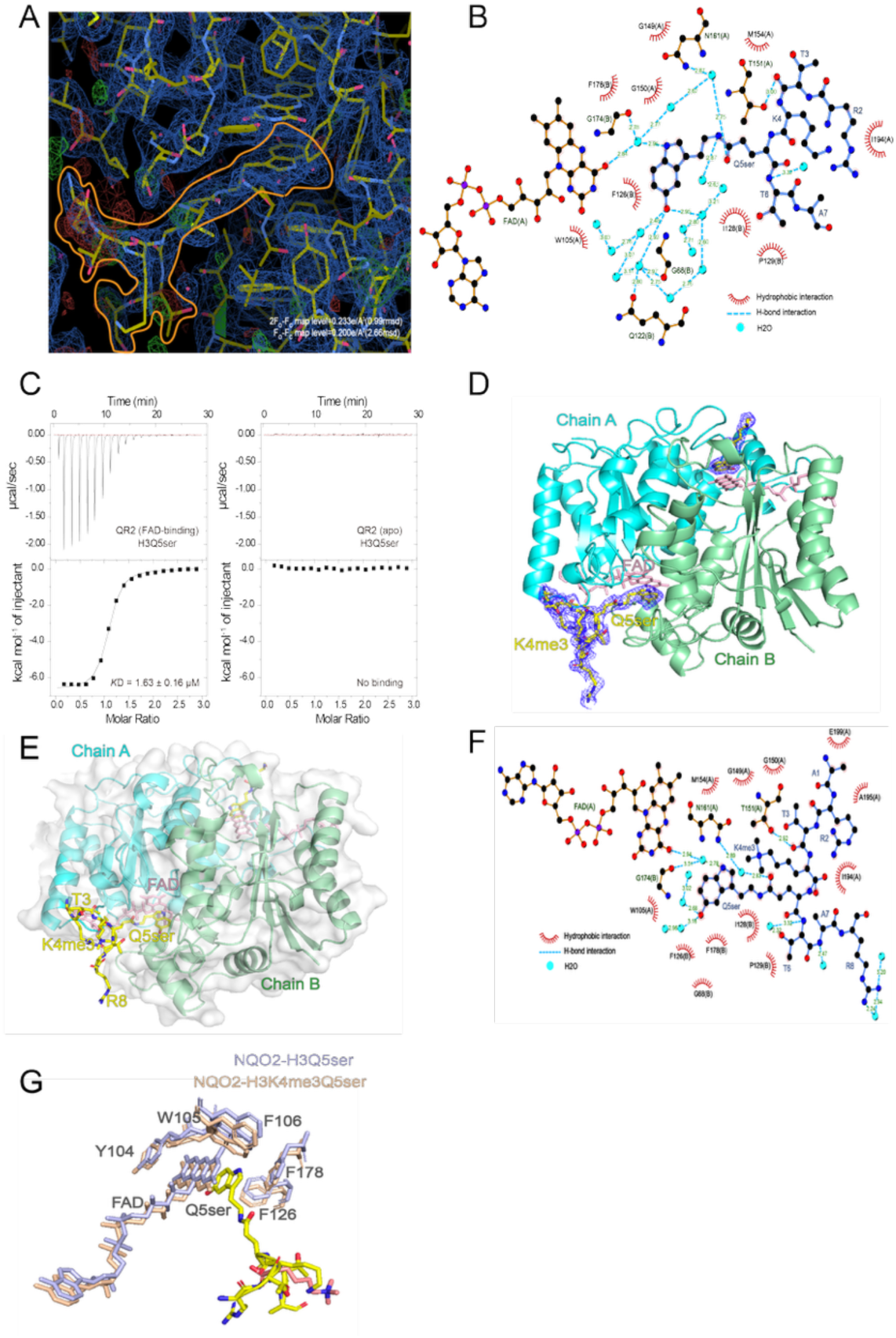
Common molecular recognition of H3Q5ser and H3K4me3Q5ser by QR2, related to Figure 2. (**A**) Corrected difference electron density map of the QR2-H3Q5ser crystal structure. The continuous density marked by the orange line corresponds to the H3Q5ser peptide, while the remaining regions represent the density of QR2. The electron density from the H3R2 to the H3A7 is visible. (**B**) LIGPLOT diagram listing critical contacts between the H3Q5ser peptide and QR2. The main chain of the peptide is shown in blue, surrounded by multiple hydrophobic interactions (red arcs) and several pairs of hydrogen bond interactions (blue lines). The distances of the hydrogen bond interactions are labeled in green text. Water molecules are indicated by cyan balls, and the FAD molecule is also displayed in the diagram. The letters A and B following the amino acid residues denote Chain A and Chain B in the QR2 dimer, respectively. Numbering of residues excludes the initiating methionine of QR2. (**C**) Titration and fitting curves of peptides titrated into wildtype and *apo* (depleted of FAD) QR2. *K*D values are provided. See **Table S1** for ITC statistics. (**D**) Structure of the QR2-H3K4me3Q5ser complex. The blue mesh shows the electron density of the H3K4me3Q5ser peptide. The Q5ser residue inserts into the binding pocket of QR2, while the K4me3 residue extends out from the QR2 surface and does not participate in recognition. (**E**) The QR2-H3K4me3Q5ser co-crystal structure reveals that Q5ser inserts into the interior of the molecule, while T3 and K4me3 are oriented toward the exterior and do not participate in recognition. (**F**) LIGPLOT diagram listing critical contacts between the H3K4me3Q5ser peptide and QR2. The main chain of the peptide is shown in blue, and K4me3 is visible in the diagram. Multiple hydrophobic interactions (red arcs) surround Q5ser, which are highly consistent with those observed in the NQO2-H3Q5ser complex. Numbering of residues excludes the initiating methionine of QR2. (**G**) Comparison of the conformations of key amino acids involved in recognition between the QR2-H3K4me3Q5ser co-crystal structure (pink) and the QR2-H3Q5ser co-crystal structure (purple). Their conformations are completely identical. The unmodified K4 is shown in yellow, while the K4me3 modification is shown in red. See **Table S2** for x-ray crystallography data collection and refinement statistics.

**Figure S3.**
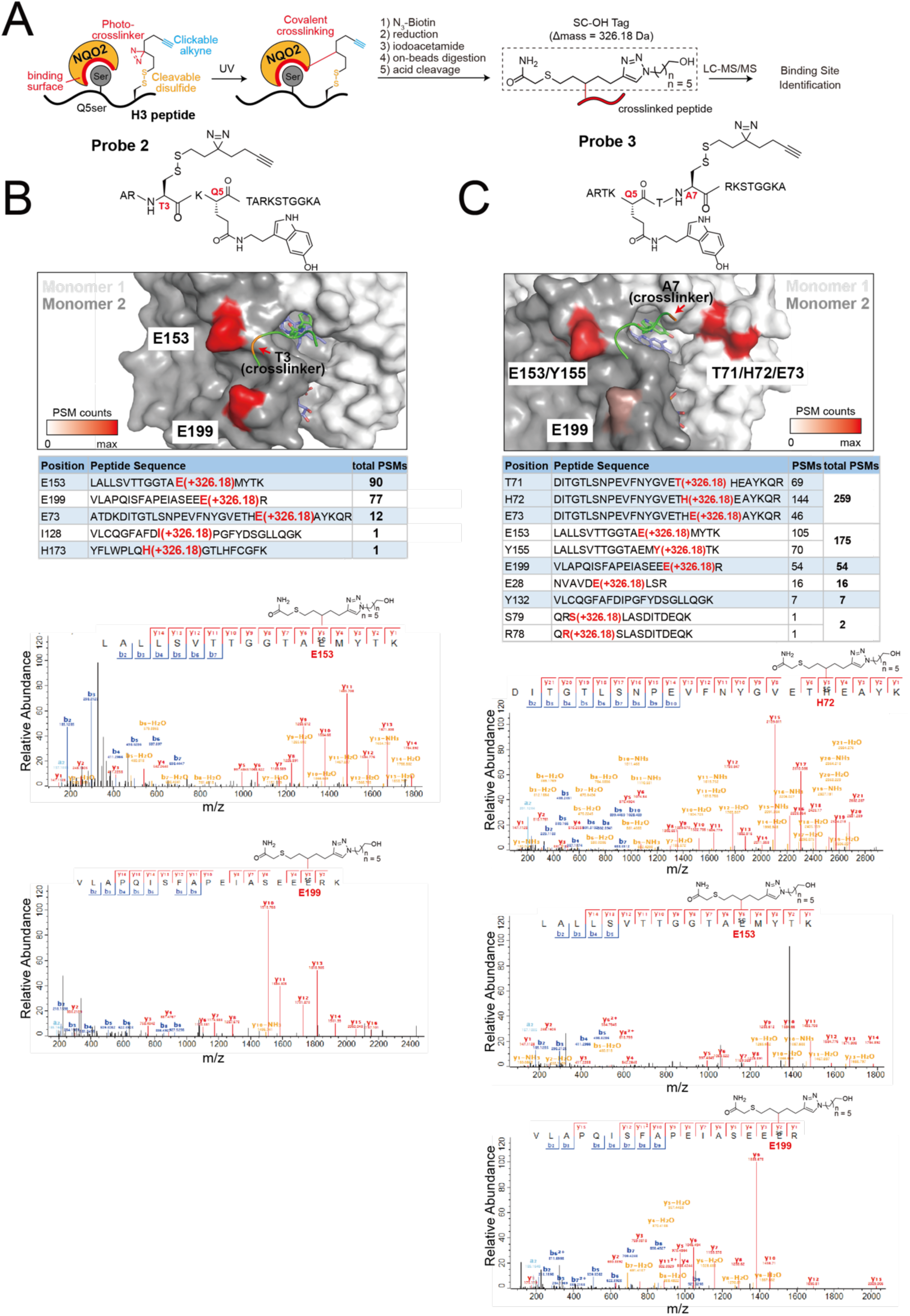
Photoaffinity probe-based mapping of the binding sites between QR2 and the H3Q5ser peptide, related to Figure 2. (**A**) Schematic workflow for binding sites mapping between recombinant QR2 and H3Q5ser peptide. QR2 was incubated with photoaffinity probes carrying serotonylation at Q5 and a tri-functional amino acid ADdis-Cys that contains a diazirine, an alkyne, and a reductively cleavable disulfide bond. Following photo-crosslinking, proteins captured by the probe will be conjugated with biotin-azide carrying an acid-labile dialkoxydiphenylsilane (DADPS) linker and then immobilized by streptavidincoated beads. After reductive cleavage, iodoacetamide-mediated thiol capping, on-bead digestion, and washing steps, the peptides remaining on the beads should be those directly cross-linked to ADdis-Cys with binding sites information. A final acid treatment released the peptides with SC-OH tags (+326.18 Da) installed at the cross-linked residues that will be identifiable by LC–MS/MS analysis. (**B**) Crosslinking sites identified with probe 2 (ADdis-Cys incorporated at T3). Upper: chemical structure of probe 2. Middle: QR2 surface representation highlighting major crosslinked residues (E153, E199) color-coded by peptide–spectrum match (PSM) counts. Lower: summary table listing crosslinked positions, corresponding peptide sequences, and total PSMs; representative MS/MS spectra for E153 and E199 with annotated b- and y-type ions and marked modification sites. (**C**) Crosslinking sites identified with probe 3 (ADdis-Cys incorporated at A7). Upper: chemical structure of Probe 3. Middle: QR2 surface representation highlighting major crosslinked residues (T71, H72, E73, E153, Y155, E199) color-coded by peptide–spectrum match (PSM) counts. Lower: summary table listing crosslinked positions, corresponding peptide sequences, and total PSMs; representative MS/MS spectra for T71/H72/E73, E153, and E199 with annotated b- and y-type ions and marked modification sites.

**Figure S4.**
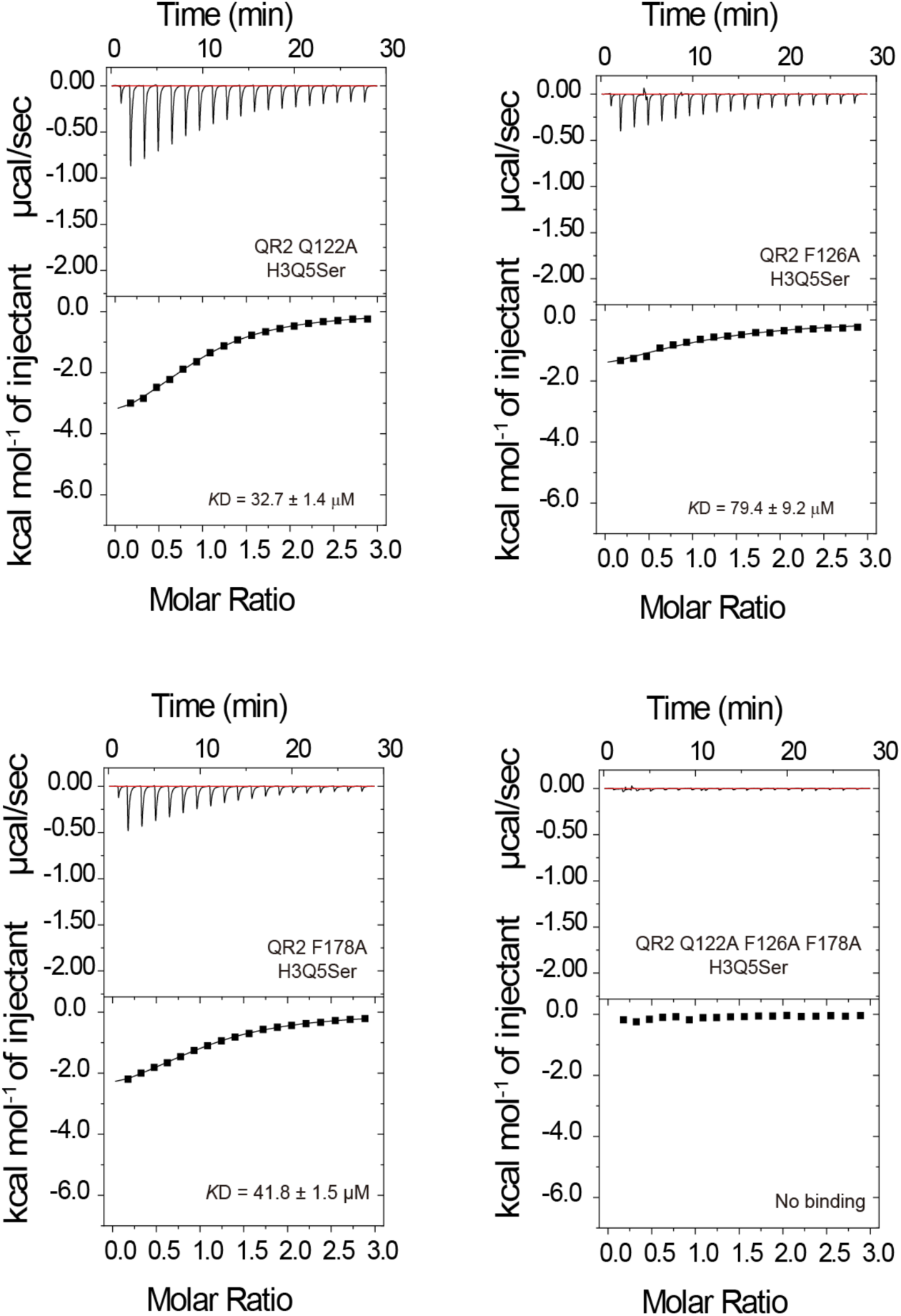
Mutational ITC validations of QR2’s molecular recognition of H3Q5ser, related to Figure 2. Titration and fitting curves of peptides titrated into mutant QR2. KD values are provided. See Table S1 for ITC statistics.

**Figure S5.**
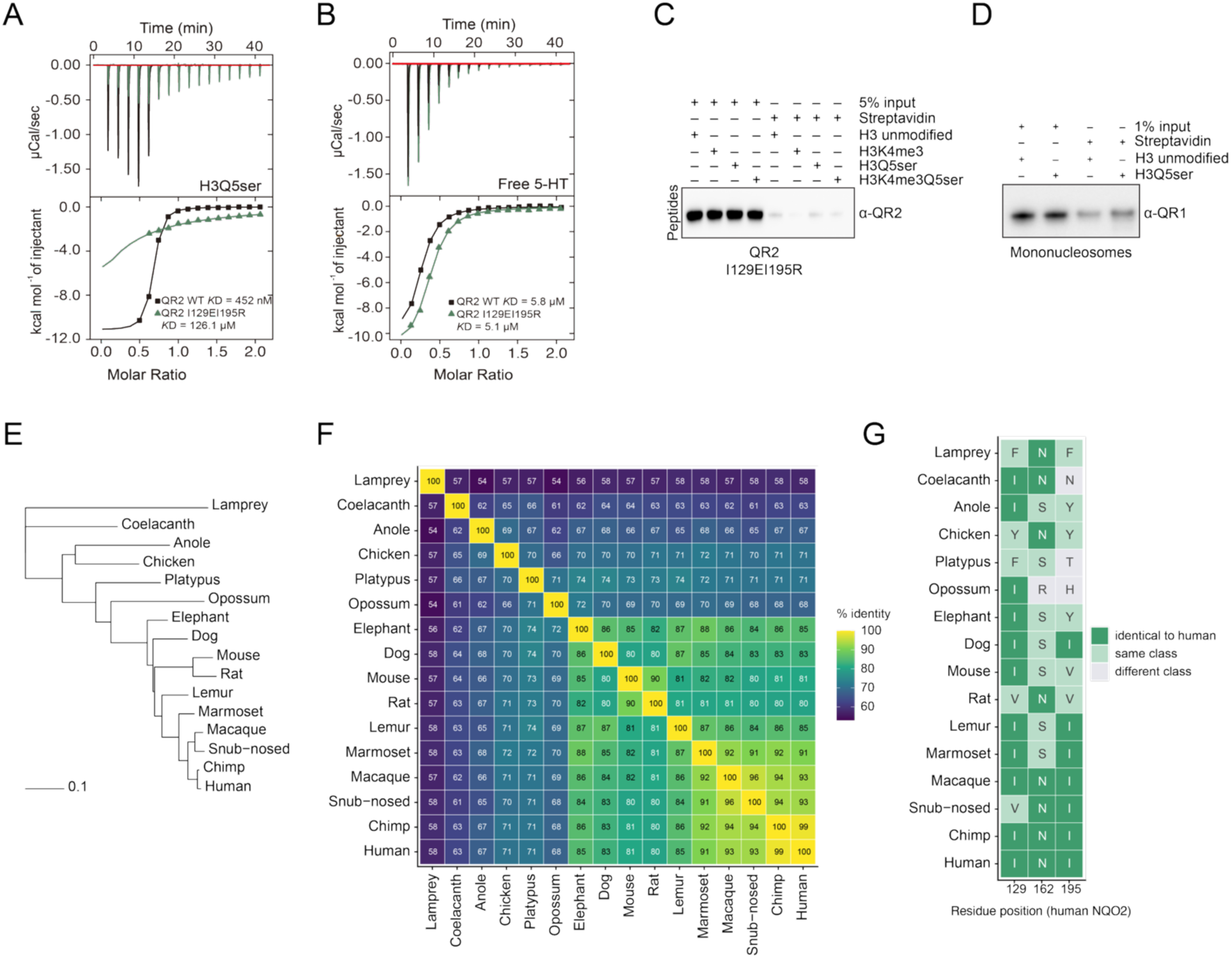
Validations of disruption of H3(K4me3)Q5ser recognition by QR2 I129EI195R mutations, related to Figure 2. (**A**) Titration and fitting curves of H3Q5ser peptides titrated into QR2 WT *vs.* mutant. *K*D values are provided. The mutants I129EI195R disrupts the recognition of QR2 to H3Q5ser peptides; (**B**) however, these mutations do not impact QR2’s binding to free 5-HT. See **Table S1** for ITC stats. (**C**) Peptide pulldowns with mutant recombinant QR2, followed by immunoblotting for QR2. (**D**) Mononucleosome pulldown validations of a lack of binding between QR1 and H3Q5ser. All immunoblotting experiments repeated 3X. See **Figure S14** for uncropped blots. **(E)** Maximum-likelihood phylogeny of NQO2 across 16 vertebrate species, from sea lamprey to human. Branch lengths were estimated under the LG+Γ model on the accepted species topology, held fixed, and the tree is rooted on *P. marinus*. Scale bar, 0.1 substitutions per site. **(F)** Pairwise amino-acid percent identity among all 16 NQO2 orthologues, computed over alignment columns at which both sequences carry a residue. Values range from 54% (lamprey versus anole and versus opossum) to 99% (human versus chimpanzee). **(G)** Residues at human NQO2 positions 129, 162 and 195 across the same 16 orthologues. Cells are coloured by whether the residue is identical to the human residue, of the same physicochemical class, or of a different class, with hydrophobic and aromatic residues grouped as one class. Position 129 is identical to human in 11 of 16 species and hydrophobic or aromatic in all 16; position 162 falls in the same class in 15 of 16; position 195 in 13 of 16.

**Figure S6.**
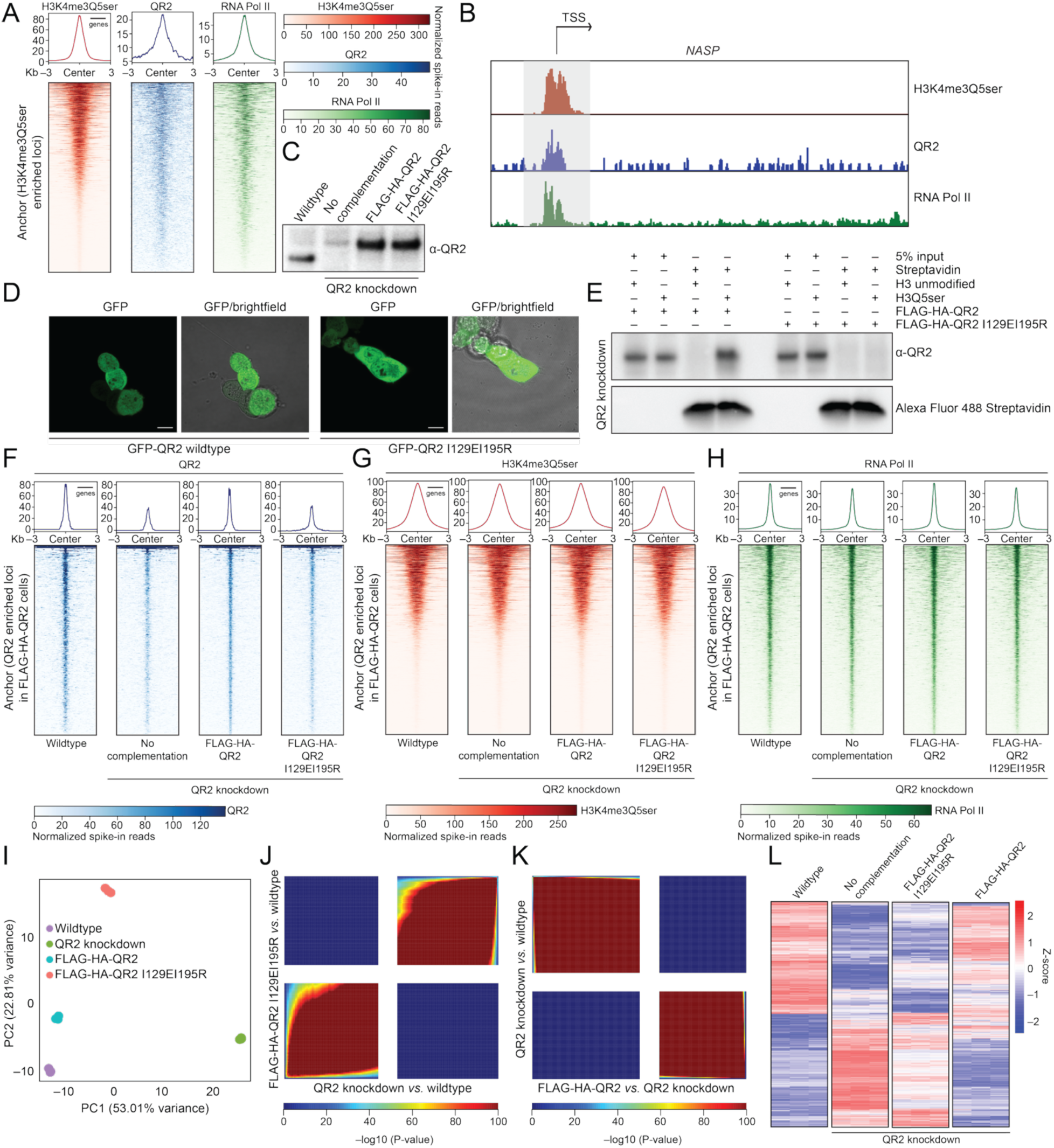
QR2 and H3Q5ser genomically co-enrich in HeLa cells to regulate gene expression, related to Figure 3. CUT&RUN-seq heatmaps of H3K4me3Q5ser, QR2, and RNA Pol II enrichment in (**A**) wildtype HeLa cells. (**B**) IGV tracks for H3K4me3Q5ser, QR2, and RNA Pol II at the *NASP* locus in wildtype HeLa cells. (**C**) Immunoblotting validation of endogenous QR2 knockdown in HeLa cells –/+ complementation with FLAG-HA-tagged wildtype *vs.* I129I195R QR2. (**D**) ICC/IF validation of QR2 subcellular localization in FLAG-HA-tagged wildtype *vs.* I129I195R QR2 complementation HeLa cells (*NQO2* knockdown background). Scale = 10 µm. (**E**) H3_1-18_Q5ser *vs.* H3_1-18_ unmodified peptide pulldowns from FLAG-HA-tagged wildtype *vs.* I129I195R QR2 complementation HeLa cells (*NQO2* knockdown background). CUT&RUN-seq heatmaps of (**F**) QR2, (**G**) H3K4me3Q5ser, and (**H**) RNA Pol II enrichment in wildtype, QR2 knockdown (no complementation), and FLAG-HA-tagged wildtype *vs.* I129I195R QR2 complementation HeLa cells (*NQO2* knockdown background). *n* = 3 for all. (**I**) RNA-seq-based PCA analysis of wildtype, QR2 knockdown (no complementation), and FLAG-HA-tagged wildtype *vs.* I129I195R QR2 complementation HeLa cells (*n* = 3 for all). RRHO analyses comparing gene expression patterns (RNA-seq) between (**J**) QR2 knockdown *vs.* wildtype and I129I195R QR2 complementation HeLa cells (*NQO2* knockdown background), as well as (**K**) QR2 knockdown *vs.* wildtype and QR2 wildtype complementation HeLa cells (*NQO2* knockdown background). (**L**) Heatmap of DEGs identified in QR2 knockdown *vs.* wildtype HeLa cells (anchor) across all four HeLa cell lines. All immunoblotting experiments repeated 3X. See **Figure S14** for uncropped blots.

**Figure S7.**
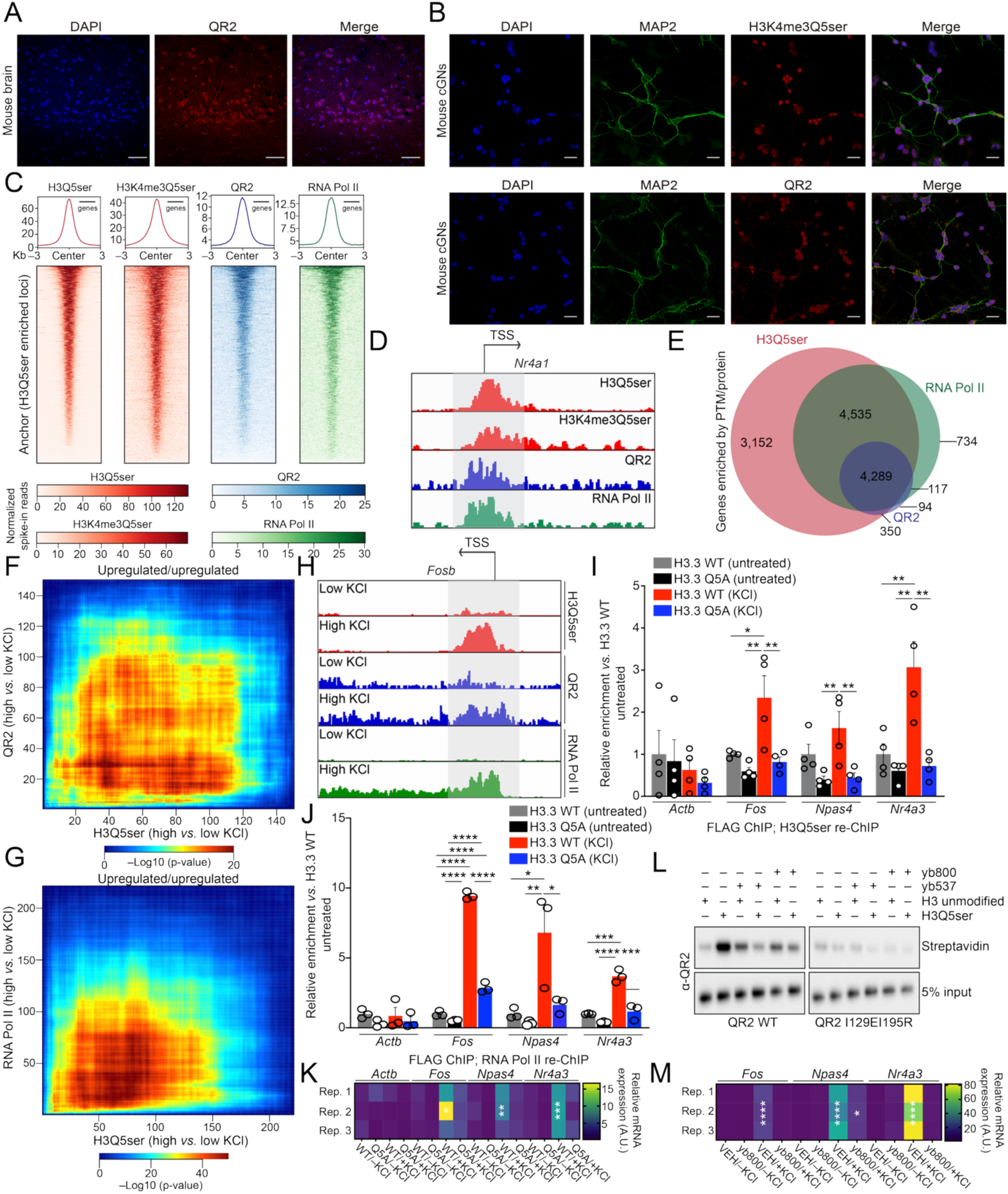
QR2 and H3Q5ser genomically co-enrich in rodent neurons and correlate with genes involved in synaptic plasticity, related to Figure 3. (**A**) IHC/IF validation of QR2 subcellular localization in wildtype mouse brain. Scale = 50 µm. (B) ICC/IF validation of QR2 subcellular localization in wildtype mouse cGNs. Scale = 20 µm. ChIP-seq heatmaps of H3(K4me3)Q5ser, QR2, and RNA Pol II enrichment in (**C**) wildtype mouse cGNs. (**D**) IGV tracks for H3Q5ser, QR2, and RNA Pol II at the *Nr4a1* locus in wildtype mouse cGNs. (**E**) Venn diagram of gene displaying overlapping enrichment for H3Q5ser, QR2, and RNA Pol II in wildtype mouse cGNs. RRHO analyses comparing (**F**) H3Q5ser enrichment in wildtype mouse cGNs –/+ KCl *vs.* QR2 enrichment in wildtype mouse cGNs –/+ KCl, and (**G**) H3Q5ser enrichment in wildtype mouse cGNs –/+ KCl *vs.* RNA Pol II enrichment in wildtype mouse cGNs –/+ KCl. (**H**) IGV tracks for H3Q5ser, QR2, and RNA Pol II at the *Fosb* locus in wildtype mouse cGNs –/+ KCl. (**I**) FLAG ChIP-/H3Q5ser re-ChIP-qPCRs following transduction of wildtype mouse cGNs –/+ KCl with H3.3 WT-FLAG *vs.* H3.3Q5A-FLAG vectors (*n* = 4; Two-way ANOVA, *P*<0.05 for effect of KCl, virus, or KCl x virus; post hoc Tukey’s MC tests – ** *P*<0.01, * *P*<0.05). (**J**) FLAG ChIP-/RNA Pol II re-ChIP-qPCRs following transduction of wildtype mouse cGNs –/+ KCl with H3.3 WT-FLAG *vs.* H3.3Q5A-FLAG vectors (*n* = 3; Two-way ANOVA, *P*<0.05 for effect of KCl, virus, or KCl x virus; post hoc Tukey’s MC tests – **** *P*<0.0001, *** *P*<0.001, ** *P*<0.01, * *P*<0.05). (**K**) qPCRs following transduction of wildtype mouse cGNs –/+ KCl with H3.3 WT-FLAG *vs.* H3.3Q5A-FLAG vectors [*n* = 3; Two-way ANOVA, *P*<0.05 for effect of KCl, virus, or KCl x virus; post hoc Dunnett’s MC tests (compared to –KCl H3.3 WT) – *** *P*<0.001, ** *P*<0.01). (**L**) H3_1-18_Q5ser *vs.* H3_1-18_ unmodified peptide pulldowns against recombinant QR2 in the presence or absence of QR2 inhibitors yb800 and yb537. (**M**) qPCRs following treatment of wildtype mouse cGNs –/+ KCl with vehicle *vs.* the QR2i yb800 [*n* = 3; Two-way ANOVA, *P*<0.05 for effect of KCl, QR2i, or KCl x QR2i; post hoc Uncorrected Fisher’s LSD (compared to –KCl vehicle) – **** *P*<0.0001, * *P*<0.05). Data shown as mean ± SEM. All immunoblotting experiments repeated 3X. See **Figure S14** for uncropped blots.

**Figure S8.**
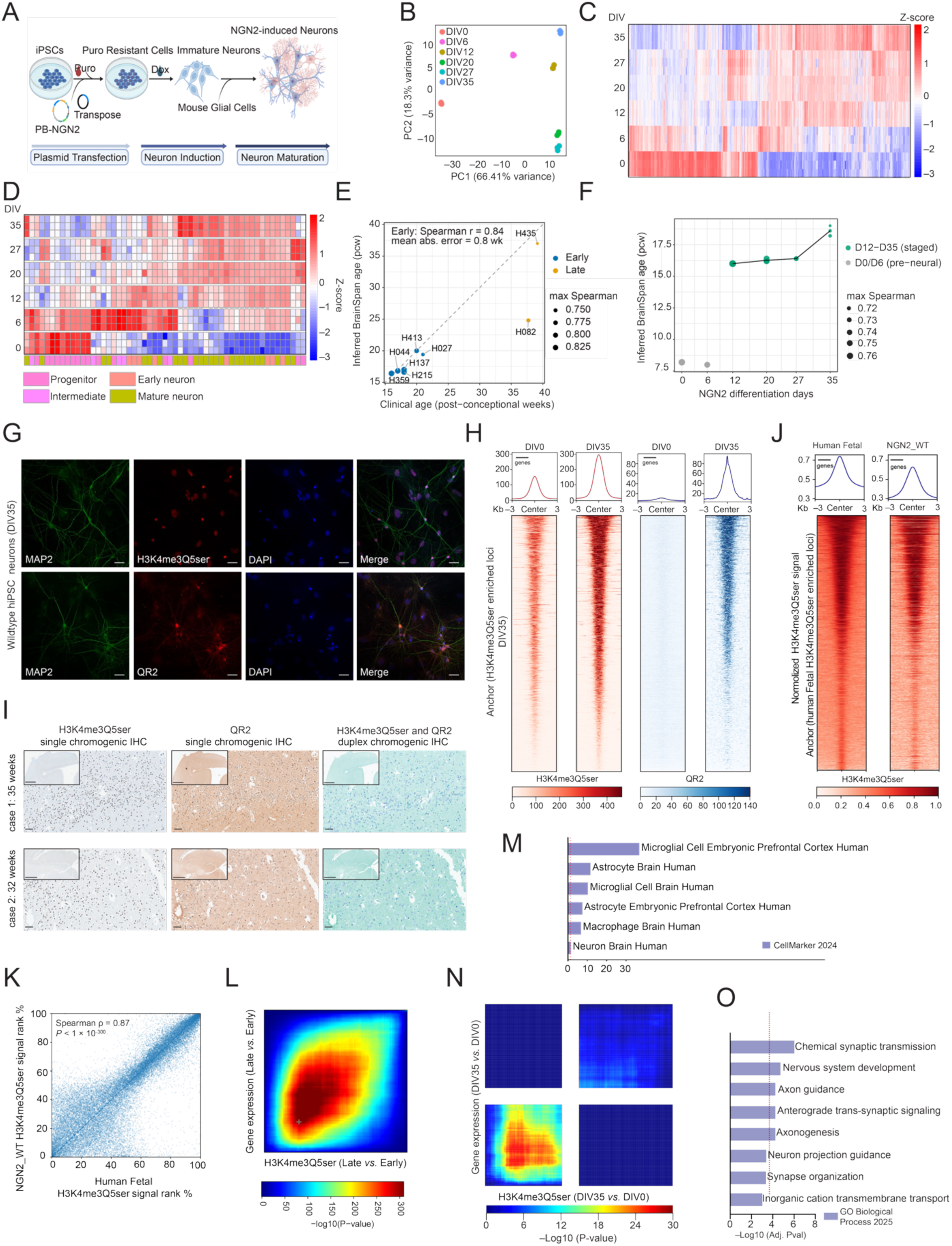
QR2-H3Q5ser dynamics in response to human neuronal differentiation and maturation, related to Figure 3. (**A**) Cartoon of NGN2-mediated induction of hiPSC neurons (images generated using Biorender). (**B**) RNA-seq-based PCA analysis of hiPSC neurons during differentiation and maturation from DIV0 to DIV 35. *n* = 3 for all. (**C**) Heatmaps comparing differential gene expression (anchor DIV35 *vs.* DIV0) across hiPSC neuronal differentiation and maturation. (**D**) Heatmaps comparing differential expression of marker genes (columns) for neural progenitors, early neurons, intermediate neurons, and fully matured neurons (anchor DIV35 *vs.* DIV0) across hiPSC neuronal differentiation and maturation. **(E)** Benchmarking of the BrainSpan staging procedure on human fetal cortical samples of known age. Each of eight post-mortem human fetal cortical samples was assigned a transcriptomic developmental age by correlating its expression of developmentally dynamic genes against the BrainSpan cortical RNA-seq reference; this inferred age (*y*, post-conceptional weeks, pcw) is plotted against the sample’s true clinical age (*x*, pcw = gestational age − 2), with the dashed line indicating identity (*y* = *x*). Points are colored by developmental window — Early (18-23 weeks, blue), Late (37-41 weeks, orange) — and sized by staging confidence (maximum Spearman correlation to the best-matching BrainSpan stage). Across the six Early samples, inferred and true ages agree closely (Spearman *r* = 0.84; mean absolute error = 0.8 weeks), validating the procedure. The two Late samples are postnatal infants (H435, 1 day old, born at 41 wk; H082, 19 days old, born at 37 wk) whose true developmental age falls in BrainSpan’s perinatal sampling gap; they are shown for completeness but excluded from the accuracy estimate. **(F)** Transcriptomic staging of the NGN2 differentiation time course by the identical, validated procedure. Inferred BrainSpan developmental age (y axis, pcw) across the WT NGN2 time course (differentiation days 0–35). Points are individual replicates (*n* = 3 per timepoint) and the connecting line traces the group means; point size denotes staging confidence (maximum Spearman correlation to BrainSpan). Pre-neural timepoints (D0/D6, grey) have no cortical counterpart and are shown but not staged; from neuronal commitment onward (D12–D35, teal) the cells map to a stable mid-second-trimester cortical age (∼16 pcw at D12–D27, rising modestly to ∼18–19 pcw by D35). (**G**) ICC/IF validation of QR2 subcellular localization in wildtype hiPSC neurons at DIV35. Scale = 50 µm. (**H**) CUT&RUN-seq heatmaps of H3K4me3Q5ser and QR2 enrichment in hiPSC neurons at DIV0 *vs.* DIV35. *n* = 3 for all. (**I**) Chromogenic immunohistochemistry of H3K4me3Q5ser and QR2 in human fetal cortex from two independent cases. H3K4me3Q5ser and QR2 were visualized individually by single-plex IHC (left and middle) and simultaneously by duplex chromogenic IHC (right), with H3K4me3Q5ser detected in purple and QR2 in green. Areas containing both signals exhibit a combined darker blue–purple appearance. Insets show low-magnification views of the corresponding cortical sections. Scale bars, 2 mm (insets) and 100 µm (main images). **(J)** Heatmap showing NGN2 WT neurons reproduce the genomic distribution of fetal H3K4me3Q5ser. Average enrichment profiles (top) and per-locus heatmaps (bottom) of H3K4me3Q5ser signal in human fetal cortex and NGN2 WT induced neurons, centered on human fetal H3K4me3Q5ser peaks (±3 kb; n = 32,453 peaks) and ordered by decreasing fetal enrichment, with the same order applied to both columns. Because the datasets were generated in independent batches with different absolute amplitudes, signal in each was log1p-transformed, scaled to its own 99th percentile of positive values, and capped at 1; color therefore reflects relative enrichment *within* each dataset, not directly comparable absolute magnitude. (**K**) Peak-level concordance of H3K4me3Q5ser between fetal cortex and NGN2 WT neurons. Each point is one fetal H3K4me3Q5ser peak, plotted by its within-dataset signal percentile in fetal cortex (x) and NGN2 WT neurons (y); dashed line, identity. Ranks were strongly correlated (Spearman ρ = 0.87, P < 1 × 10⁻³⁰⁰), and 86.3% of peaks in the top fetal quartile fell in the top NGN2 WT quartile — a 3.45-fold enrichment over the 25% expected by chance. **(L)** Developmental gain of promoter H3K4me3Q5ser in human fetal cortex tracks transcriptional up-regulation. The 11,164 genes carrying a H3K4me3Q5ser peak (≤3 kb from a TSS) and expressed at baseMean ≥ 50 were ranked independently by their Late-versus-Early change in H3K4me3Q5ser (x; DiffBind fold change) and in expression (y; DESeq2 signed −log10 P), from greatest increase at the origin to least; color is the −log10 hypergeometric P for the overlap of the top *i* and top *j* genes (Late = weeks 37, 41; Early = weeks 18–23). Overlap peaked at the top 2,756 × 2,438 genes (cross), where 1,351 genes were shared versus 602 expected (2.24-fold), exceeding all 2,000 gene-label permutations (P < 5 × 10⁻⁴); Spearman ρ = +0.463 between the two ranking statistics. **(M)** Enrichr (CellMarker 2024) cell-type enrichment of the 1,351 genes gaining both H3K4me3Q5ser and expression (Late versus Early), tested against all 11,164 H3K4me3Q5ser-marked, expressed genes as background. Bars, −log10 adjusted P; dotted line, adjusted P = 0.05; set/term overlap shown per bar. Microglial and astrocytic signatures — including stage- and tissue-matched embryonic prefrontal cortex references — were strongly enriched (adjusted P = 1.4 × 10⁻³⁷ to 2.5 × 10⁻¹²; 1.6–2.4-fold), whereas a generic neuronal signature was marginal (Neuron Brain Human, adjusted P = 0.034, 1.30-fold). (**N**) RRHO2 analysis comparing DEGs at DIV35 *vs.* DIV0 to H3K4me3Q5ser enrichment at DIV35 *vs.* DIV0 in hiPSC neurons. (**O**) Go Biological Process 2025 enrichment for genes displaying concordant induction of H3K4me3Q5ser and gene expression across hiPSC neuronal differentiation (FDR<0.05).

**Figure S9.**
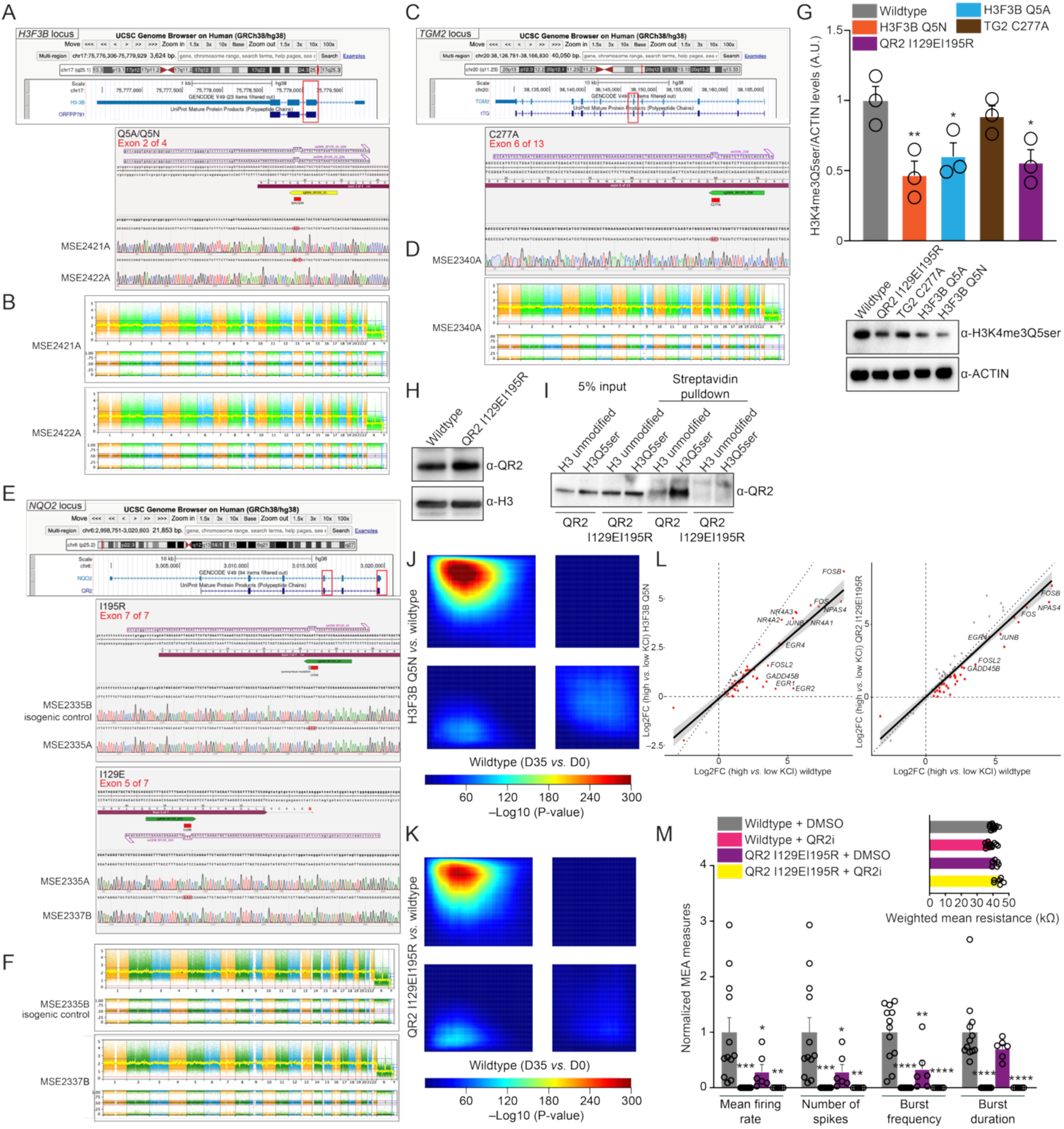
Generation and validation of mutant hiPSC lines, related to Figure 3. (**A**) Overview of the *H3F3B* genomic locus and Sanger sequencing confirming precise introduction of the Q5A and Q5N missense substitutions. Sequences of the sgRNA and the ssODN used to mediate homology-directed repair (HDR) are shown. Traces from the edited lines (MSE2421A and MSE2422A) are shown together with their isogenic controls, with edited nucleotides highlighted. (**B**) Whole-genome copy-number and allele-frequency plots indicating that both knock-in lines (MSE2421A and MSE2422A) maintain normal karyotypic profiles following genome engineering. (**C**) Overview of the *TGM2* genomic locus and Sanger sequencing confirming precise introduction of the C277A missense substitution in MSE2340A. The edited nucleotide is highlighted within the exon 6 sequence. (**D**) Whole-genome copy-number and allele-frequency plots indicating that the knock-in line (MSE2340A) maintains a normal karyotypic profile following genome engineering. (**E**) Overview of the *NQO2* genomic locus and Sanger sequencing confirming precise introduction of the I195R and I129E missense substitutions. Traces from edited lines are shown together with their isogenic controls, with edited nucleotides highlighted. (**F**) Whole-genome copy-number and allele-frequency plots indicating that both the knock-in line (MSE2337B) and its control maintain normal karyotypic profiles following genome engineering. (**G**) Immunoblotting validation of the impact of hiPSC mutations *vs.* wildtype on H3K4me3Q5ser levels in mature neurons (DIV35). *n* = 3/line. One-way ANOVA (F_4,10_ = 6.145, p=0.0092); post hoc Dunnett’s MC test – H3F3B Q5N (\*\**P*=0.0074), H3F3B Q5A (\**P*=0.0393), and QR2 I129EI195R (\**P*=0.0220). ACTIN was used as a loading control. (**H**) Immunoblotting validation that the QR2 I129EI195R mutations do not impact QR2 expression. (**I**) H3_1-18_Q5ser *vs.* H3_1-18_ unmodified peptide pulldowns from wildtype *vs.* QR2 I129EI195R hiPSC neurons at DIV35. RRHO analyses (RNA-seq) comparing the transcriptional impact of (**J**) H3F3B Q5N or (**K**) QR2 I129EI195R mutations in hiPSC neurons to normal patterns of gene expression during neuronal maturation from DIV0 to DIV35. (**L**) Plots of activity-dependent (KCl-induced) gene expression in wildtype hiPSC neurons (dashed line) *vs.* activity-dependent (KCl-induced) gene expression in H3F3B Q5N or QR2 I129EI195R lines, indicating attenuated responses in the mutants. (**M**) MEA-based quantification of: normalized mean firing rate [two-way ANOVA, main effect of QR2i: F_1,32_ = 10.79, *P*=0.0025; Dunnett’s MC test – *** *P*<0.001, ** *P*<0.01, * *P*<0.05]; normalized spike number [two-way ANOVA, main effect of QR2i: F_1,32_ = 10.79, *P*=0.0025; post hoc: Dunnett’s MC test – *** *P*<0.001, ** *P*<0.01, * *P*<0.05]; normalized burst frequency [two-way ANOVA, main effects of QR2i (F_1,32_ = 31.01, *P*<0.0001), mutation (F_1,32_ = 7.615, *P*=0.0095) and QR2i x mutation (F_1,32_ = 7.615, *P*=0.0095); post hoc: Dunnett’s MC test – **** *P*<0.0001, ** *P*<0.01]; and normalized burst duration [two-way ANOVA, main effect of QR2i: F_1,32_ = 46.96, *P*<0.0001; post hoc: Dunnett’s MC test – **** *P*<0.0001] in hiPSC wildtype (*n* = 12) or QR2 I129EI195R (*n* = 6) neurons –/+ the QR2i yb800. Inset: weighted mean resistance for each condition indicating no impact of genotype/treatment on neuronal viability.

**Figure S10.**
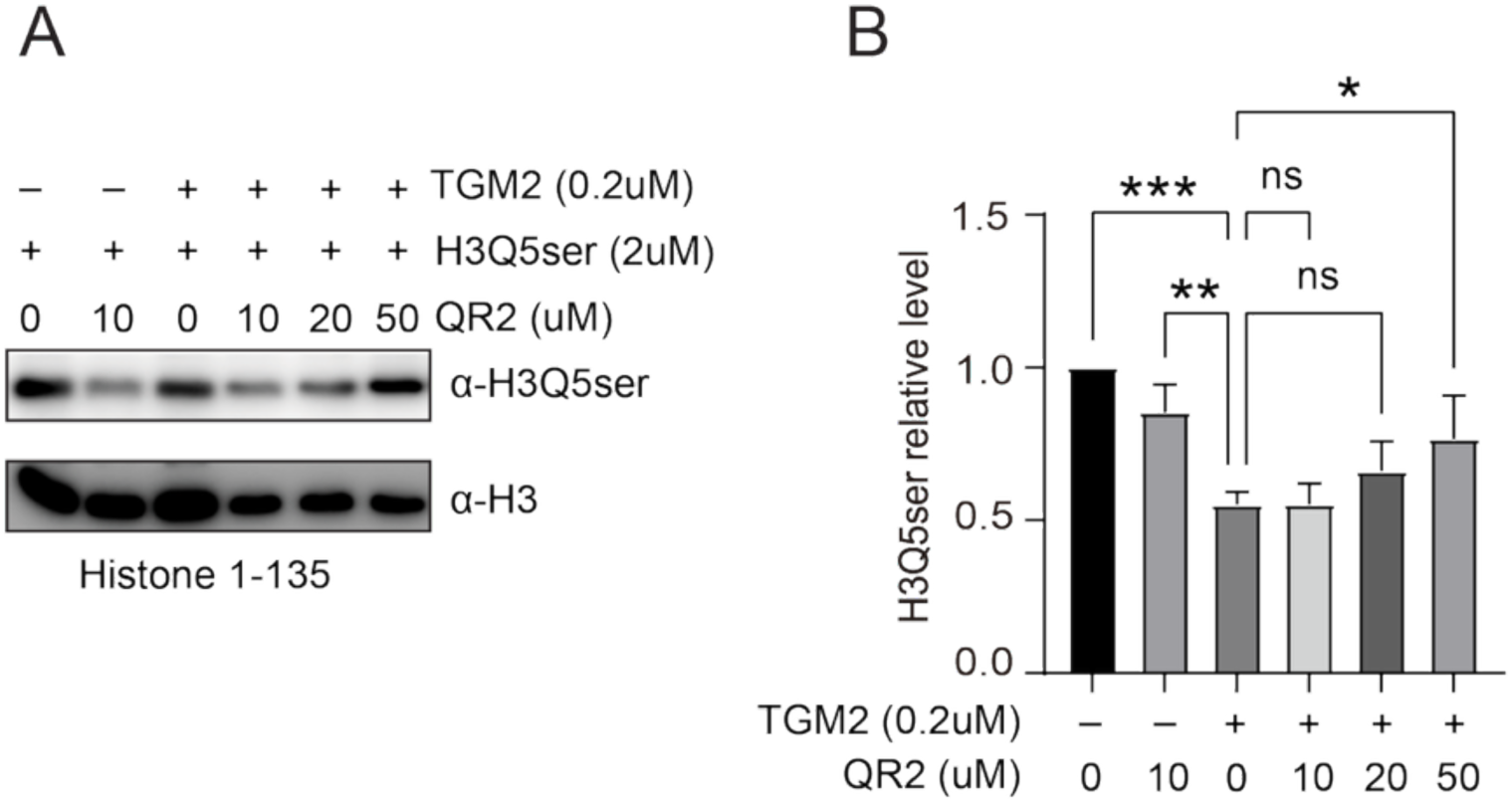
QR2 binding to H3Q5ser protects the mark from TGM2-mediated erasure, related to Figure 3. (**A**) *In vitro* erasure–protection assay. Recombinant histone H3 (residues 1–135) monoserotonylated at Gln5 (H3Q5ser; 2 µM) was incubated without or with the eraser TGM2 (0.2 µM) and with increasing concentrations of QR2 (0, 10, 20, 50 µM); residual mark was detected by immunoblot with an anti-H3Q5ser antibody, and total histone H3 (anti-H3) served as a loading control (left; lane conditions indicated above). (**B**) Quantification of H3Q5ser normalized to total H3 and expressed relative to the no-TGM2/no-QR2 condition (set to 1.0); bars are mean ± SD of n = 3 independent replicates. TGM2 markedly reduced H3Q5ser (versus the no-enzyme control; P = 0.0001, ***), whereas adding QR2 protected the mark dose-dependently, reaching significance at 50 µM (P = 0.0358, *; 10 and 20 µM, not significant); in the absence of TGM2, QR2 did not erase the mark, which remained significantly higher than the TGM2-only condition (P = 0.0039, **). All comparisons are versus the eraser-only condition (TGM2⁺/QR2 0) by one-way ANOVA; *P < 0.05, **P < 0.01, ***P < 0.001; ns, not significant. These results indicate that QR2 binding shields H3Q5ser from TGM2-catalyzed removal, consistent with a reader that protects the mark it engages.

**Figure S11.**
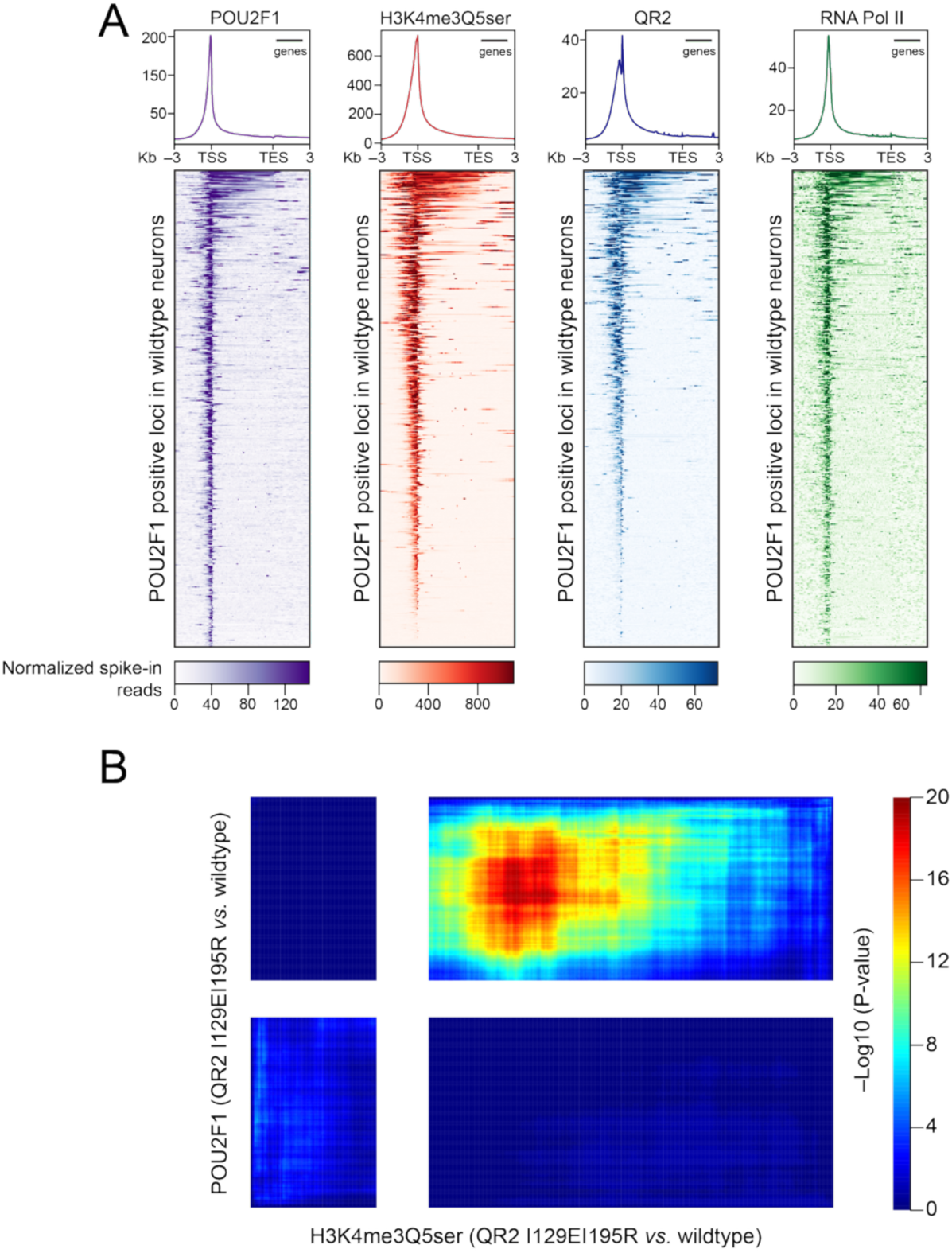
POU2F1 co-enriches with H3K4me3Q5ser and QR2 in hiPSC neurons, related to Figure 5. (**A**) CUT&RUN-seq heatmaps of POU2F1, H3K4me3Q5ser, QR2, and RNA Pol II enrichment (anchored on POU2F1 enriched genes) in wildtype hiPSC neurons at DIV35. *n* = 3 for all. (**B**) RRHO analyses (CUT&RUN-seq) comparing the impact of QR2 I129EI195R mutations on POU2F1 *vs.* H3K4me3Q5ser enrichment.

**Figure S12.**
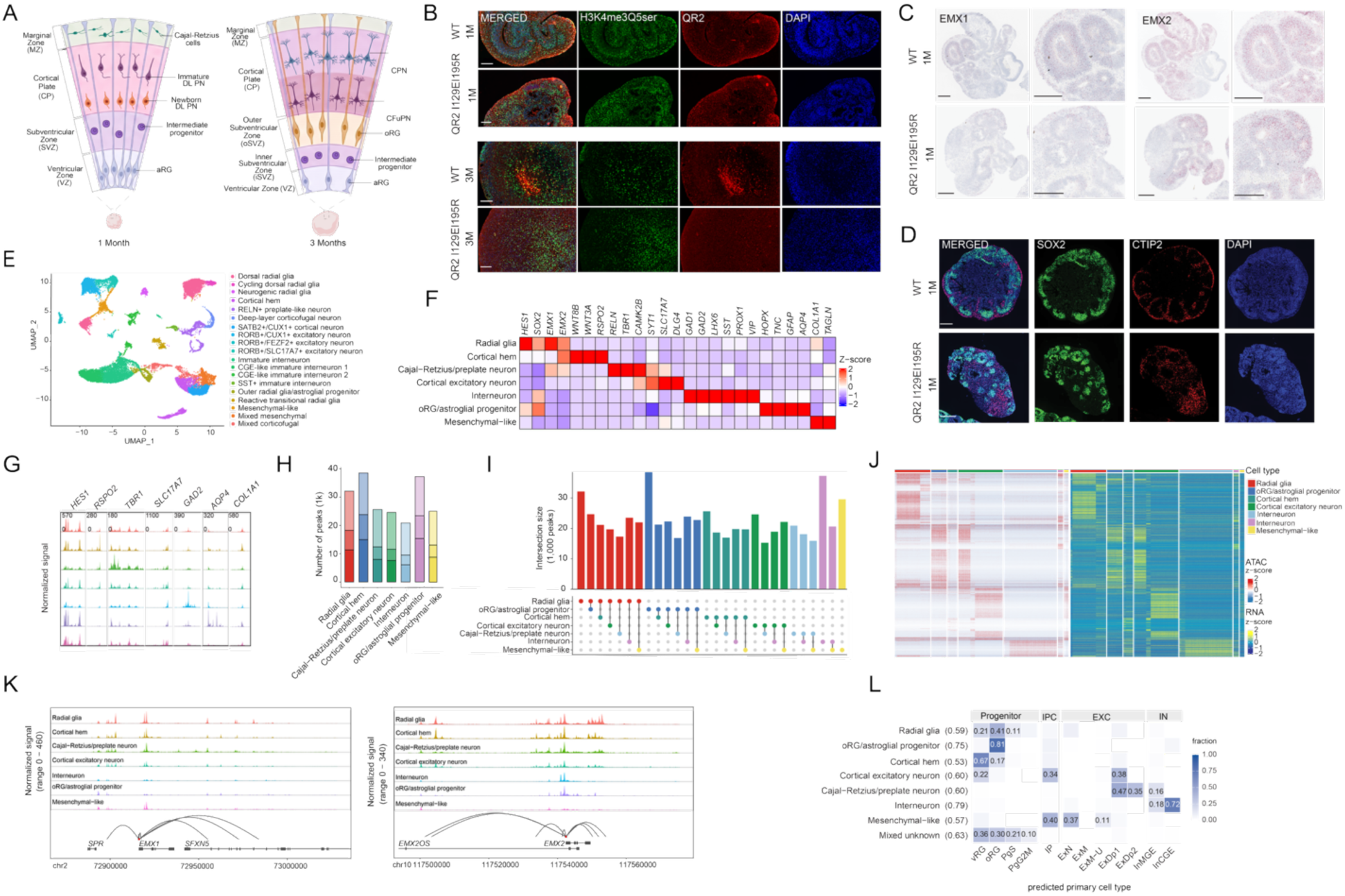
Single-nucleus multiome atlas of the cortical organoid: cell-type annotation, cell-type-specific chromatin accessibility concordant with expression, and fidelity to human fetal cortex, related to Figure 6. (**A**) Schematic of cell-type composition of dorsally patterned cortical organoids at one and three months, drawn from the single-cell and spatial transcriptomic atlas of this protocol^57^. At one month, apical radial glia (aRG) predominate, with intermediate progenitors (IP) above them and newborn and immature deep-layer projection neurons (DL PN) and Cajal–Retzius cells more superficially. By three months, outer radial glia (oRG) are present above the aRG, and corticofugal (CFuPN) and callosal (CPN) projection neurons occupy progressively more superficial positions. Immature DL PN and CFuPN are shown in the same color to indicate their shared deep-layer identity. Zones and cell morphologies are drawn by analogy with the developing human cortex and do not represent measured lamination in organoids; unspecified projection neurons, cortical hem cells and non-cortical cells are omitted. **(B)** Immunofluorescence for H3K4me3Q5ser (green) and QR2 (red) with DAPI (blue) in wild-type and QR2 I129EI195R organoids at one and three months; merged and single channels are shown for each of the four genotype-by-timepoint conditions. Whole organoids are shown at one month and fields within the organoid at three months. Both the mark and the reader are detectable in both genotypes at both timepoints. Scale bars, 100 μm. **(C)** RNAscope *in situ* hybridization for *EMX1* and *EMX2* in one-month wild-type and QR2 I129EI195R organoids; for each probe and genotype a whole-organoid view (left) and a higher-magnification view (right) are shown. Scale bars, 200 μm. **(D)** Immunofluorescence for SOX2 (green) and CTIP2 (red) with DAPI (blue) in one-month wild-type and QR2 I129EI195R organoids; merged and single channels are shown. Scale bars, 200 μm. **(E)** Weighted-nearest-neighbor (WNN) UMAP integrating the snRNA and snATAC modalities, colored by fine-grained cluster identity (key at right); nineteen clusters: dorsal, cycling-dorsal and neurogenic radial glia; cortical hem; RELN+ preplate-like and deep-layer corticofugal neurons; SATB2+/CUX1+ cortical neurons and RORB+/CUX1+, RORB+/FEZF2+ and RORB+/SLC17A7+ excitatory neurons; immature, CGE-like (1 and 2) and SST+ immature interneurons; outer radial glia/astroglial progenitor; reactive transitional radial glia; mesenchymal-like; and mixed mesenchymal and mixed corticofugal populations. **(F)** Canonical-marker heatmap validating the broad annotation: row-scaled mean expression (Z-score) of canonical markers (columns) across the seven broad cell types (rows) – radial glia (*HES1, SOX2, EMX1*), cortical hem (*EMX2, WNT8B, WNT3A, RSPO2*), Cajal–Retzius/preplate (*RELN, TBR1*), cortical excitatory (*CAMK2B, SYT1, SLC17A7, DLG4*), interneuron (*GAD1, GAD2, LHX6, SST, VIP, PROX1*), oRG/astroglial (*TNC, HOPX, GFAP, AQP4*), and mesenchymal-like (*COL1A1, TAGLN*). **(G)** Aggregated snATAC coverage at representative marker loci (*HES1, RSPO2, TBR1, SLC17A7, GAD2, AQP4, COL1A1*) across the seven broad cell types. Rows follow the cell-type order in (F) and are colored as in (J). Each locus is scaled independently to its own maximum, given at the top left of each column, so signal is comparable between cell types within a locus but not between loci. Accessibility tracks expression most clearly at *RSPO2* (cortical hem), *TBR1* (Cajal–Retzius/preplate), *GAD2* (interneuron) and *AQP4* (oRG/astroglial progenitor); other loci, including the broadly expressed progenitor gene *HES1*, retain accessibility across cell types, so identity is reflected in the relative magnitude of signal rather than in its presence or absence. **(H)** Number of ATAC peaks called per broad cell type (thousands), partitioned by genomic context (promoter, distal intronic/intergenic, gene body). Across all cell types most peaks are distal, consistent with cell-type-specific accessibility being carried predominantly by candidate enhancers. **(I)** UpSet plot of peak-set intersections across the seven cell types; bar height, intersection size (thousands of peaks); dot matrix, the contributing cell type(s) (single dot, cell-type-private peaks; connected dots, shared). The largest bars are cell-type-private sets, indicating each population contributes a substantial private accessible repertoire alongside a shared core. **(J)** Paired heatmaps of cell-type-specific features (rows grouped into cell-type blocks; top color bar and key at right, giving radial glia, oRG/astroglial progenitor, cortical hem, cortical excitatory neuron, Cajal–Retzius/preplate neuron, interneuron and mesenchymal-like): left, chromatin accessibility of cell-type-specific peaks (ATAC Z-score); right, expression of their linked genes (RNA Z-score). The block-diagonal structure in both panels shows that peaks accessible in a given cell type are matched by elevated linked-gene expression in the same cell type. Color scales, Z-score −2 to +2. **(K)** Aggregated snATAC coverage and Signac LinkPeaks peak-to-gene links at the dorsal-cortical transcription-factor loci *EMX1* (chr2:72,900,000– 73,000,000) and *EMX2* (chr10:117,500,000–117,560,000) across the seven broad cell types (group-normalized; y-axis range 0–460 for *EMX1*, 0–340 for *EMX2*). Arcs connect linked distal peaks to the target-gene transcription start site (|correlation| ≥ 0.1; gene models, EnsDb, GRCh38). Accessibility is dorsal-restricted – present in radial glia and the cortical hem, sustained in Cajal– Retzius/preplate and cortical excitatory neurons, and markedly lower in interneurons and mesenchymal-like cells – consistent with pallial identity. **(L)** Fidelity of organoid cell classes to primary fetal cortex. Organoid cells (single-nucleus multiome, RNA modality) were assigned to primary human fetal cortical cell types by label transfer onto the mid-gestation human neocortex atlas^59^ (Drop-seq, gestational weeks 17–18); oligodendrocyte precursor, microglial, endothelial and pericyte reference types, which are absent from one-to three-month organoids, were excluded. Each cell of the matrix gives the fraction of an organoid class (rows; mean prediction score in parentheses) assigned to each primary type (columns, grouped by lineage). Progenitor classes mapped to primary radial glia (outer radial glia/astroglial progenitor to oRG, 0.81 of cells; the broad radial glia class distributed across oRG 0.41, vRG 0.21 and PgS 0.11 rather than resolving to one of them), interneurons to primary interneurons (InCGE 0.72, InMGE 0.18), and cortical excitatory neurons to intermediate progenitors and deep-layer projection neurons (IP 0.34, ExDp1 0.38). The cortical hem and Cajal–Retzius/preplate populations have no dedicated counterpart in the mid-gestation atlas and were assigned to the nearest available classes – the hem to vRG (0.67), consistent with its ventricular-zone progenitor character, and Cajal–Retzius/preplate neurons to the deep-layer excitatory classes (ExDp1 0.47, ExDp2 0.35); both are shown for completeness and neither was used to claim excitatory or progenitor fidelity. The mesenchymal-like and mixed-unknown rows likewise map to no single primary type and were excluded from the lineage analyses. vRG, ventricular radial glia; oRG, outer radial glia; PgS and PgG2M, cycling progenitors; IP, intermediate progenitor; ExN, migrating excitatory neurons; ExM, maturing excitatory neurons; ExM-U, upper-layer-enriched maturing excitatory neurons; ExDp1 and ExDp2, deep-layer excitatory neurons; InMGE and InCGE, interneurons of medial and caudal ganglionic eminence origin.

**Figure S13.**
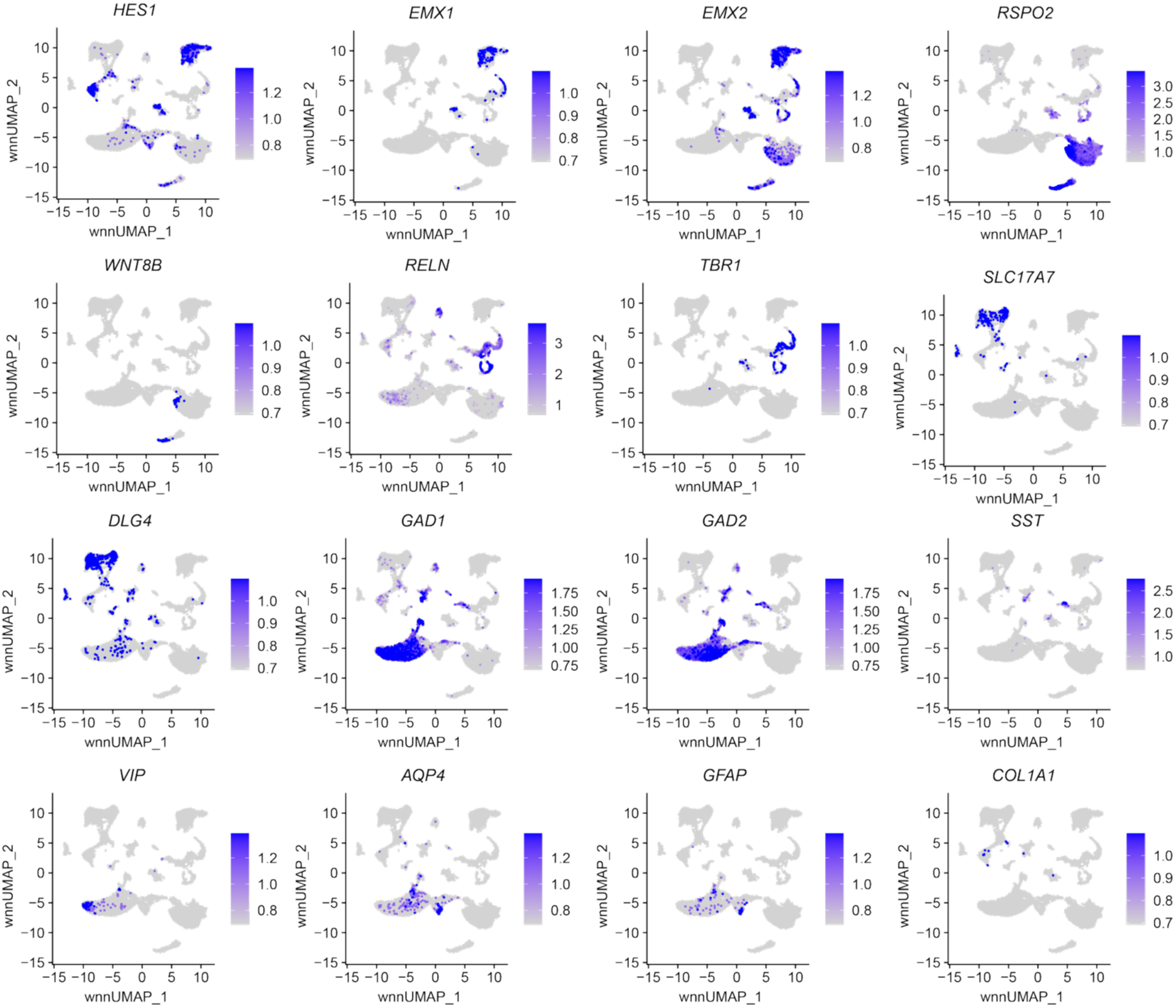
Canonical marker-gene expression validates the cell-type annotation of the cortical organoid, related to Figure 6. Feature plots of 16 canonical cell-type marker genes projected onto the weighted-nearest-neighbor (WNN) UMAP that integrates the single-nucleus RNA and ATAC modalities of the cortical organoid multiome, computed on all nuclei pooled across all samples (both genotypes and both ages). Each nucleus is colored by log-normalized snRNA expression of the indicated gene, from low (grey) to high (blue); color scales are independent per gene (per-panel color bar); axes are WNN UMAP dimensions 1 and 2. Markers label the expected dorsal-cortical populations: neural progenitors / radial glia (*HES1*); dorsal telencephalic (pallial) identity (*EMX1*, *EMX2*); cortical hem / WNT-high dorsomedial progenitors (*RSPO2*, *WNT8B*); Cajal–Retzius / preplate neurons (*RELN*); cortical excitatory neurons (*TBR1*, deep-layer/early-born; *SLC17A7*/VGLUT1; *DLG4*/PSD-95, postsynaptic); GABAergic inhibitory neurons (*GAD1*, *GAD2*) and their subtypes (*SST*, *VIP*); astroglia / astrocytes (*AQP4*, *GFAP*); and mesenchymal / non-neural cells (*COL1A1*). The spatially restricted, largely mutually exclusive expression of these markers confirms that the organoids generate the major dorsal cortical cell types.

**Figure S14:**
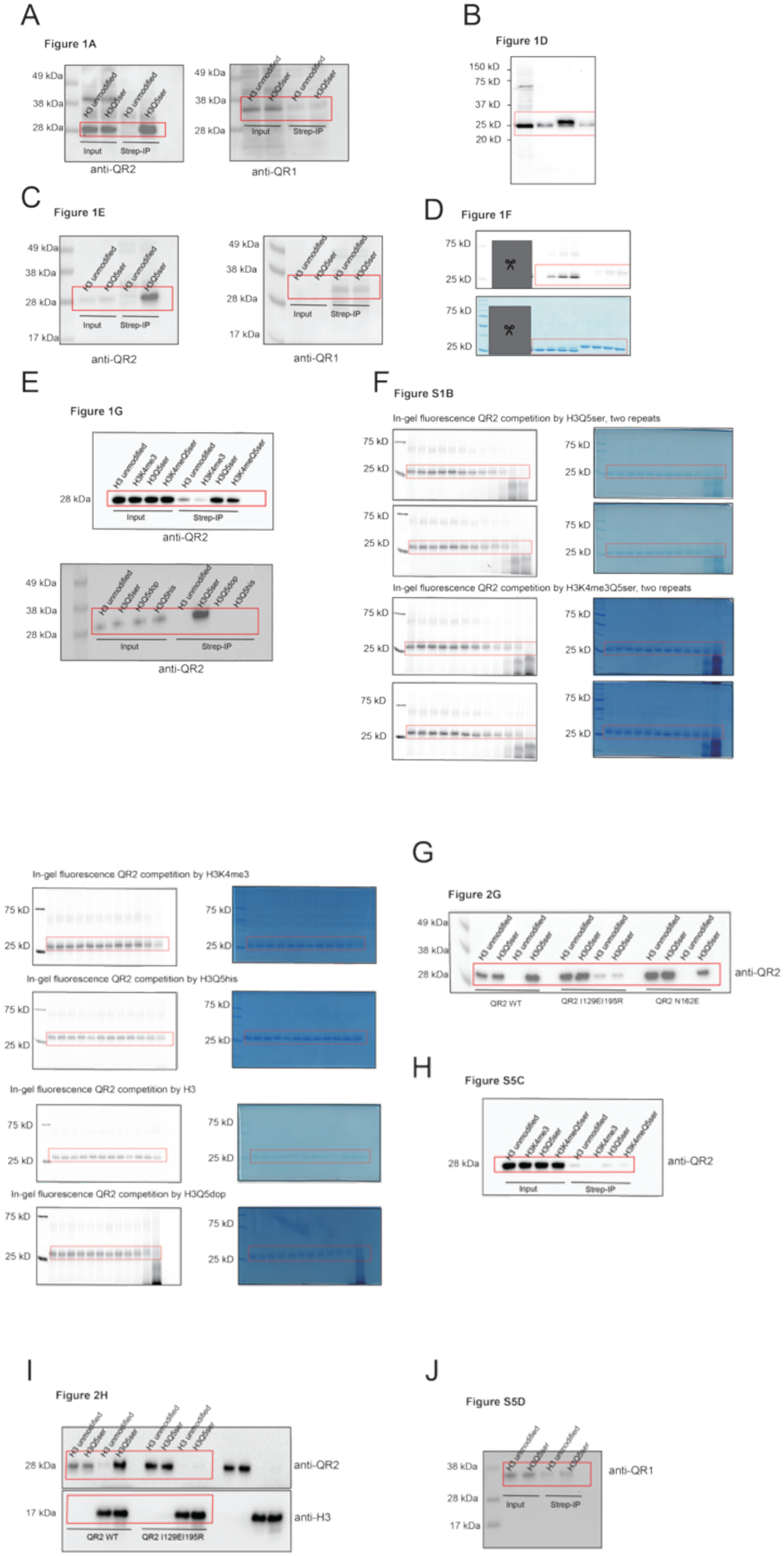
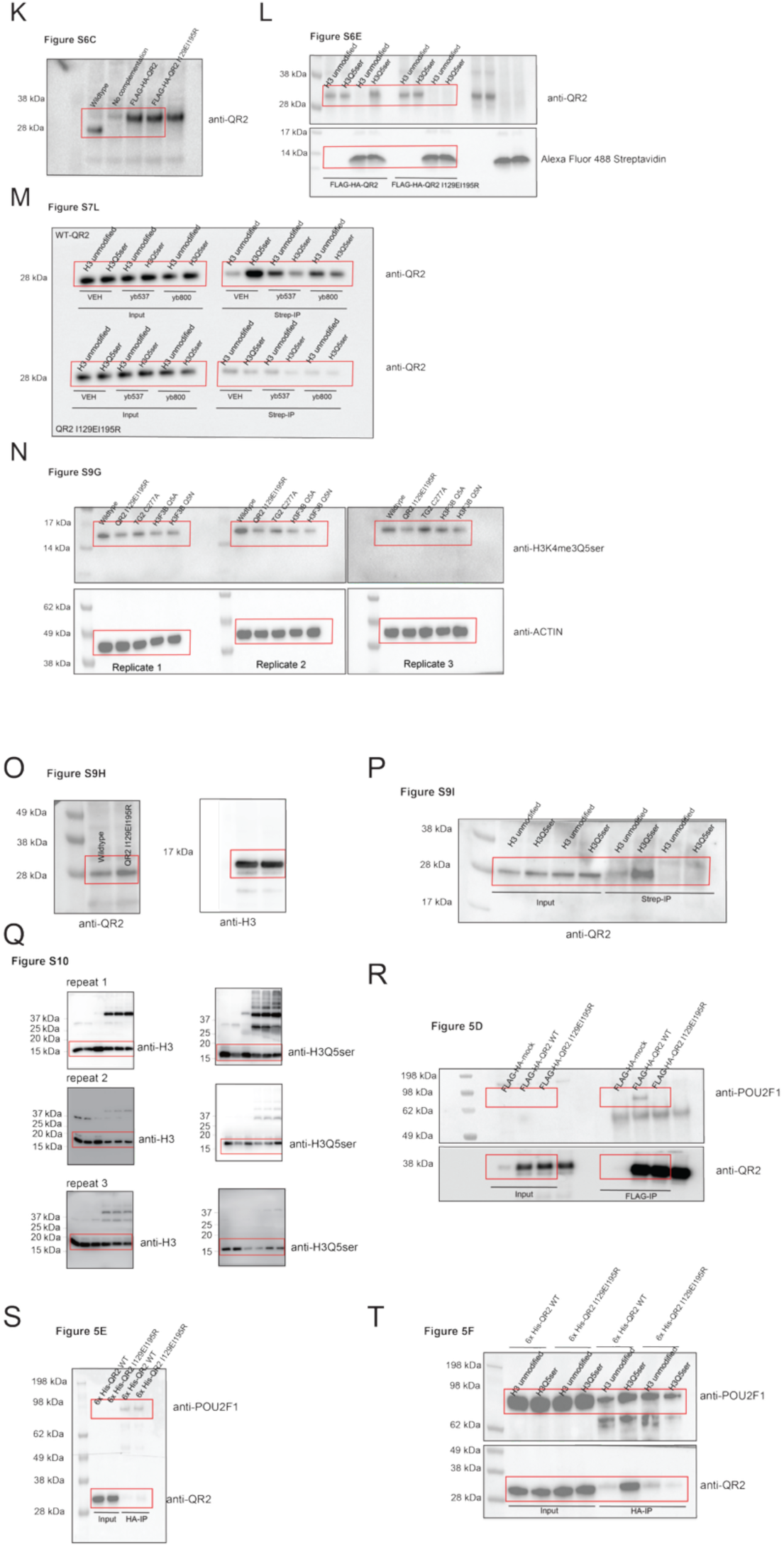
Uncropped immunoblots. Uncropped immunoblots related to (**A**) Figure 1A, (**B**) Figure 1D, (**C**) Figure 1E, (**D**) Figure 1F, (**E**) Figure 1G, (**F**) Figure S1B, (**G**) Figure 2G, (**H**) Figure S5C, (**I**) Fig. 2H, (**J**) Figure S5D, (**K**) Figure S6C, (**L**) Figure S6E, (**M**) Figure S7L, (**N**) Figure S9G, (**O**) Figure S9H, (**P**) Figure S9I, (**Q**) Figure S10, (**R**) Figure 5D, (**S**) Figure 5E, and (**T**) Figure 5F.

**Table S1:** ITC statistics.

| NO. | Proteins | Ligands | $\Delta H$ (kcal/mol) | $\Delta S$ (cal/mol/deg) | KD | N value | Relevant Figure Panel |
| --- | --- | --- | --- | --- | --- | --- | --- |
| 1 | QR2 | H3(1-10) | — | — | No binding detected | — | Figure 1H |
| 2 | QR2 | H3(1-10)K4me3 | — | — | No binding detected | — | Figure 1H |
| 3 | QR2 | H3(1-10)Q5ser | $-7.31 \pm 0.05$ | 0.726 | $3.08 \pm 0.15 \mu\text{M}$ | $0.840 \pm 0.004$ | Figure 1H |
| 4 | QR2 | H3(1-10)K4me3Q5ser | $-5.83 \pm 0.08$ | 5.71 | $2.99 \pm 0.24 \mu\text{M}$ | $1.000 \pm 0.007$ | Figure 1H |
| 5 | QR2 | H3(1-15)Q5his | — | — | No binding detected | — | Figure 1I |
| 6 | QR2 | H3(1-15)Q5dop | — | — | No binding detected | — | Figure 1I |
| 7 | QR2 (FAD-binding) | H3(1-10)Q5ser | $-6.65 \pm 0.06$ | 4.18 | $1.63 \pm 0.16 \mu\text{M}$ | $1.04 \pm 0.01$ | Figure S2C |
| 8 | QR2 (apo) | H3(1-10)Q5ser | — | — | No binding detected | — | Figure S2C |
| 9 | QR2-Q122A | H3(1-10)Q5ser | $-4.31 \pm 0.08$ | 6.07 | $32.7 \pm 1.4 \mu\text{M}$ | $0.93 \pm 0.01$ | Figure S4 |
| 10 | QR2-F126A | H3(1-10)Q5ser | $-2.51 \pm 0.11$ | 10.4 | $79.4 \pm 9.2 \mu\text{M}$ | $1.00 \pm 0.00$ | Figure S4 |
| 11 | QR2-F178A | H3(1-10)Q5ser | $-3.21 \pm 0.05$ | 9.27 | $41.8 \pm 1.5 \mu\text{M}$ | $1.03 \pm 0.01$ | Figure S4 |
| 12 | QR2-Q122A/F126A/F178A | H3(1-10)Q5ser | — | — | No binding detected | — | Figure S4 |
| 13 | QR2 (WT) | H3(1-10)Q5ser | $-11.16 \pm 0.07$ | -8.41 | $452 \pm 21 \text{ nM}$ | $0.626 \pm 0.001$ | Figure S5A |
| 14 | QR2 (WT) | 5-HT | $-11.18 \pm 0.24$ | -13.5 | $5.78 \pm 0.29 \mu\text{M}$ | $0.233 \pm 0.004$ | Figure S5B |
| 15 | QR2 I129E I195R | H3(1-10)Q5ser | n.d. | n.d. | $126.1 \pm 45.3 \mu\text{M}$ | n.d. | Figure S5A |
| 16 | QR2 I129E I195R | 5-HT | $-11.53 \pm 0.27$ | -14.4 | $5.08 \pm 0.44 \mu\text{M}$ | $0.362 \pm 0.006$ | Figure S5B |

**Table S2:** X-ray crystallography data collection and refinement statistics, related to Figures 2A–2D, S2A, S2B and S2D–S2G.

| Crystal | QR2-H3Q5ser | QR2-H3K4me3Q5ser |
| --- | --- | --- |
| **Data collection** |  |  |
| Wavelength(Å) | 0.9792 | 0.9792 |
| Space group | P4<sub>3</sub>2<sub>1</sub>2 | P4<sub>3</sub>2<sub>1</sub>2 |
| Cell dimensions |  |  |
| a, b, c (Å) | 119.0, 119.0, 149.8 | 119.9, 119.9, 150.2 |
| α, β, γ (°) | 90, 90, 90 | 90, 90, 90 |
| Resolution (Å) | 50 - 2.40 (2.44 - 2.40) | 50 - 2.59 (2.63 - 2.59) |
| No. of reflections | 42961 (2094) | 34824 (1749) |
| R<sub>merge</sub> (%) | 20.0 (98.3) | 21.1 (120.6) |
| R<sub>pim</sub> (%) | 4.0 (20.0) | 7.1 (42.7) |
| I/σ (I) | 20.0 (3.0) | 14.3 (2.6) |
| CC1/2 (%) | 99.5 (89.4) | 99.1 (77.5) |
| Completeness (%) | 100.0 (100.0) | 99.6 (100.0) |
| Redundancy | 25.8 (24.8) | 9.6 (8.4) |
| **Refinement (F>0)** |  |  |
| R<sub>work</sub>/R<sub>free</sub> (%) | 16.3/21.2 | 15.5/20.2 |
| No. of non-H atoms |  |  |
| Protein | 7290 | 7285 |
| Peptide | 300 | 300 |
| Water | 472 | 315 |
| Average B-factor (Å<sup>2</sup>) |  |  |
| Protein | 40.1 | 36.5 |
| Peptide | 38.9 | 33.7 |
| Water | 38.7 | 32.8 |
| RMSD bonds (Å) | 0.003 | 0.005 |
| RMSD angels (°) | 0.595 | 0.806 |
| **Ramachandran plot** |  |  |
| Favored (%) | 96.01 | 97.13 |
| Allowed (%) | 3.99 | 2.87 |
| Outliers (%) | 0.00 | 0.00 |

**Table S3.**
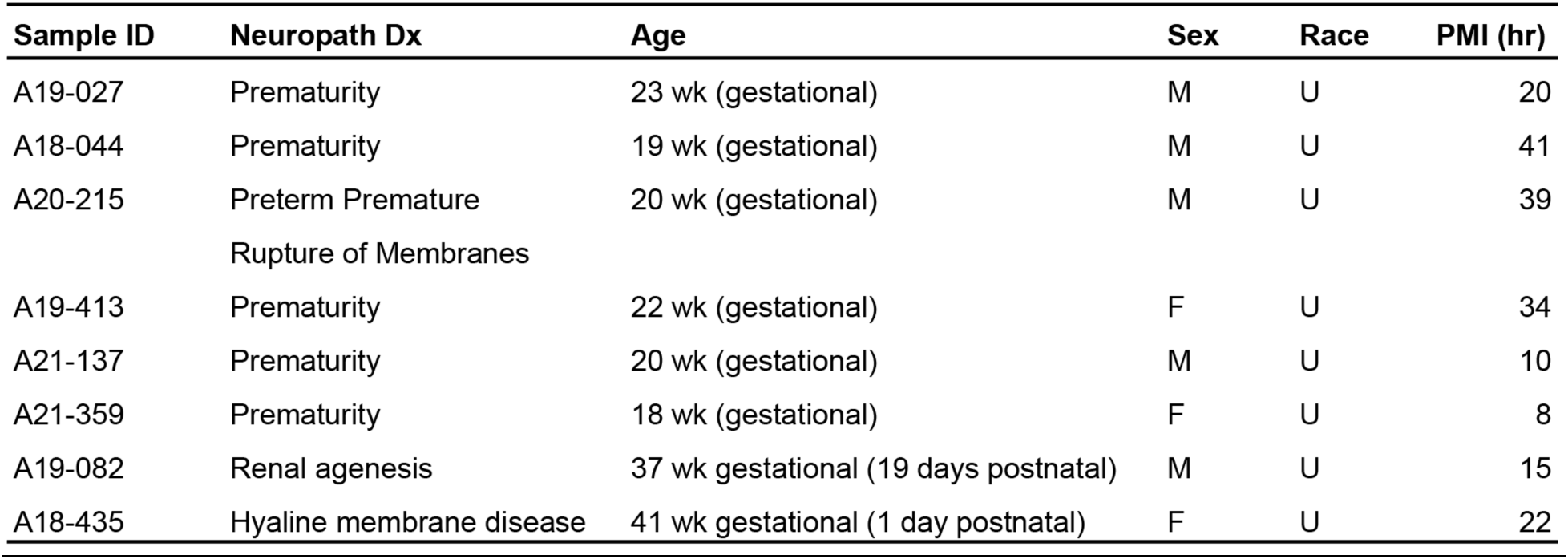
Human postmortem brain cases used for developing human cortex RNA-seq and CUT&RUN-seq, related to Figures 3A, 3H, S8E and S8J–S8M.

**Data S1:** LC-MS/MS analysis from HeLa cell Nes, related to Figure 1A

**Data S2:** LC-MS/MS analysis from SK-N-SH cell lysates, related to Figure 1C

**Data S3:** NQO2 orthologs, elated to Figures S5E–S5G

**Data S4:** CUT&RUN-seq from wildtype HeLa cells (H3K4me3Q5ser, QR2, RNA Pol II), related to Figures S6A and S6B

**Data S5:** CUT&RUN-seq from wildtype *vs.* complementation HeLa cells (H3K4me3Q5ser, QR2, RNA Pol II), related to Figures S6F–S6H

**Data S6:** RNA-seq from wildtype *vs.* complementation HeLa cells, related to Figures S6I–S6L

**Data S7:** ChIP-seq from mouse cGNs −/+ KCl (H3K4me3Q5ser, H3Q5ser, QR2, RNA PolII), related to Figures S7C–S7H

**Data S8:** RNA-seq from hiPSC neurons (time course), related to Figures S8B–S8D

**Data S9:** CUT&RUN-seq from hiPSC neurons (DIV0/DIV35; H3K4me3Q5ser, QR2), related to Figure S8H

**Data S10:** RNA-seq from developing human cortex, related to Figures S8E and S8L

**Data S11:** CUT&RUN-seq from developing human cortex (H3K4me3Q5ser, QR2), related to Figures 3A and S8J–S8L

**Data S12:** Cell-type enrichment of genes gaining both H3K4me3Q5ser and expression across fetal cortical development, related to Figure S8M

**Data S13:** Gene ontologies related to Figure S8O

**Data S14:** CUT&RUN-seq from wildtype *vs.* mutant hiPSC neurons (H3K4me3Q5ser, QR2, RNA Pol II), related to Figures 3B–3G

**Data S15:** RNA-seq from wildtype *vs.* mutant hiPSC neurons, related to Figures 3H–3L and S9J–S9K

**Data S16:** 1,193 Overlapping genes displaying regulation in mutant *vs.* wildtype hiPSC neurons, related to Figures 3M–3O

**Data S17:** Gene ontologies, related to Figure 3P

**Data S18:** LC-MS/MS analysis from HeLa NEs (FLAG-HA-tagged QR2 wildtype *vs.* Flag-HA mock; FLAG-HA-tagged QR2 I129EI195R *vs.* FLAG-HA-tagged QR2 wildtype), related to Figures 5B and 5C

**Data S19:** CUT&RUN-seq from wildtype *vs.* mutant hiPSC neurons (POU2F1), related to Figures 5G, S11A and S11B

**Data S20:** ATAC-seq from wildtype *vs.* mutant hiPSC neurons, related to Figures 5H and 5I

**Data S21:** Gene ontologies related to Figure 5J

**Data S22:** GSEA of radial glia pseudobulk profiles, related to Figure 6E and 6F

**Data S23:** POU2F1-linked peak–gene pairs in cortical excitatory neurons, related to Figures 6G and 6H

## References

1. Dincer, E., Oguz, M., and Baris, Y. (2018). Biochemical and pharmacological properties of biogenic amines. Biogenic Amines. 10.5772/intechopen.81569.

2. Al-Kachak, A., and Maze, I. (2023). Post-translational modifications of histone proteins by monoamine neurotransmitters. Current Opinion in Chemical Biology 74, 102302. 10.1016/j.cbpa.2023.102302.

3. Fukumoto, T., Kema, I.P., and Levin, M. (2005). Serotonin signaling is a very early step in patterning of the left-right axis in chick and frog embryos. Current Biology 15, 794–803. 10.1016/j.cub.2005.03.044.

4. Neumann, J., Hofmann, B., Dhein, S., and Gergs, U. (2023). Cardiac roles of serotonin (5-HT) and 5-HT-receptors in health and disease. Int J Mol Sci 24. 10.3390/ijms24054765.

5. Cirillo, C., Vanden Berghe, P., and Tack, J. (2011). Role of serotonin in gastrointestinal physiology and pathology. Minerva Endocrinol 36, 311–324.

6. Teleanu, R.I., Niculescu, A.G., Roza, E., Vladâcenco, O., Grumezescu, A.M., and Teleanu, D.M. (2022). Neurotransmitters-key factors in neurological and neurodegenerative disorders of the central nervous system. Int J Mol Sci 23. 10.3390/ijms23115954.

7. Yang, X., Lou, J., Shan, W., Ding, J., Jin, Z., Hu, Y., Du, Q., Liao, Q., Xie, R., and Xu, J. (2021). Pathophysiologic role of neurotransmitters in digestive diseases. Front Physiol 12, 567650. 10.3389/fphys.2021.567650.

8. Walther, D.J., Stahlberg, S., and Vowinckel, J. (2011). Novel roles for biogenic monoamines: from monoamines in transglutaminase-mediated post-translational protein modification to monoaminylation deregulation diseases. FEBS J 278, 4740–4755. 10.1111/j.1742-4658.2011.08347.x.

9. Walther, D.J., Peter, J.U., Winter, S., Höltje, M., Paulmann, N., Grohmann, M., Vowinckel, J., Alamo-Bethencourt, V., Wilhelm, C.S., Ahnert-Hilger, G., and Bader, M. (2003). Serotonylation of small GTPases is a signal transduction pathway that triggers platelet alpha-granule release. Cell 115, 851–862. 10.1016/s0092-8674(03)01014-6.

10. Lepack, A.E., Werner, C.T., Stewart, A.F., Fulton, S.L., Zhong, P., Farrelly, L.A., Smith, A.C.W., Ramakrishnan, A., Lyu, Y., Bastle, R.M., et al. (2020). Dopaminylation of histone H3 in ventral tegmental area regulates cocaine seeking. Science 368, 197–201. 10.1126/science.aaw8806.

11. Stewart, A.F., Lepack, A.E., Fulton, S.L., Safovich, P., and Maze, I. (2023). Histone H3 dopaminylation in nucleus accumbens, but not medial prefrontal cortex, contributes to cocaine-seeking following prolonged abstinence. Mol Cell Neurosci 125, 103824. 10.1016/j.mcn.2023.103824.

12. Fulton, S.L., Mitra, S., Lepack, A.E., Martin, J.A., Stewart, A.F., Converse, J., Hochstetler, M., Dietz, D.M., and Maze, I. (2022). Histone H3 dopaminylation in ventral tegmental area underlies heroin-induced transcriptional and behavioral plasticity in male rats. Neuropsychopharmacology 47, 1776–1783. 10.1038/s41386-022-01279-4.

13. Zheng, Q., Weekley, B.H., Vinson, D.A., Zhao, S., Bastle, R.M., Thompson, R.E., Stransky, S., Ramakrishnan, A., Cunningham, A.M., Dutta, S., et al. (2025). Bidirectional histone monoaminylation dynamics regulate neural rhythmicity. Nature 637, 974–982. 10.1038/s41586-024-08371-3.

14. Farrelly, L.A., Thompson, R.E., Zhao, S., Lepack, A.E., Lyu, Y., Bhanu, N.V., Zhang, B., Loh, Y.-H.E., Ramakrishnan, A., Vadodaria, K.C., et al. (2019). Histone serotonylation is a permissive modification that enhances TFIID binding to H3K4me3. Nature 567, 535–539. 10.1038/s41586-019-1024-7.

15. Zhao, S., Chuh, K.N., Zhang, B., Dul, B.E., Thompson, R.E., Farrelly, L.A., Liu, X., Xu, N., Xue, Y., Roeder, R.G., et al. (2021). Histone H3Q5 serotonylation stabilizes H3K4 methylation and potentiates its readout. Proc Natl Acad Sci U S A 118. 10.1073/pnas.2016742118.

16. Lukasak, B.J., Mitchener, M.M., Kong, L., Dul, B.E., Lazarus, C.D., Ramakrishnan, A., Ni, J., Shen, L., Maze, I., and Muir, T.W. (2022). TGM2-mediated histone transglutamination is dictated by steric accessibility. Proc Natl Acad Sci U S A 119, e2208672119. 10.1073/pnas.2208672119.

17. Pulido-Cortés, L., Gielingh, H., Thijssen, V., Liu, M., Yoshisada, R., Romão Soares, L., Nizamuddin, S., Friedrich, F., Greschik, H., Peng, L., et al. (2025). Molecular determinants for recognition of serotonylated chromatin. Nucleic Acids Res 53. 10.1093/nar/gkaf612.

18. Zhao, J., Chen, W., Pan, Y., Zhang, Y., Sun, H., Wang, H., Yang, F., Liu, Y., Shen, N., Zhang, X., et al. (2021). Structural insights into the recognition of histone H3Q5 serotonylation by WDR5. Sci Adv 7. 10.1126/sciadv.abf4291.

19. O’Chan, J.C., Weekley, B.H., Yang, C., Chen, M., Cunningham, A.M., Dutta, S., Chen, W., Iyer, R.R., Ahmed, N.I., Cetin, B., et al. (2026). KMT2E recruitment by H3 serotonylation regulates neurodevelopmental chromatin dynamics. bioRxiv, 2026.2009.2020.752984. 10.64898/2026.09.20.752984.

20. Sardar, D., Cheng, Y.T., Woo, J., Choi, D.J., Lee, Z.F., Kwon, W., Chen, H.C., Lozzi, B., Cervantes, A., Rajendran, K., et al. (2023). Induction of astrocytic Slc22a3 regulates sensory processing through histone serotonylation. Science 380, eade0027. 10.1126/science.ade0027.

21. Al-Kachak, A., Di Salvo, G., Fulton, S.L., Chan, J.C., Farrelly, L.A., Lepack, A.E., Bastle, R.M., Kong, L., Cathomas, F., Newman, E.L., et al. (2024). Histone serotonylation in dorsal raphe nucleus contributes to stress- and antidepressant-mediated gene expression and behavior. Nat Commun 15, 5042. 10.1038/s41467-024-49336-4.

22. Chan, J.C., Alenina, N., Cunningham, A.M., Ramakrishnan, A., Shen, L., Bader, M., and Maze, I. (2024). Serotonin Transporter-dependent Histone Serotonylation in Placenta Contributes to the Neurodevelopmental Transcriptome. J Mol Biol 436, 168454. 10.1016/j.jmb.2024.168454.

23. Chen, H.C., He, P., McDonald, M., Williamson, M.R., Varadharajan, S., Lozzi, B., Woo, J., Choi, D.J., Sardar, D., Huang-Hobbs, E., et al. (2024). Histone serotonylation regulates ependymoma tumorigenesis. Nature 632, 903–910. 10.1038/s41586-024-07751-z.

24. Zhang, N., Wu, J., Hossain, F., Peng, H., Li, H., Gibson, C., Chen, M., Zhang, H., Gao, S., Zheng, X., et al. (2024). Bioorthogonal Labeling and Enrichment of Histone Monoaminylation Reveal Its Accumulation and Regulatory Function in Cancer Cell Chromatin. J Am Chem Soc. 10.1021/jacs.4c04249.

25. Ling, T., Dai, Z., Wang, H., Kien, T.T., Cui, R., Yu, T., and Chen, J. (2024). Serotonylation in tumor-associated fibroblasts contributes to the tumor-promoting roles of serotonin in colorectal cancer. Cancer Lett 600, 217150. 10.1016/j.canlet.2024.217150.

26. Zhang, N., Wu, J., Gao, S., Peng, H., Li, H., Gibson, C., Wu, S., Zhu, J., and Zheng, Q. (2024). pH-Controlled Chemoselective Rapid Azo-Coupling Reaction (CRACR) Enables Global Profiling of Serotonylation Proteome in Cancer Cells. J Proteome Res 23, 4457–4466. 10.1021/acs.jproteome.4c00409.

27. Dong, R., Wang, T., Dong, W., Zhang, H., Li, Y., Tao, R., Liu, Q., Liang, H., Chen, X., Zhang, B., and Zhang, X. (2025). TGM2-mediated histone serotonylation promotes HCC progression via MYC signalling pathway. J Hepatol 83, 105–118. 10.1016/j.jhep.2024.12.038.

28. Liu, K., Zhang, Y., Du, G., Chen, X., Xiao, L., Jiang, L., Jing, N., Xu, P., Zhao, C., Liu, Y., et al. (2025). 5-HT orchestrates histone serotonylation and citrullination to drive neutrophil extracellular traps and liver metastasis. J Clin Invest 135. 10.1172/JCI183544.

29. Navarro-Corcuera, A., and Martinez-Chantar, M.L. (2025). Histone serotonylation in HCC: Decoding the impact of “happy” histones on liver cancer progression. J Hepatol 83, 18–20. 10.1016/j.jhep.2025.02.020.

30. Lin, S., Tan, S., Peng, Y., Tulamaiti, A., Du, W., Ding, K., Chen, C., Wu, J., Li, H., Xu, W., et al. (2025). Histone serotonylation promotes pancreatic cancer development via lipid metabolism remodeling. Nat Commun 16, 5947. 10.1038/s41467-025-61197-z.

31. Ji, Y., Ju, C.W., Chen, L., Shen, K., Su, R., Li, A., Liu, X., Liu, B., Zhang, X., Lyu, R., et al. (2025). Serotonin modulates lineage plasticity in neuroendocrine prostate cancer via epigenetic reprogramming. Cancer Discov. 10.1158/2159-8290.CD-25-0974.

32. Li, A., Xu, S., Fan, J., Liu, J., Song, X., Cao, L., Gong, Z., and Lu, X. (2026). 5-HT reuptake blockade induces pyroptosis in BRAF(V600E)-mutated melanomas via remodeling histone serotonylation. Cell Rep Med 7, 102537. 10.1016/j.xcrm.2025.102537.

33. Deller, S., Macheroux, P., and Sollner, S. (2008). Flavin-dependent quinone reductases. Cellular and Molecular Life Sciences 65, 141–160. 10.1007/s00018-007-7300-y.

34. Vella, F., Ferry, G., Delagrange, P., and Boutin, J.A. (2005). NRH:quinone reductase 2: an enzyme of surprises and mysteries. Biochem Pharmacol 71, 1–12. 10.1016/j.bcp.2005.09.019.

35. Islam, F., Leung, K.K., Walker, M.D., Al Massri, S., and Shilton, B.H. (2022). The Unusual Cosubstrate Specificity of NQO2: Conservation Throughout the Amniotes and Implications for Cellular Function. Frontiers in Pharmacology Volume 13 - 2022. 10.3389/fphar.2022.838500.

36. Islam, F., Basilone, N., Yoo, V., Ball, E., and Shilton, B. (2024). Evolutionary analysis of Quinone Reductases 1 and 2 suggests that NQO2 evolved to function as a pseudoenzyme. Protein Science 33, e5234. 10.1002/pro.5234.

37. Ferry, G., and Boutin, J.A. (2022). Measurement of NQO2 Catalytic Activity and of Its Inhibition by Melatonin. Methods Mol Biol 2550, 315–321. 10.1007/978-1-0716-2593-4_33.

38. Calamini, B., Santarsiero, B.D., Boutin, J.A., and Mesecar, A.D. (2008). Kinetic, thermodynamic and X-ray structural insights into the interaction of melatonin and analogues with quinone reductase 2. Biochem J 413, 81–91. 10.1042/bj20071373.

39. Calamini, B., Ferry, G., and Boutin, J.A. (2022). Cloning, Expression, Purification, Crystallization, and X-Ray Structural Determination of the Human NQO2 in Complex with Melatonin. Methods Mol Biol 2550, 291–304. 10.1007/978-1-0716-2593-4_31.

40. Gould, N.L., Scherer, G.R., Carvalho, S., Shurrush, K., Kayyal, H., Edry, E., Elkobi, A., David, O., Foqara, M., Thakar, D., et al. (2023). Specific quinone reductase 2 inhibitors reduce metabolic burden and reverse Alzheimer’s disease phenotype in mice. J Clin Invest 133. 10.1172/jci162120.

41. Brouillette, J., and Quirion, R. (2008). Transthyretin: a key gene involved in the maintenance of memory capacities during aging. Neurobiol Aging 29, 1721–1732. 10.1016/j.neurobiolaging.2007.04.007.

42. Gould, N.L., Elkobi, A., Edry, E., Daume, J., and Rosenblum, K. (2020). Muscarinic-Dependent miR-182 and QR2 Expression Regulation in the Anterior Insula Enables Novel Taste Learning. eNeuro 7. 10.1523/eneuro.0067-20.2020.

43. Hashimoto, T., and Nakai, M. (2011). Increased hippocampal quinone reductase 2 in Alzheimer’s disease. Neurosci Lett 502, 10–12. 10.1016/j.neulet.2011.07.008.

44. Rappaport, A.N., Jacob, E., Sharma, V., Inberg, S., Elkobi, A., Ounallah-Saad, H., Pasmanik-Chor, M., Edry, E., and Rosenblum, K. (2015). Expression of Quinone Reductase-2 in the Cortex Is a Muscarinic Acetylcholine Receptor-Dependent Memory Consolidation Constraint. J Neurosci 35, 15568–15581. 10.1523/jneurosci.1170-15.2015.

45. Janda, E., Boutin, J.A., De Lorenzo, C., and Arbitrio, M. (2024). Polymorphisms and Pharmacogenomics of NQO2: The Past and the Future. Genes (Basel) 15. 10.3390/genes15010087.

46. Benoit, C.E., Bastianetto, S., Brouillette, J., Tse, Y., Boutin, J.A., Delagrange, P., Wong, T., Sarret, P., and Quirion, R. (2010). Loss of quinone reductase 2 function selectively facilitates learning behaviors. J Neurosci 30, 12690–12700. 10.1523/JNEUROSCI.2808-10.2010.

47. Lin, J., Bao, X., and Li, X.D. (2021). A tri-functional amino acid enables mapping of binding sites for posttranslational-modification-mediated protein-protein interactions. Mol Cell 81, 2669–2681 e2669. 10.1016/j.molcel.2021.04.001.

48. Skene, P.J., and Henikoff, S. (2017). An efficient targeted nuclease strategy for high-resolution mapping of DNA binding sites. Elife 6. 10.7554/eLife.21856.

49. Maze, I., Wenderski, W., Noh, K.M., Bagot, R.C., Tzavaras, N., Purushothaman, I., Elsasser, S.J., Guo, Y., Ionete, C., Hurd, Y.L., et al. (2015). Critical Role of Histone Turnover in Neuronal Transcription and Plasticity. Neuron 87, 77–94. 10.1016/j.neuron.2015.06.014.

50. Zhang, Y., Pak, C., Han, Y., Ahlenius, H., Zhang, Z., Chanda, S., Marro, S., Patzke, C., Acuna, C., Covy, J., et al. (2013). Rapid single-step induction of functional neurons from human pluripotent stem cells. Neuron 78, 785–798. 10.1016/j.neuron.2013.05.029.

51. Kang, H.J., Kawasawa, Y.I., Cheng, F., Zhu, Y., Xu, X., Li, M., Sousa, A.M., Pletikos, M., Meyer, K.A., Sedmak, G., et al. (2011). Spatio-temporal transcriptome of the human brain. Nature 478, 483–489. 10.1038/nature10523.

52. Kreitzer, F.R., Salomonis, N., Sheehan, A., Huang, M., Park, J.S., Spindler, M.J., Lizarraga, P., Weiss, W.A., So, P.L., and Conklin, B.R. (2013). A robust method to derive functional neural crest cells from human pluripotent stem cells. Am J Stem Cells 2, 119–131.

53. Funk, O.H., Qalieh, Y., Doyle, D.Z., Lam, M.M., and Kwan, K.Y. (2022). Postmitotic accumulation of histone variant H3.3 in new cortical neurons establishes neuronal chromatin, transcriptome, and identity. Proc Natl Acad Sci U S A 119, e2116956119. 10.1073/pnas.2116956119.

54. Shen, Z., Kang, J., Shakya, A., Tabaka, M., Jarboe, E.A., Regev, A., and Tantin, D. (2017). Enforcement of developmental lineage specificity by transcription factor Oct1. Elife 6. 10.7554/eLife.20937.

55. Velasco, S., Kedaigle, A.J., Simmons, S.K., Nash, A., Rocha, M., Quadrato, G., Paulsen, B., Nguyen, L., Adiconis, X., Regev, A., et al. (2019). Individual brain organoids reproducibly form cell diversity of the human cerebral cortex. Nature 570, 523–527. 10.1038/s41586-019-1289-x.

56. Uzquiano, A., Kedaigle, A.J., Pigoni, M., Paulsen, B., Adiconis, X., Kim, K., Faits, T., Nagaraja, S., Antón-Bolaños, N., Gerhardinger, C., et al. (2022). Proper acquisition of cell class identity in organoids allows definition of fate specification programs of the human cerebral cortex. Cell 185, 3770–3788.e3727. 10.1016/j.cell.2022.09.010.

57. Gorski, J.A., Talley, T., Qiu, M., Puelles, L., Rubenstein, J.L., and Jones, K.R. (2002). Cortical excitatory neurons and glia, but not GABAergic neurons, are produced in the Emx1-expressing lineage. J Neurosci 22, 6309–6314. 10.1523/jneurosci.22-15-06309.2002.

58. Polioudakis, D., de la Torre-Ubieta, L., Langerman, J., Elkins, A.G., Shi, X., Stein, J.L., Vuong, C.K., Nichterwitz, S., Gevorgian, M., Opland, C.K., et al. (2019). A Single-Cell Transcriptomic Atlas of Human Neocortical Development during Mid-gestation. Neuron 103, 785–801.e788. 10.1016/j.neuron.2019.06.011.

59. Kang, M., Gulati, G.S., Brown, E.L., Qi, Z., Avagyan, S., Armenteros, J.J.A., Gleyzer, R., Zhang, W., Steen, C.B., D’Silva, J.P., et al. (2025). Improved reconstruction of single-cell developmental potential with CytoTRACE 2. Nature Methods 22, 2258–2263. 10.1038/s41592-025-02857-2.

60. Simon-Areces, J., Dopazo, A., Dettenhofer, M., Rodriguez-Tebar, A., Garcia-Segura, L.M., and Arevalo, M.A. (2011). Formin1 mediates the induction of dendritogenesis and synaptogenesis by neurogenin3 in mouse hippocampal neurons. PLoS One 6, e21825. 10.1371/journal.pone.0021825.

61. Forneris, F., Binda, C., Vanoni, M.A., Mattevi, A., and Battaglioli, E. (2005). Histone demethylation catalysed by LSD1 is a flavin-dependent oxidative process. FEBS Lett 579, 2203–2207. 10.1016/j.febslet.2005.03.015.

62. Shakya, A., Cooksey, R., Cox, J.E., Wang, V., McClain, D.A., and Tantin, D. (2009). Oct1 loss of function induces a coordinate metabolic shift that opposes tumorigenicity. Nat Cell Biol 11, 320–327. 10.1038/ncb1840.

63. Tantin, D., Schild-Poulter, C., Wang, V., Haché, R.J., and Sharp, P.A. (2005). The octamer binding transcription factor Oct-1 is a stress sensor. Cancer Res 65, 10750–10758. 10.1158/0008-5472.Can-05-2399.

64. Gould, N.L., Sharma, V., Hleihil, M., Kolatt Chandran, S., David, O., Edry, E., and Rosenblum, K. (2020). Dopamine-Dependent QR2 Pathway Activation in CA1 Interneurons Enhances Novel Memory Formation. J Neurosci 40, 8698–8714. 10.1523/JNEUROSCI.1243-20.2020.

65. Gould, N.L., Kolatt Chandran, S., Kayyal, H., Edry, E., and Rosenblum, K. (2021). Somatostatin Interneurons of the Insula Mediate QR2-Dependent Novel Taste Memory Enhancement. eNeuro 8. 10.1523/ENEURO.0152-21.2021.

66. Iskander, K., and Jaiswal, A.K. (2005). Quinone oxidoreductases in protection against myelogenous hyperplasia and benzene toxicity. Chem Biol Interact 153-154, 147–157. 10.1016/j.cbi.2005.03.019.

67. Long, D.J., 2nd, Iskander, K., Gaikwad, A., Arin, M., Roop, D.R., Knox, R., Barrios, R., and Jaiswal, A.K. (2002). Disruption of dihydronicotinamide riboside:quinone oxidoreductase 2 (NQO2) leads to myeloid hyperplasia of bone marrow and decreased sensitivity to menadione toxicity. J Biol Chem 277, 46131–46139. 10.1074/jbc.M208675200.

68. Celli, C.M., Tran, N., Knox, R., and Jaiswal, A.K. (2006). NRH:quinone oxidoreductase 2 (NQO2) catalyzes metabolic activation of quinones and anti-tumor drugs. Biochem Pharmacol 72, 366–376. 10.1016/j.bcp.2006.04.029.

69. Vagin, A., and Teplyakov, A. (2010). Molecular replacement with MOLREP. Acta Crystallogr D Biol Crystallogr 66, 22–25. 10.1107/S0907444909042589.

70. Emsley, P., and Cowtan, K. (2004). Coot: model-building tools for molecular graphics. Acta Crystallogr D Biol Crystallogr 60, 2126–2132. 10.1107/S0907444904019158.

71. Adams, P.D., Afonine, P.V., Bunkoczi, G., Chen, V.B., Davis, I.W., Echols, N., Headd, J.J., Hung, L.W., Kapral, G.J., Grosse-Kunstleve, R.W., et al. (2010). PHENIX: a comprehensive Python-based system for macromolecular structure solution. Acta Crystallogr D Biol Crystallogr 66, 213–221. 10.1107/S0907444909052925.

72. Flood, D.T., Hintzen, J.C.J., Bird, M.J., Cistrone, P.A., Chen, J.S., and Dawson, P.E. (2018). Leveraging the Knorr Pyrazole Synthesis for the Facile Generation of Thioester Surrogates for use in Native Chemical Ligation. Angew Chem Int Ed Engl 57, 11634–11639. 10.1002/anie.201805191.

73. Simon, M.D., Chu, F., Racki, L.R., de la Cruz, C.C., Burlingame, A.L., Panning, B., Narlikar, G.J., and Shokat, K.M. (2007). The site-specific installation of methyl-lysine analogs into recombinant histones. Cell 128, 1003–1012. 10.1016/j.cell.2006.12.041.

74. Tyanova, S., Temu, T., and Cox, J. (2016). The MaxQuant computational platform for mass spectrometry-based shotgun proteomics. Nat Protoc 11, 2301–2319. 10.1038/nprot.2016.136.

75. Bilimoria, P.M., and Bonni, A. (2008). Cultures of cerebellar granule neurons. CSH Protoc 2008, pdb.prot5107. 10.1101/pdb.prot5107.

76. Malik, A.N., Vierbuchen, T., Hemberg, M., Rubin, A.A., Ling, E., Couch, C.H., Stroud, H., Spiegel, I., Farh, K.K., Harmin, D.A., and Greenberg, M.E. (2014). Genome-wide identification and characterization of functional neuronal activity-dependent enhancers. Nat Neurosci 17, 1330–1339. 10.1038/nn.3808.

77. Bray, N.L., Pimentel, H., Melsted, P., and Pachter, L. (2016). Near-optimal probabilistic RNA-seq quantification. Nature Biotechnology 34, 525–527. 10.1038/nbt.3519.

78. Love, M.I., Huber, W., and Anders, S. (2014). Moderated estimation of fold change and dispersion for RNA-seq data with DESeq2. Genome Biol 15, 550. 10.1186/s13059-014-0550-8.

79. Cahill, K.M., Huo, Z., Tseng, G.C., Logan, R.W., and Seney, M.L. (2018). Improved identification of concordant and discordant gene expression signatures using an updated rank-rank hypergeometric overlap approach. Scientific Reports 8, 9588. 10.1038/s41598-018-27903-2.

80. Kuleshov, M.V., Jones, M.R., Rouillard, A.D., Fernandez, N.F., Duan, Q., Wang, Z., Koplev, S., Jenkins, S.L., Jagodnik, K.M., Lachmann, A., et al. (2016). Enrichr: a comprehensive gene set enrichment analysis web server 2016 update. Nucleic Acids Res 44, W90–97. 10.1093/nar/gkw377.

81. Chen, E.Y., Tan, C.M., Kou, Y., Duan, Q., Wang, Z., Meirelles, G.V., Clark, N.R., and Ma’ayan, A. (2013). Enrichr: interactive and collaborative HTML5 gene list enrichment analysis tool. BMC Bioinformatics 14, 128. 10.1186/1471-2105-14-128.

82. Hu, C., Li, T., Xu, Y., Zhang, X., Li, F., Bai, J., Chen, J., Jiang, W., Yang, K., Ou, Q., et al. (2023). CellMarker 2.0: an updated database of manually curated cell markers in human/mouse and web tools based on scRNA-seq data. Nucleic Acids Res 51, D870–d876. 10.1093/nar/gkac947.

83. Kim, D., Langmead, B., and Salzberg, S.L. (2015). HISAT: a fast spliced aligner with low memory requirements. Nature Methods 12, 357–360. 10.1038/nmeth.3317.

84. Zhang, Y., Liu, T., Meyer, C.A., Eeckhoute, J., Johnson, D.S., Bernstein, B.E., Nusbaum, C., Myers, R.M., Brown, M., Li, W., and Liu, X.S. (2008). Model-based Analysis of ChIP-Seq (MACS). Genome Biology 9, R137. 10.1186/gb-2008-9-9-r137.

85. Stark, R., and Brown, G. (2011). DiffBind: differential binding analysis of ChIP-Seq peak data. R package version 100, 2–21.

86. Yu, G., Wang, L.-G., and He, Q.-Y. (2015). ChIPseeker: an R/Bioconductor package for ChIP peak annotation, comparison and visualization. Bioinformatics 31, 2382–2383. 10.1093/bioinformatics/btv145.

87. Ramírez, F., Dündar, F., Diehl, S., Grüning, B.A., and Manke, T. (2014). deepTools: a flexible platform for exploring deep-sequencing data. Nucleic acids research 42, W187–W191.

88. Skene, P.J., Henikoff, J.G., and Henikoff, S. (2018). Targeted in situ genome-wide profiling with high efficiency for low cell numbers. Nat Protoc 13, 1006–1019. 10.1038/nprot.2018.015.

89. Meers, M.P., Bryson, T.D., Henikoff, J.G., and Henikoff, S. (2019). Improved CUT&RUN chromatin profiling tools. Elife 8. 10.7554/eLife.46314.

90. Langmead, B., and Salzberg, S.L. (2012). Fast gapped-read alignment with Bowtie 2. Nat Methods 9, 357–359. 10.1038/nmeth.1923.

91. Li, H., Handsaker, B., Wysoker, A., Fennell, T., Ruan, J., Homer, N., Marth, G., Abecasis, G., and Durbin, R. (2009). The Sequence Alignment/Map format and SAMtools. Bioinformatics 25, 2078–2079. 10.1093/bioinformatics/btp352.

92. Fisher, R.A. (1922). On the Interpretation of χ2 from Contingency Tables, and the Calculation of P. Journal of the Royal Statistical Society 85, 87–94. 10.2307/2340521.

93. Bentsen, M., Goymann, P., Schultheis, H., Klee, K., Petrova, A., Wiegandt, R., Fust, A., Preussner, J., Kuenne, C., Braun, T., et al. (2020). ATAC-seq footprinting unravels kinetics of transcription factor binding during zygotic genome activation. Nature Communications 11, 4267. 10.1038/s41467-020-18035-1.

94. Hao, Y., Stuart, T., Kowalski, M.H., Choudhary, S., Hoffman, P., Hartman, A., Srivastava, A., Molla, G., Madad, S., Fernandez-Granda, C., and Satija, R. (2024). Dictionary learning for integrative, multimodal and scalable single-cell analysis. Nature Biotechnology 42, 293–304. 10.1038/s41587-023-01767-y.

95. Stuart, T., Srivastava, A., Madad, S., Lareau, C.A., and Satija, R. (2021). Single-cell chromatin state analysis with Signac. Nat Methods 18, 1333–1341. 10.1038/s41592-021-01282-5.

96. Rainer, J., Gatto, L., and Weichenberger, C.X. (2019). ensembldb: an R package to create and use Ensembl-based annotation resources. Bioinformatics 35, 3151–3153. 10.1093/bioinformatics/btz031.

97. Yang, S., Corbett, S.E., Koga, Y., Wang, Z., Johnson, W.E., Yajima, M., and Campbell, J.D. (2020). Decontamination of ambient RNA in single-cell RNA-seq with DecontX. Genome Biol 21, 57. 10.1186/s13059-020-1950-6.

98. Germain, P.L., Lun, A., Garcia Meixide, C., Macnair, W., and Robinson, M.D. (2021). Doublet identification in single-cell sequencing data using scDblFinder. F1000Res 10, 979. 10.12688/f1000research.73600.2.

99. Choudhary, S., and Satija, R. (2022). Comparison and evaluation of statistical error models for scRNA-seq. Genome Biol 23, 27. 10.1186/s13059-021-02584-9.

100. Gulati, G.S., Sikandar, S.S., Wesche, D.J., Manjunath, A., Bharadwaj, A., Berger, M.J., Ilagan, F., Kuo, A.H., Hsieh, R.W., Cai, S., et al. (2020). Single-cell transcriptional diversity is a hallmark of developmental potential. Science 367, 405–411. 10.1126/science.aax0249.

101. Liberzon, A., Birger, C., Thorvaldsdóttir, H., Ghandi, M., Mesirov, J.P., and Tamayo, P. (2015). The Molecular Signatures Database (MSigDB) hallmark gene set collection. Cell Syst 1, 417–425. 10.1016/j.cels.2015.12.004.

102. Korotkevich, G., Sukhov, V., Budin, N., Shpak, B., Artyomov, M.N., and Sergushichev, A. (2021). Fast gene set enrichment analysis. bioRxiv, 060012. 10.1101/060012.

